# A multi-omic atlas in the African turquoise killifish reveals increased glucocorticoid signaling as a hallmark of brain aging

**DOI:** 10.64898/2026.04.09.717549

**Authors:** Rapheal G. Williams, Bryan B. Teefy, Aaron J.J. Lemus, Evelyn H. Lee, Rajyk Bhala, Minhoo Kim, Hengyu Zhou, Ari Adler, Atul Kashyap, George M. Cardenas, Steven A. McCarroll, John Tower, Bérénice A. Benayoun

**Affiliations:** Leonard Davis School of Gerontology, University of Southern California, Los Angeles, CA 90089, USA; Molecular and Cellular Biosciences, USC Dornsife College of Letters, Arts and Sciences, Los Angeles, CA 90089, USA; Department of Genetics, Harvard Medical School, Boston, MA 02115, USA; Howard Hughes Medical Institute, Boston, MA 02115, USA; Cancer Biology Department, USC Keck School of Medicine, Los Angeles, CA 90033, USA; Pharmacology and Pharmaceutical Sciences Department, USC Mann School of Pharmacy, Los Angeles, CA 90033, USA; USC Norris Comprehensive Cancer Center, Epigenetics and Gene Regulation, Los Angeles, CA 90089, USA; USC Stem Cell Initiative, Los Angeles, CA 90089, USA

## Abstract

Aging is the leading risk factor for cognitive impairment and neurodegeneration, yet molecular changes that unfold in the brain over time, and how they drive this vulnerability, remain unclear. The naturally short-lived African turquoise killifish (*Nothobranchius furzeri*) offers a powerful model to understand brain aging on an accelerated timescale and test the impact of potential interventions. Here, we present a multi-omic atlas of brain aging of female and male African turquoise killifish from 2 independent genetic strains of different captive lifespans, encompassing single-nuclei RNA-seq, single nuclei ATAC-seq, and bulk ATAC-seq to capture transcriptional and regulatory changes. Interestingly, our atlas indicates that aging leads to a significant expansion of microglia numbers, regardless of sex or strain, which we independently validate using *in-situ* hybridization. In addition, we identify robust and conserved gene regulation changes, that are consistent with activation of glucocorticoid signalling as a hallmark (and potential driver) of vertebrate brain aging. Furthermore, pharmacological inhibition of glucocorticoid receptor activity starting at middle-age led to significant rescue of key molecular and cellular aging phenotypes. Thus, our study provides a powerful resource and framework to leverage the African turquoise killifish and rapidly uncover actionable pathways driving brain aging.

## Introduction

Biological aging is a gradual and dynamic process marked by declining cellular homeostasis and resilience. Aging is also the greatest risk factor for a myriad of diseases, notably through systemic innate immune activation and defective response to pathogen exposure^1–5^. In the brain, aging can manifest as synaptic remodeling, altered proteostasis, increased inflammation, and subsequent glial activation^6^. Together, these hallmarks of brain aging lead to cognitive decline and increased neurodegeneration risk^6^.

The African turquoise killifish (*Nothobranchius furzeri*) has emerged as a uniquely tractable vertebrate model for aging research due to its naturally short lifespan (∼4-6 months) and spontaneous development of age-associated molecular and cellular hallmarks^7–12^. Importantly, unlike traditional long-lived vertebrate model organisms, the turquoise killifish enables longitudinal and interventional analyses of neuroinflammation, neurodegeneration, gliosis, proteostasis collapse, cellular senescence, and cognitive decline within experimentally tractable timeframes^13–18^. Valuable resources, including reference genomes^19,20^ and transgenesis protocols^21,22^, have been established in this model. In addition, the turquoise killifish is also amenable to pharmacological interventions to promote partial rescue of some age-related phenotypes^23,24^. Mechanistic, spatially resolved studies of brain aging in this species are now supported by emerging neuroanatomical cell– and region-specific atlases^25–27^.

In the context of brain aging, turquoise killifish exhibit alterations consistent with accumulation of senescent cells and protein aggregation, akin to phenotypes seen in patients with Parkinson’s, Alzheimer’s, and Huntington’s diseases^9,28,29^. Previous studies have examined the impact of aging on the turquoise killifish brain in a single sex and/or only longer-lived non-inbred strains^24,30–32^. These studies revealed important biology relating to loss of neural progenitors with aging, or a dramatic impact of proteostasis collapse and protein-gene “decoupling”^30,31^, although not generalizable across sex and/or genetic backgrounds. Still, little is known about how the genomic regulation landscape of the turquoise killifish brain changes during aging, across sex and genetic backgrounds.

Here, we present a longitudinal multi-omic atlas of brain aging across the lifespan of both female and male African turquoise killifish, in two independent strains of diverging lifespans (GRZ and ZMZ-1001). Using a combination of automated annotation strategies and manual curation, we identify 16 major cell types (*e.g*. neurons, astrocytes/radial glia, oligodendrocytes, microglia) in the turquoise killifish brain, consistent with expectations for a vertebrate brain. Our transcriptomic data reveals both a robust expansion of the microglia compartment in the aging brain, as well as a strong upregulation of inflammatory gene programs across cell types, regardless of sex and strain, which we validate using spatially resolved multiplex, RNA fluorescent *in-situ* hybridization (FISH). We also identify a signature of increased glucocorticoid receptor (*nr3c1*) activity across ‘omic’ layers, which is conserved in mouse and human datasets. Lastly, we show that we can ameliorate age-related molecular and cellular phenotypes by using mid-life-initiated treatment with the high-affinity, competitive antagonist of glucocorticoid receptors, mifepristone. By explicitly integrating sex and strain variables with cell-type resolved analyses, our atlas extends on prior turquoise killifish resources and provides a powerful, more generalizable platform to elucidate robust contributors to vertebrate brain aging.

## Results

### An atlas of brain aging in the naturally short-lived African turquoise killifish

We leveraged the African turquoise killifish, the shortest-lived vertebrate that can be bred in captivity, to better understand robust molecular features of vertebrate brain aging as a function of sex and life expectancy. Specifically, we performed multi-omic profiling of aging brain tissue in this species, using both the standard short-lived inbred lab strain GRZ^19,33^ and the longer-lived wild-derived ZMZ-1001^19,34^ strain (**Fig. 1a**). The ZMZ-1001 strain was derived from wild African turquoise killifish collected in southern Mozambique in Spring 2010^19^, and robustly displays a longer median captive lifespan compared to the standard highly inbred GRZ strain derived from the Gona-Re-Zhou National Park of Zimbabwe in 1970^19,34^ (∼20-26 weeks *vs.* ∼40 weeks; **Fig. 1a**). The ZMZ-1001 strain is of the same yellow tail morph as the GRZ strain, and it is genetically more similar to the GRZ strain than other commonly used long-lived strains (<10^6^ fixed single nucleotide polymorphisms compared to the GRZ reference genome; **Extended Data Fig. 1a-c**). These features facilitate direct comparisons of biological processes as a function of aging in shorter– and longer-lived individuals of the same species, allowing the exploration of whether longer lifespan may lead to a slowdown or delay of brain aging phenotypes.

**Figure 1:**
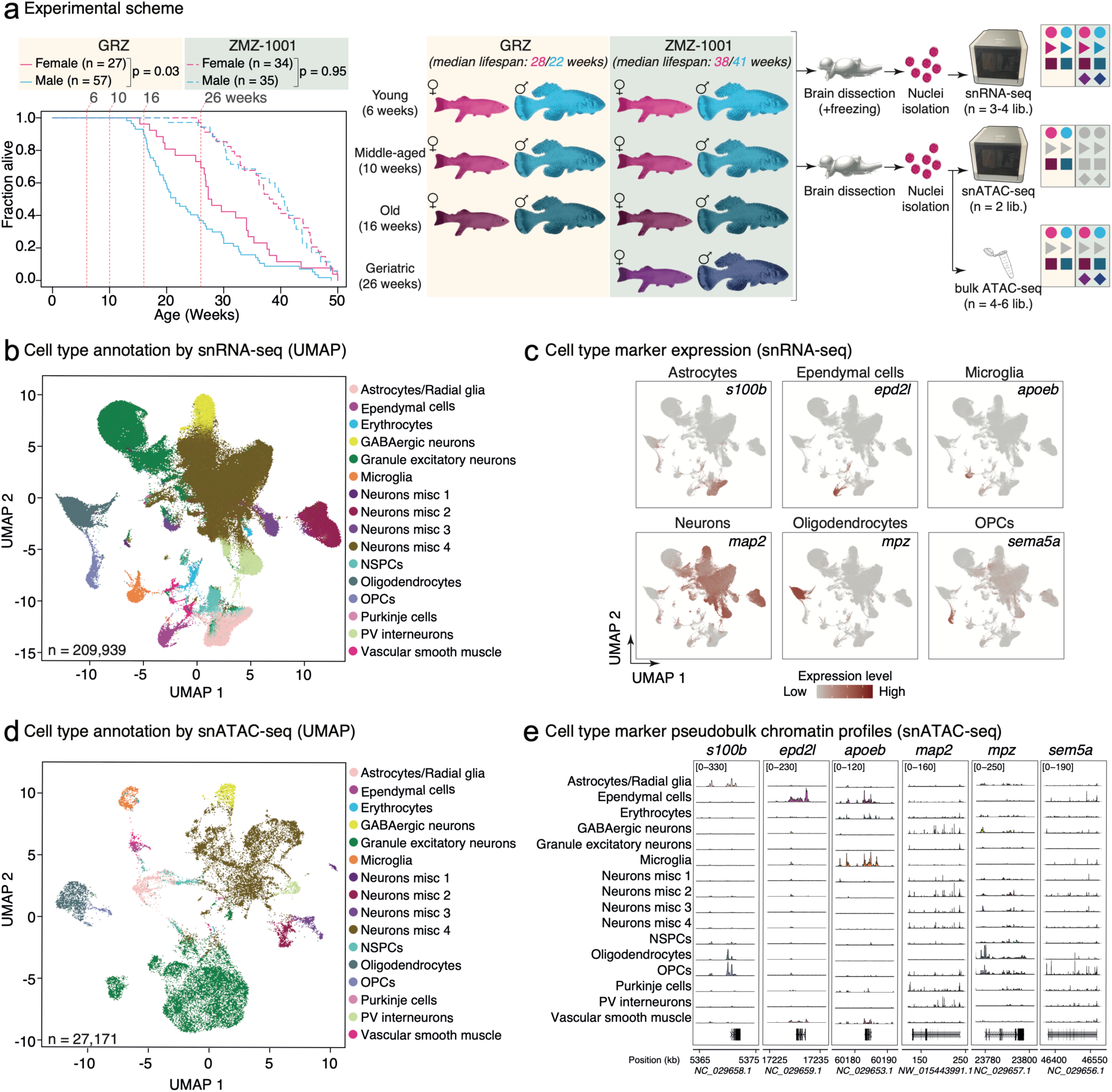
An atlas of brain aging in the naturally short-lived African turquoise killifish across sex and genetic strains. (**a**) Experimental scheme for the study. [Left] Survival analysis of fish from the GRZ and ZMZ-1001 strains at USC, showing the position of chosen ages on fish lifespan curves. Significance of longevity differences between sexes in each strain are reported using a Cox proportion hazard model. [Right] Experimental groups and workflow for generation of single-nuclei and bulk profiling of aging brain samples. (**b**) UMAP plot of our single nuclei RNA-seq dataset of turquoise killifish brain aging by annotated cell type. OPCs: Oligodendrocyte Progenitor Cells; NSPCs: Neural Stem and Progenitor Cells. (**c**) Expression UMAP plot for top cell type marker genes for astrocytes, ependymal cells, microglia, neurons, oligodendrocytes and oligodendrocyte progenitor cells [OPCs]. (**d**) UMAP plot of our single nuclei ATAC-seq dataset of turquoise killifish brain aging by annotated cell type. (**e**) Pseudobulk chromatin accessibility profiles of top cell type marker genes for astrocytes, ependymal cells, microglia, neurons, oligodendrocytes and oligodendrocyte progenitor cells [OPCs].

To map out the impact of aging on the brain epigenomic and transcriptomic landscapes, we generated (i) single nuclei RNA-seq (snRNA-seq) profiles using the 10x Genomics platform (GRZ and ZMZ-1001), (ii) single nuclei ATAC-seq (snATAC-seq) profiles using the 10x Genomics platform (GRZ only), and (iii) bulk ATAC-seq profiles using the Omni-ATAC protocol^35^ (GRZ and ZMZ-1001) from brain tissues throughout adulthood and aging (**Fig. 1a**). For GRZ fish, we defined young animals as ∼6 weeks old (sexually mature adults, generally at peak health), middle-aged animals as ∼10 weeks old, and old animals as ∼16 weeks old. Due to their longer lifespan, in addition to these 3 time points, we also included an additional geriatric time point for ZMZ-1001 fish, defined as ∼26 weeks old. We chose the 16 weeks old time point for the GRZ strain and the 26 weeks old time point for ZMZ-1001 fish as the oldest age to profile, as these time points correspond to the ∼90^th^ percentile of survival for each strain (**Fig. 1a**), to avoid known issues related to survivorship bias in experimental aging research^36^. For each assayed biological group, we generated 3-4 independent replicate snRNA-seq libraries, 2 independent replicate snATAC-seq libraries, and 4-6 independent replicate bulk ATAC-seq libraries, for a total of 44 snRNA-seq libraries, 8 snATAC-seq libraries and 48 bulk ATAC-seq libraries (**Fig. 1a**). To our knowledge, these datasets are the largest multi-omic atlas of brain aging in this species, which also encompasses both sexes and two independent strains with different captive lifespan.

After pre-processing and quality filtering, we obtained 209,939 high quality brain nuclei transcriptomes across 5 experimental batches, achieving a median sequencing depth of ∼39,441 reads per nucleus and a median sequence saturation rate of ∼77.3% across libraries (**Extended Data Table 1a**). Since the turquoise killifish genome is rich in transposable elements (TEs)^19,33,34,37,38^, we analyzed single-cell transcriptomes for both gene and TE-derived transcripts (see **Methods**). Importantly, we observed good mixing across UMAP space as a function of strain, sex, age and processing batch (**Extended Data Fig. 2a-d**). To annotate nuclei transcriptomes to their respective cell types, we use a semi-supervised annotation approach combining (i) unsupervised clustering, (ii) cross species marker-based annotation using mouse and zebrafish brain cell type markers^39^, (iii) cross species reference-based annotation methods using published mouse snRNA-seq brain atlases^40,41^, with remaining ambiguous clusters annotated after manual inspection (see **Methods**). With this approach, we identified 16 distinct brain cell types (8 neuronal and 8 non neuronal), encompassing expected vertebrate brain cell types with clear expression of canonical marker genes (**Fig. 1b-c; Extended Data Fig. 2e; Extended Data Table 1d**). Intriguingly, 4 defined populations of mature neurons exhibited features of glutamatergic neurons, but their identity remained partially ambiguous as top specifically expressed genes did not match known neuron subtypes, which we labeled Neurons misc 1-4 (**Fig. 1b**; **Extended Data Fig. 2e**; see **Methods**). For ease of use by the research community, we have made this dataset searchable with an interactive R shiny application (https://minhooki.shinyapps.io/killifish-brain-atlas/).

We next moved to process and annotate the snATAC-seq dataset, which focused on GRZ animals due to the need of freshly isolated brain tissue (see **Methods**; **Fig. 1a**). After pre-processing and quality filtering, we obtained 27,171 high quality brain nuclei epigenomes across 2 experimental batches, achieving a median sequencing depth of ∼32,913 fragments per nuclei and a median sequence saturation rate of ∼57.0% across libraries (**Extended Data Table 1b**). Similar to our snRNA-seq data, we observed good overall mixing across UMAP space as a function of sex, age, and processing batch (**Extended Data Fig. 3a-c**). For annotation purposes, promoter chromatin accessibility was used as a surrogate for gene expression activity, and our annotated snRNA-seq dataset was used as a reference atlas to assign cell identities to nuclei (see **Methods**). All 16 brain cell types from our snRNA-seq annotation were detected in the snATAC-seq dataset, with robust marker gene activity resolution (**Fig. 1d-e; Extended Data Fig. 3d; Extended Data Table 1d**).

Finally, we processed our bulk ATAC-seq datasets (derived from brain tissues of both GRZ and ZMZ-1001 fish), which yielded high-quality libraries at a median sequencing depth of 65,420,916 read pairs per library and a median alignment rate of ∼72.5% across libraries (see **Methods; Extended Data Table 1c**).

### Global features of the aging brain in the naturally short-lived African turquoise killifish

We queried our single nuclei transcriptomic and epigenomic resource to provide insights into the overall cellular composition of the African turquoise killifish brain, and how relative cell abundances may shift with aging (**Fig. 2a**). Unsurprisingly, neurons (specifically glutamatergic Neurons misc 4 and granule excitatory neurons) were the most abundant cell types in our snRNA-seq (**Fig. 2b**) and snATAC-seq (**Fig. 2c**) datasets. Although overall cell type abundances were consistent in the snRNA-seq datasets from GRZ and ZMZ-1001 strains, cell type abundances derived from our GRZ snATAC-seq dataset slightly differed from those derived from snRNA-seq (**Fig. 2b,c**). This discrepancy is likely the result of differential efficiency of nuclei isolation from frozen (snRNA-seq) and fresh (snATAC-seq) brain tissue (see **Methods**).

**Figure 2:**
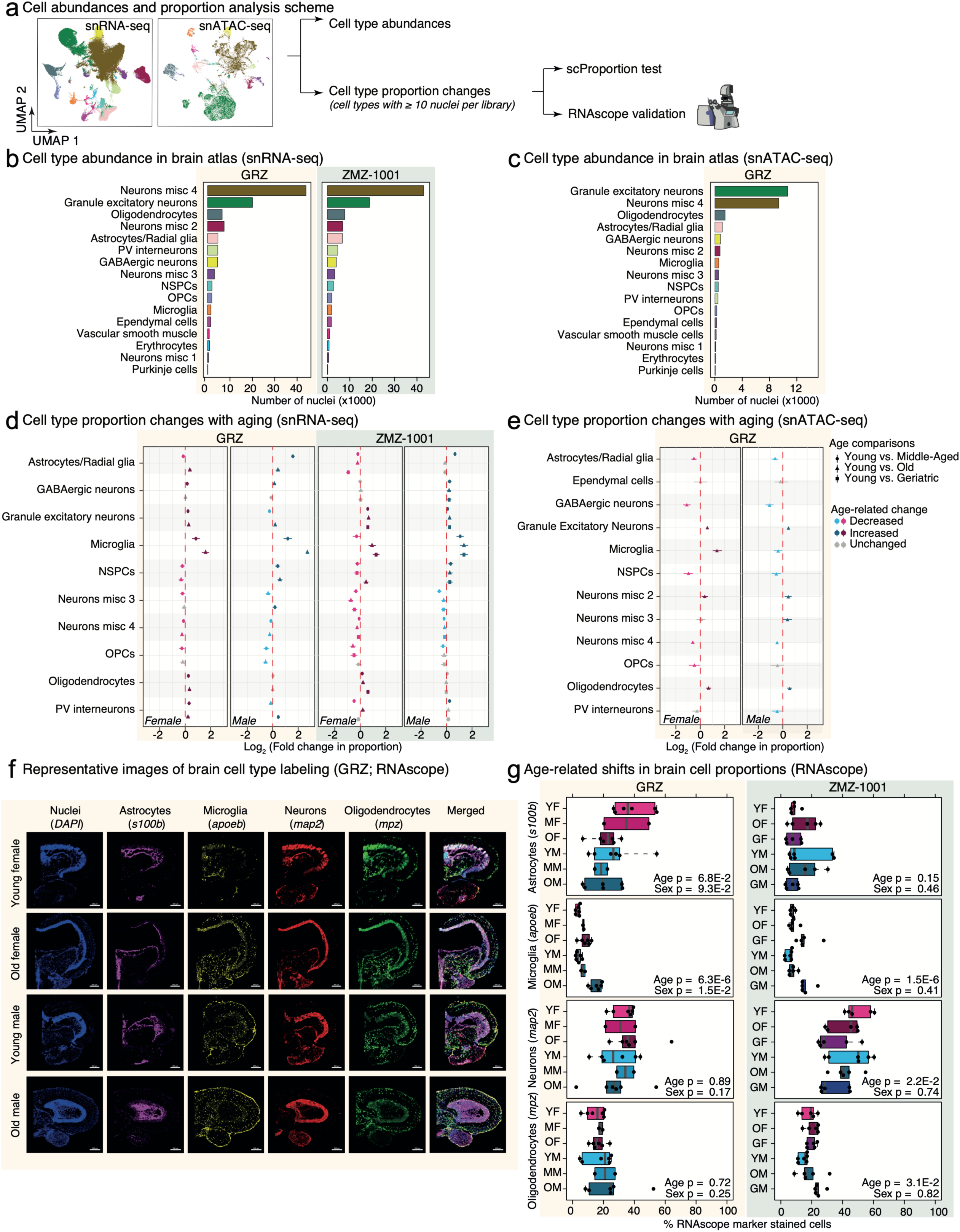
Cell type abundances and aging effects on cell proportions in the African turquoise killifish brain. (**a**) Scheme for cell type-level analysis of the aging African turquoise killifish brain. Analyses were performed using the annotated snRNA-seq and snATAC-seq datasets. Changes in cell type proportion with aging were validated using RNAscope *in-situ* hybridization. (**b-c**) Annotated cell type abundances in the snRNA-seq brain atlas for the GRZ and ZMZ-1001 strains (**b**) and in the snATAC-seq brain atlas for the GRZ strain (**c**). (**d-e**) Cell type proportion change analysis with aging across sexes based on our snRNA-seq (**d**) and snATAC-seq (**e**) datasets using scProportionTest. Left-shifted cell types are more abundant in young brains and right-shifted cell types are more abundant in old brains. Data is colored according to significance, with significant changes corresponding to FDR < 5% based on scProportionTest statistics. Note consistent increases in microglia proportion with aging across comparisons. (**f**) Representative maximum z-projected images using 4-plex RNAscope for each of the GRZ conditions’ cell-type markers: *apoeb* in yellow*, mpz* in green*, s100b* in magenta*, map2* in red and DAPI is in blue. See **Extended Data** Figure 5a for representative images from the ZMZ-1001 strain. Scale bars are 250mm in length. (**g**) Quantification of cell abundances in African turquoise killifish brain samples using top markers for *in-situ* hybridization by RNAscope. Values for each animal are reported as the average QuPath inferred values obtained across 3 Z-planes. Note that for this sample set, the middle-aged time point for GRZ corresponds to 13-week-old (rather than 10-week-old) fish. The effect of age and sex were evaluated using ANOVA for normally distributed data (GRZ astrocytes, microglia and neurons; ZMZ-1001 astrocytes, neurons, and oligodendrocytes) and non-parametric ART-ANOVA for non-normally distributed data (GRZ oligodendrocytes, ZMZ-1001 microglia). Normality of ANOVA residuals was determined using a Shapiro-Wilkes test, with p < 0.05 indicating violation of the normality assumption. Y: Young (6-7-week-old), M: middle-aged (13-week-old), O: Old (16-week-old), G: Geriatric (26-week-old). F: Female, M: Male.

To identify which cell type(s) were transcriptionally and epigenetically most responsive to aging, we applied the Augur^42,43^ cell prioritization algorithm on our snRNA-seq and snATAC-seq datasets (**Extended Data Fig. 4a**). Surprisingly, our Augur analysis revealed that cell types with the most consistent strong response to aging across sex, strains and ‘omic’ layers were oligodendrocyte progenitor cells [OPCs], oligodendrocytes, microglia, and neural stem and progenitor cells [NSPCs], rather than neurons themselves (**Extended Data Fig. 4a**). Thus, these analyses suggest that glial cells seem to be the cell type category most robustly impacted by aging, regardless of sex, genetic strain, and ‘omic’ regulatory layer.

Next, we used the single-cell scProportionTest^44^ tool to investigate whether there were any robust differences in relative cell proportions in the aging brain across sex and genetic strain, using our snRNA-seq and snATAC-seq datasets (**Fig. 2d,e**). The scProportionTest tool uses a permutation test to evaluate the magnitude and significance of shifts in cell type proportions^44^. Interestingly, many changes in brain cell type proportions with aging were detected in a sex-specific manner (*e.g*. NSPCs decreasing or oligodendrocytes increasing predominantly with brain aging in females; **Fig. 2d,e**) or omic-specific manner (*i.e*. robust decrease in astrocyte/radial glia proportion in snATAC-seq but not snRNA-seq dataset; **Fig. 2d,e**). Strikingly, we detected robust changes in cell proportions with aging across sex, strains and ‘omic’ datasets, including an increased proportion of microglia, a decreased proportion of the neuron misc 4 population, and an increased proportion of granule excitatory neurons (**Fig. 2d,e**).

To determine whether observed cell type proportion changes also occurred in intact, non-dissociated African turquoise killifish brain tissue, we performed RNAscope^45^ fluorescent RNA *in-situ* hybridization (RNA-ISH), using markers for astrocytes/radial glia (*s100b*), microglia (*apoeb*), neurons (*map2*) and oligodendrocytes (*mpz*), together with a DAPI stain to estimate overall cell numbers (**Fig. 2f,g**; **Extended Data Fig. 5a**). After watershed segmentation of cells based on DAPI signal in each confocal image using QuPath^46^, we classified each cell to their most likely cell type using an artificial neural network (ANN) to assign cell identity based on marker gene expression represented by fluorescence intensity (see **Methods**). Interestingly, we were able to validate microglia expansion in the aged turquoise killifish brain, regardless of sex or strain (**Fig. 2g**). To note, the scale of microglia expansion showed sex-dimorphism in the GRZ strain, with stronger increases in male fish, while no effect of sex was observed in the ZMZ-1001 strain (**Fig. 2g**). This observation may reflect the fact that female GRZ fish are longer lived than males in our facility (and may thus be biologically “younger” at the same chronological age), while no sex differences in the lifespan for ZMZ-1001 fish are observed (**Fig. 1a**). Other compositional changes were not consistently observed across sexes and strains (**Fig. 2g**), suggesting that they are not robust features of brain aging in this species.

### Multi-omic analysis of genomic regulation reveals widespread cell-specific genomic remodeling with aging

Next, we evaluated differential gene expression and chromatin accessibility profiles as a response to aging across our multi-omic atlas (**Fig. 3; Extended Data Fig. 6-9; Extended Data Table 2,3,4**). Intriguingly, and in contrast to our observations in peripheral cell types from the African turquoise killifish^34^, sex only had a relatively minor influence on the overall gene expression or chromatin accessibility explained variance in our snRNA-seq (**Fig. 3a-b**), snATAC-seq (**Extended Data Fig. 8a-b**), and bulk ATAC-seq (**Extended Data Fig. 9a-b**) datasets. These observations suggest that the brain is relatively insulated from sex-specific influences in this species, potentially contrasting with observations in mammals^47–50^. However, the relatively modest contribution of biological sex to brain ‘omic’ profiles provided us with increased statistical power to identify age-related changes by including sex as a modelling covariate in our DESeq2-based differential analyses (see **Methods**). Importantly, since single-cell level differential analyses suffer from high false discovery rates^51,52^, we used muscat^52^ to obtain pseudobulk profiles for each cell type across independent libraries and perform differential gene expression and chromatin accessibility analyses in our snRNA-seq and snATAC-seq datasets (**Fig. 3a; Extended Data Fig. 8a, 9a**), which enables use of conventional bulk computational methods. To limit the impact of technical noise and improve our ability to interpret results, only non-ambiguously annotated cell types with at least 10 nuclei in each snRNA-seq or snATAC-seq library were further analyzed, yielding 8 cell types for snRNA-seq analysis (astrocytes/radial glia, GABAergic neurons, granule excitatory neurons, microglia, NSPCs, oligodendrocytes, OPCs and PV interneurons) and 9 cell types for snATAC-seq analysis (same as snRNA-seq, with the addition of ependymal cells; see **Methods**).

**Figure 3:**
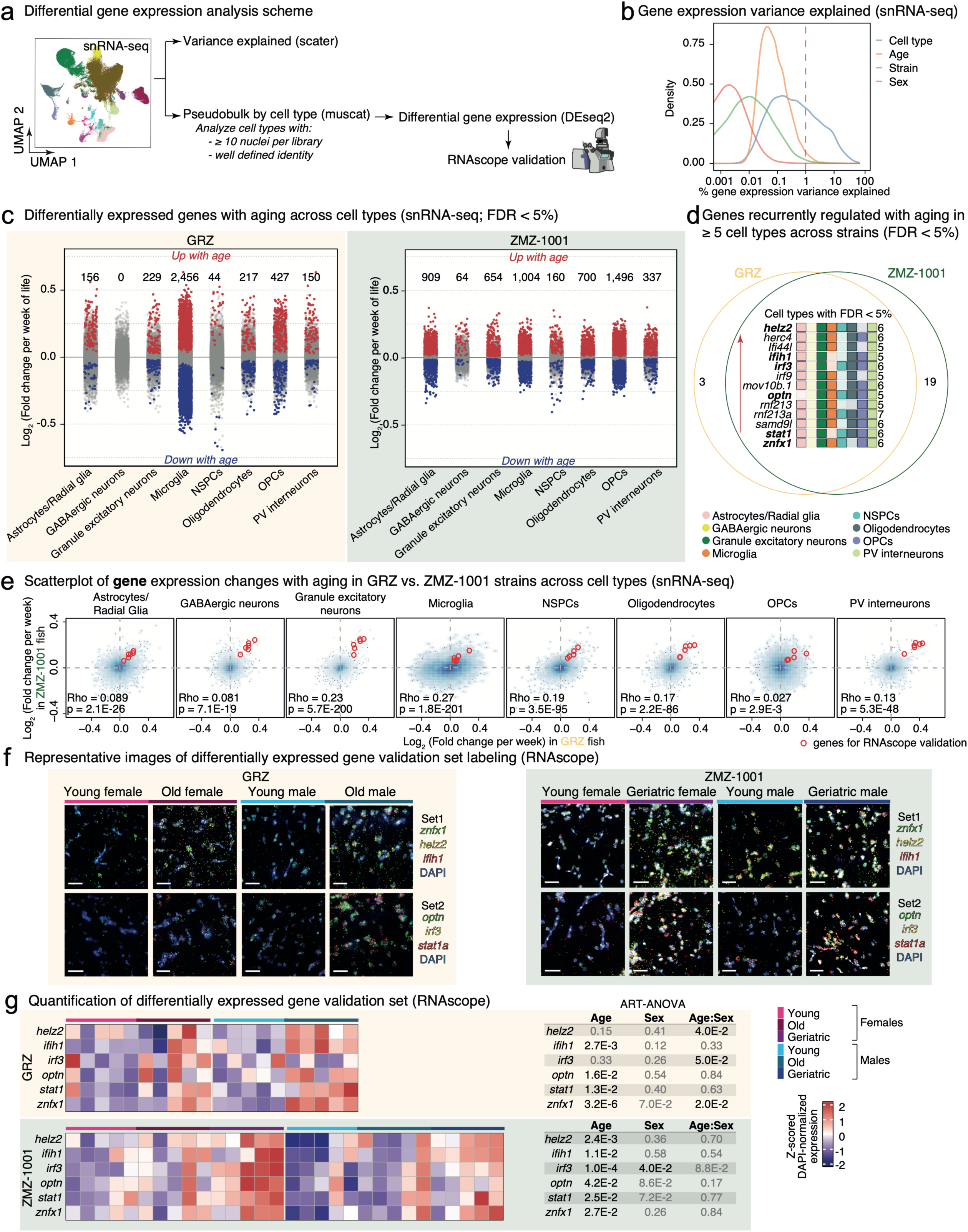
Differential gene expression analysis reveals cell-specific remodeling of gene expression with aging in the African turquoise killifish brain. **(a)** Scheme of differential gene expression analysis. Explained gene expression variance was estimated using scater. Cell types were partitioned in each strain according to their annotation in our snRNA-seq atlas and analyzed for differential regulation using pseudobulking with muscat. Differential gene expression analysis was performed with DESeq2. To maximize robustness and interpretability of pseudobulk analyses, only cell types with (i) ≥ 10 nuclei per snRNA-seq library and (ii) well-defined cell identity were analyzed. Top differentially regulated genes across cell types were validated using RNAscope *in-situ* hybridization. (**b**) Density plot of explained gene expression variance in snRNA-seq dataset as a function of cell type, age, strain, and sex. Notice that aging is the largest contributor to gene expression variance after cell type. (**c**) Strip plots of differentially expressed genes per cell type (FDR < 5%) in aging GRZ (left) and ZMZ-1001 (right) brains (sex was used as a covariate). Differentially expressed transcripts are highlighted while non-significant genes are shown in gray. Transcripts above the midline are upregulated with aging while transcripts below the midline downregulated with aging. (**d**) Analysis of recurrently upregulated genes across cell types with aging. Genes upregulated (as indicated by the red up-pointing arrow) at FDR < 5% with aging in ≥5 cell types were compared in the GRZ vs. ZMZ-1001 strains, with 13 genes shared. Bolded genes were selected for downstream validation based on a literature search. See **Extended Data Table 1d** for NCBI gene name correspondence. (**e**) Scatterplot of gene expression log_2_(Fold changes) with aging in GRZ vs. ZMZ-1001 cells across robust cell types. Inset are the Spearman Rho values and their significance. Top differentially expressed genes shared across cell types selected for RNAscope validation from panel (**d**) are highlighted in red. (**f**) Representative maximum z-projected images of the top differentially expressed gene set validations across age using two different 3-plex RNAscope assays for spatial validation in both the GRZ (left) and ZMZ-1001 (right) strains. Each age (young vs. old/geriatric) and sex (female vs. male) is represented for both sets. All channels (green, yellow, and red) for mRNA targets of interest of both Set 1 (top rows; *znfx1, helz2, ifih1*) and Set 2 (bottom rows; *optn, irf3, stat1a*) are merged for each image with DAPI. See **Extended Data** Figure 7a**,b** for split-channel representation. Scale bars are 30mm in length. (**g**) Quantificaton of differentially expressed gene validation set using *in-situ* hybridization by RNAscope. The number of labeled RNA spots was normalized to DAPI to account for cell numbers in any given brain slice. Values for each animal are reported as the average QuPath inferred quantification of the ratio of puncta to cells obtained across 3 Z-planes. Data was derived from n = 5 independent animals per biological group. The effects of age, sex and potential age:sex interactions were evaluated using non-parametric ART-ANOVA. Non-significant p-values are greyed out for ease of visualization.

Interestingly, the scale of age-related changes in gene and TE expression varied greatly across cell types, with some cell types showing few changes in expression (*i.e*. GABAergic neurons, NSPCs) and others showing abundant changes (*i.e*. microglia, OPCs) (**Fig. 3c; Extended Data Fig. 6a**). To note, cell-type-specific age-related changes in expression showed significant overlap and correlation in GRZ vs. ZMZ-1001 fish (**Fig. 3c; Extended Data Fig. 6a-d**), supporting the notion that aging leads to consistent changes in brain gene expression regardless of sex and strain, at least in this species. To note, the magnitude of gene expression changes, in terms of change per week of life, was larger in GRZ vs. ZMZ-1001 fish (**Fig. 3c; Extended Data Fig. 6a**), consistent with the notion that aging is slowed in the longer lived ZMZ-1001 strain compared to the shorter-lived GRZ strain, at least in the brain.

We next asked whether there were genes that were consistently differentially expressed with aging across cell types and strains (**Fig. 3d; Extended Data Fig. 6b**). Although we could not identify commonly downregulated genes with aging, we identified 13 genes showing upregulation in 5 out of 8 cell types in both GRZ and ZMZ-1001 fish with aging, many of which have been associated to inflammatory responses (**Fig. 3d; Extended Data Fig. 6b**). We then selected 6 of these genes for validation in intact brain tissue using RNAscope based on the current knowledge that supports their role in brain inflammation and disease progression: *helz2*^53,54^, *ifih1*^55,56^, *irf3*^57,58^, *optn*^59^, *stat1a*^60,61^ and *znfx1*^62,63^ (**Fig. 3d-e; Extended Data Fig. 6d**). Even in cell types where they did not pass the FDR significance threshold, these genes showed trends of upregulation with aging across sexes and strains (**Fig. 3e; Extended Data Fig. 6d**), suggesting that they represent good candidates for validation in a cell type-agnostic manner. Using QuPath, we then quantified the puncta representing each mRNA of interest by using the ‘subcellular detection’ tool after performing cell segmentation. We then counted the puncta within the soma using the DAPI signal in 3 distinct z-planes of each image and recorded the ratio of the number of ‘spots’ to the number of cells detected (**Fig. 3f,g; Extended Data Fig. 7a,b**). Based on our spot quantification, we confirmed a significant effect of aging on RNAscope expression of 4/6 and 6/6 of our candidate genes in GRZ and ZMZ-1001 brains, respectively (non-parametric ART-ANOVA; **Fig. 3f,g**). Interestingly, for *helz2* and *irf3*, which did not reach significance with aging in RNAscope staining of GRZ brains, a significant interaction between age and sex was detected, consistent with a potentially sex-specific aging effect on their expression (*i.e*. *helz2*: in males; *irf3*: in females; **Fig. 3g**). Together, our RNAscope quantification validates robustly upregulated candidate genes in the aging brain of the African turquoise killifish.

We next analyzed differential chromatin accessibility changes with aging in our snATAC-seq data from GRZ fish (**Extended Data Fig. 8c**). Interestingly, cell types that were most impacted by aging transcriptionally were not necessarily most impacted epigenetically (**Fig. 3c; Extended Data Fig. 8c**). For instance, granule excitatory neurons showed the most age-related changes in chromatin accessibility across cell types by an order of magnitude (13,527 differentially accessible peaks with aging, compared to 1,298 peaks for the next most impacted cell type, oligodendrocytes; **Extended Data Fig. 8c**), although they showed only modest changes at the transcriptional level (**Fig. 3c**). A recent study in aging mouse brain observed increased chromatin accessibility at heterochromatin domains in aged excitatory neurons^64^. Thus, our observations on chromatin accessibility with aging in granule excitatory neurons may reflect conservation of this feature in African turquoise killifish, although future studies mapping out heterochromatin domains would be needed to validate this finding. Finally, at the bulk ATAC-seq level, we observed strong impact of aging on chromatin accessibility landscapes, regardless of sex and strain (**Extended Data Fig. 9c,d**). Regions of differential chromatin accessibility with aging were found across genomic regions (**Extended Data Fig. 9e,f**). Regions of increased accessibility with aging tended to be more enriched for proximal regions (promoters/genes), while those of decreased accessibility with aging were more likely to overlap distal intergenic regions (likely enhancers; **Extended Data Fig. 9e,f**).

### Functional enrichment analyses in the aging African turquoise killifish brain reveal conserved features of vertebrate brain aging

We then set out to determine (i) whether transcriptional changes observed in mouse and human brain aging were conserved in the African turquoise killifish, and (ii) which functional processes were regulated in the aging African turquoise killifish brain across cell types (**Fig. 4a**; **Extended Data Fig. 10a**). First, we leveraged UCell^65^ to compute gene set activity scores for (i) robust human brain aging genes from GTEx^66^, (ii) a previously reported common aging gene [CAG] signature^50^ of upregulated genes in aged mouse brains, and (iii) the SenMayo signature of cell senescence^67^ (**Fig. 4b**). Genes upregulated in mouse and human aging brain showed consistent trends for upregulation with aging across brain cell types in both GRZ and ZMZ-1001 African turquoise killifish (**Fig. 4b**). In contrast, GTEx downregulated genes did not show consistent downregulation in our dataset (**Fig. 4b**). Intriguingly, we did not detect robust age-related gene changes of the SenMayo senescence signature across cell types (with a notable exception of the microglia; **Fig. 4b**), which may reflect the complexity of defining senescent cells in the brain^68^.

**Figure 4:**
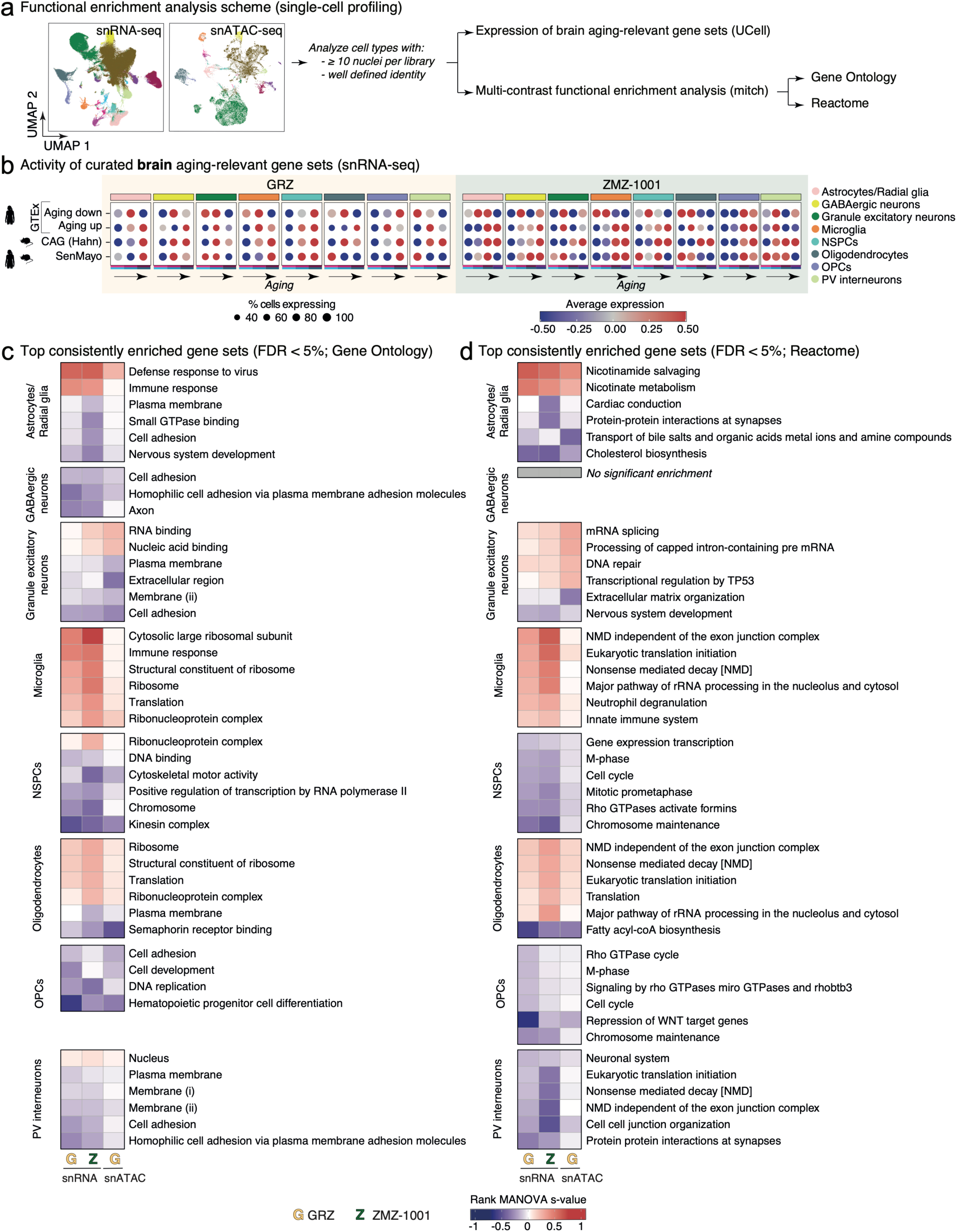
Functional enrichment analyses of differential gene expression in the aging African turquoise killifish brain. (**a**) Scheme of the functional enrichment analysis pipeline. Single-cell level gene set activity analysis with UCell was used to evaluate expression of curated brain aging-relevant gene sets across cell types with aging. Multi-contrast pathway enrichment was performed using mitch to identify genesets consistently regulated with aging in GRZ and ZMZ-1001 snRNA-seq, and GRZ snATAC-seq datasets in each cell type using Gene Ontology and Reactome gene set definitions. (**b**) UCell gene set activity for genes homologous to (i) genes regulated across brain regions with human aging in the GTEx dataset^66^, (ii) common aging genes [CAG] identified across mouse brain regions with aging^50^, and (ii) genes associated with the SenMayo signature^67^. Activity was calculated by age groups and reported separately in GRZ and ZMZ-1001 dataset in robust cell types. (**c-d**) Top Gene Ontology (c) and Reactome (d) gene sets regulated consistently with brain aging across robust cell types. The top 6 most enriched gene sets are represented, all with FDR < 5% (or however many reached significance). Heatmaps are colored based on the s statistic of a rank MANOVA test performed with mitch. Due to disambiguate GO naming redundancy: membrane (i) refers to GO:0016020, and membrane (ii) refers to GO:0016021.

Next, to better understand which functional processes may be impacted by aging, we leveraged multi-contrast pathway enrichment using mitch^69^ together with Gene Ontology (GO) and Reactome gene sets (**Fig. 4c,d**; **Extended Data Fig. 10c,d**; **Extended Data Table 5,6**). We observed regulation of a number of pathways related to hallmarks of aging^1,2,70^, including upregulation of inflammation-related pathways (*e.g*. GO “Immune response” in astrocytes/radial glia and microglia and Reactome “Innate immune system” in microglia and whole brain), NAD metabolism-related pathways (*e.g*. Reactome “Nicotinamide salvaging” in astrocytes/radial glia), DNA-damage/genomic instability-related pathways (*e.g*. Reactome “DNA repair” in granule excitatory neurons), protein homeostasis-related pathways (*e.g*. GO “Ribosome” and GO and Reactome “Translation” in microglia and oligodendrocytes), and extracellular matrix (ECM) changes (*e.g*. Reactome “Extracellular matrix organization” in granule excitatory neurons and whole brain; GO “Cell adhesion” in astrocytes/radial glia, GABAergic neurons, granule excitatory neurons, OPCs, PV interneurons and whole brain) (**Fig. 4c,d; Extended Data Fig. 10c,d; Extended Data Table 5,6**). More surprisingly, we observed misregulation of RNA-processing pathways (*e.g*. Reactome “Nonsense mediated decay [NMD]” in microglia, oligodendrocytes and PV interneurons; Reactome “Metabolism of RNA” and “RNA splicing” in whole brain) and of membrane-related pathways (e.g. GO “Membrane” in granule excitatory neurons, PV interneurons and whole brain; GO “Plasma membrane” in astrocytes/radial glia, granule excitatory neurons, oligodendrocytes, and PV interneurons), suggesting these might represent new less established hallmarks of aging (**Fig. 4c,d; Extended Data Fig. 10c,d; Extended Data Table 5,6**). Finally, we observed downregulation of cell proliferation-related pathways in NSPCs and OPCs (*e.g*. Reactome “Cell cycle” and “M-phase”; **Fig. 4c,d; Extended Data Table 5,6**), consistent with the notion of neural stem cells entering a dysfunctional state of deeper quiescence with aging^71^.

### Microglia undergo profound and conserved phenotypic remodeling with aging

Our previous analyses of our dataset found that microglia were among the most impacted cell types in the aging African turquoise killifish brain, including an age-related expansion (**Fig. 2d-g**), the largest consistent changes in gene expression (**Fig. 3c**), and strong signatures of inflammation/activation with aging (**Fig. 4c,d**). Microglia are the resident macrophage population of the brain, which perform critical roles in the brain throughout life (*e.g*. synapse pruning, cytokine and chemokine production, antigen presentation, phagocytosis of debris and protein aggregates), and whose dysfunction is thought to promote neuroinflammation in aging and neurodegenerative diseases^72–74^. Thus, we decided to analyze age-related changes in the microglia of African turquoise killifish to (i) characterize conservation of age-related transcriptional changes, (ii) determine potential shifts in activation and polarization phenotypes, (iii) validate increased microglia activation with aging by microscopy 3D reconstruction, and (iv) evaluate kinetics of microglia transcriptional aging in the GRZ and ZMZ-1001 strains (**Fig. 5a**).

**Figure 5:**
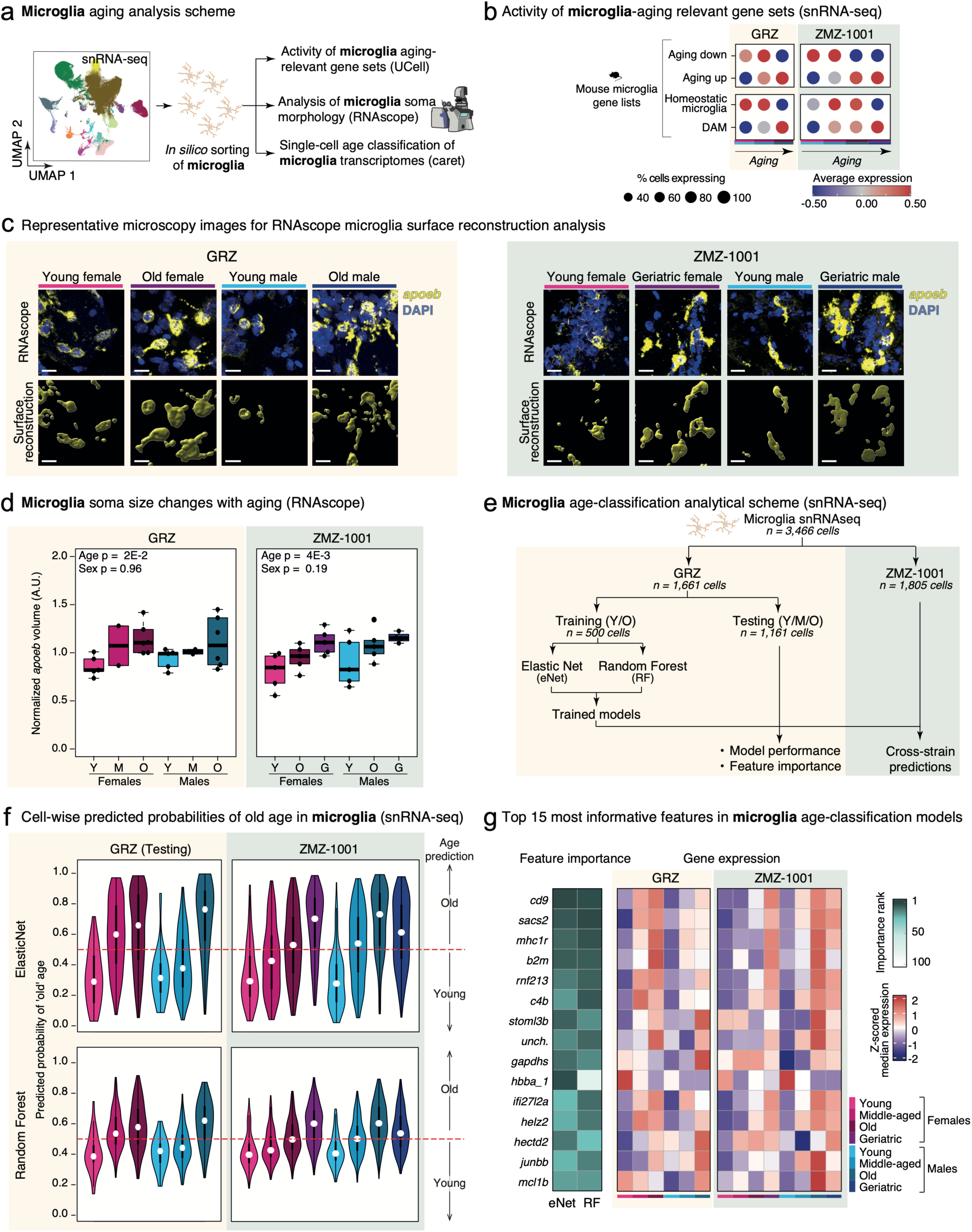
Analysis of microglia in the aging African turquoise killifish brain. (**a**) Scheme of a microglia-focused analysis in the aging brain. Microglia were sorted *in silico* for downstream analyses, including validation of top microglia-specific age regulated genes using RNAscope, evaluation of microglia aging-relevant gene set activity using UCell and machine-learning analysis of microglia. (**b**) UCell gene set activity for genes homologous to mouse (i) microglia aging regulated genes^75–77^, and (ii) signatures of homeostatic and damage-associated microglia [DAM]^75^. (**c**) Representative maximum z-projected images of microglia (*apoeb* signal;yellow) overlaid with DAPI (blue) in both GRZ (left) and ZMZ-1001 (right). Each age (young vs. old/geriatric) and sex (male vs. female) are represented. Using Imaris, 3D ‘surface’ reconstructions of the microglia are represented (bottom row). All scale bars are 15mm in length. (**d**) Relative microglia soma size with aging in African turquoise killifish brain samples derived from *apoeb* RNAscope 3D surface reconstruction (reanalysis of Fig. 2g images). Values for each animal are reported as the median volume of reconstructed microglia in the sample, normalized to the median value across each experimental batch of the cognate strain. Note that for this sample set, the middle-aged time point for GRZ corresponds to 13-week-old (rather than 10-week-old) fish. One significant outlier among GRZ young female samples, one significant outlier among GRZ young male samples and one significant outlier among ZMZ-1001 geriatric male samples were identified using the Grubbs test, which are omitted from the plot and statistical analysis. The effect of age and sex were evaluated using ANOVA. Normality of ANOVA residuals was determined using a Shapiro-Wilkes test (p > 0.05 indicating failure to reject the normality assumption). Y: Young (6-7-week-old), M: middle-aged (13-week-old), O: Old (16-week-old), G: Geriatric (26-week-old). F: Female, M: Male. (**e**) Scheme of machine-learning model training and performance evaluation for microglia aging, data was split by strain. For training, we randomly sampled 250 microglia transcriptomes from young and old GRZ animals. Remaining microglia (including those from middle-aged animals) were withheld for downstream model testing. After training, model performance was validated (see **Extended Data** Figure 12a-c), and analyses of predictions on withheld GRZ data and on ZMZ-1001 data was performed. (**f**) Violin plots of cell-wise predicted probabilities of belonging to an old brain using withheld microglia from GRZ snRNA-seq data and ZMZ-1001 snRNA-seq data, using our ElasticNet (top; eNET) or Random Forest (bottom; RF) models. (**g**) Heatmap of feature importance (*i.e*. mean decrease in Gini and absolute regularized coefficient), as well as Z-scored median expression for the top 15 features contributing to the eNET and RF models for microglia aging.

As suggested by our systematic analyses, we found that microglia showed strong separation based on ‘omic’ profiles with aging, with negligible impact of sex, at both the transcriptional and chromatin levels, according to multi-dimensional scaling (MDS) analysis (**Extended Data Fig. 11a**). We then asked how similar killifish microglia aging was to mammalian microglia aging, by using UCell scoring of mouse microglia age-regulated genes^75–77^ (**Fig. 5b**, upper panel). Consistent with a conserved response to aging in vertebrate microglia, we observed clear trend in aging turquoise killifish microglia for upregulation of genes that are upregulated with age in mouse microglia, and downregulation of genes downregulated with age in mouse microglia (**Fig. 5b**, upper panel). In addition, we also observed downregulation of markers of homeostatic microglia^75^, a concomitant upregulation of markers of a panoply of microglia activation signatures (*i.e*. damage-associated microglia [DAM]^75^, interferon-response microglia [IRM]^75^, and lipid-droplet-accumulating microglia [LDAM]^75^, terminally inflammatory microglia [TIM]^78^), as well as increased expression of M1/M2 polarization markers^79,80^ (**Fig. 5b**, lower panel; **Extended Data Fig. 11b**; see **Methods**). Thus, similar to mammalian microglia, turquoise killifish microglia seem to acquire a more activated phenotype with aging, irrespective of sex and strain.

Importantly, aged/activated microglia display increased soma volume in mouse neocortex^81^, acquiring a so-called “reactive” phenotype. We leveraged our *apoeb* RNAscope labeling data (**Fig. 2f,g**) to reconstruct microglial surfaces within the whole image and determine whether the microglia showed any morphometric changes during aging using the Imaris ‘surface’ creation tool (see **Methods**; **Fig. 5c,d**). As expected from observed transcriptional microglia activation signatures, we found that *apoeb*-labeled microglial soma volumes increased significantly with aging across sex and genetic strains, consistent with age-related activation of microglia in the African turquoise killifish brain (**Fig. 5c,d**).

Previous studies in mice found that microglia transcriptomes could be used to build highly accurate single-cell transcriptional aging clocks^82,83^. Thus, we decided to build microglia aging clock classification models trained using microglia from GRZ animals, which could then be used to evaluate predicted aging rates in microglia from longer-lived ZMZ-1001 animals (**Fig. 5e**; see **Methods**). Consistent with previous work in mice^82,83^, we were able to build highly accurate aging classification models with 2 algorithms, specifically elasticNet [eNet] and random forest [RF] (**Extended Data Fig. 12a-c**). Training performance was strong, with AUROC performance metric of ∼0.85 for both algorithms (**Extended Data Fig. 12a,b**). In addition, analysis of balanced accuracy for our trained models also showed strong accuracy on withheld testing data (>75%; **Extended Data Fig. 12c**), consistent with successful model training. We then applied our models to predict the probability of any microglia being from the brain of an “old” animal (defined as 16-week-old GRZ fish based on training framework; **Fig. 5e**; see **Methods**), using both withheld testing microglia single nuclei transcriptomes from GRZ fish and from ZMZ-1001 fish. Consistent with expectations, microglia from middle-aged (∼10-week-old) GRZ fish showed intermediate predicted probabilities of being old compared to those from young (∼6-week-old) or old (∼16-week-old) GRZ fish (**Fig. 5f**; **Extended Data Table 7a**). Surprisingly, microglia from female ZMZ-1001 fish did not show higher median predicted probability of being old (>60%) until the geriatric (26-week-old) time point, although those from male ZMZ-1001 fish reached that threshold at the old age time point (16-week-old; **Fig. 5f**; **Extended Data Table 7a**).

Finally, we examined our models to identify the top genes whose expression contributed most strongly to model performance in both algorithms (**Fig. 5g**; **Extended Data Table 7b**; see **Methods**). Interestingly, *helz2*, one of the genes recurrently upregulated with age across cell types, was among top predictors of microglia age (**Fig. 5g**). In addition, among top predictors we also found *b2m*, which encodes Beta-2 microglobulin, and *cd9*, which encodes a cell surface glycoprotein of the tetraspanin family, both previously described as DAM marker genes^84,85^ (**Fig. 5g**). Another top predictor for microglia age was *Ifi27l2a*, which was recently identified as a key regulator of microglia age-related inflammation in mice^86^, again suggesting conservation of aging phenotypes from fish to mammals (**Fig. 5g**). Consistent with their impact on the accuracy of our models, top predictor genes showed trends for age-related regulation across sex and strains (**Fig. 5g**).

### Coordinated changes in transcription factor [TF] activity across cell types in the aging African turquoise killifish brain

To identify potential master regulators of turquoise killifish brain aging across sex, cell types, and strains, we next decided to evaluate measures of TF activity across our multi-omic resource, using (i) differential predicted TF regulon activity (snRNA-seq) and (ii) differential TF motif accessibility (snATAC-seq and bulk ATAC-seq; **Fig. 6a**; **Extended Data Fig. 13-15**; see **Methods**).

**Figure 6:**
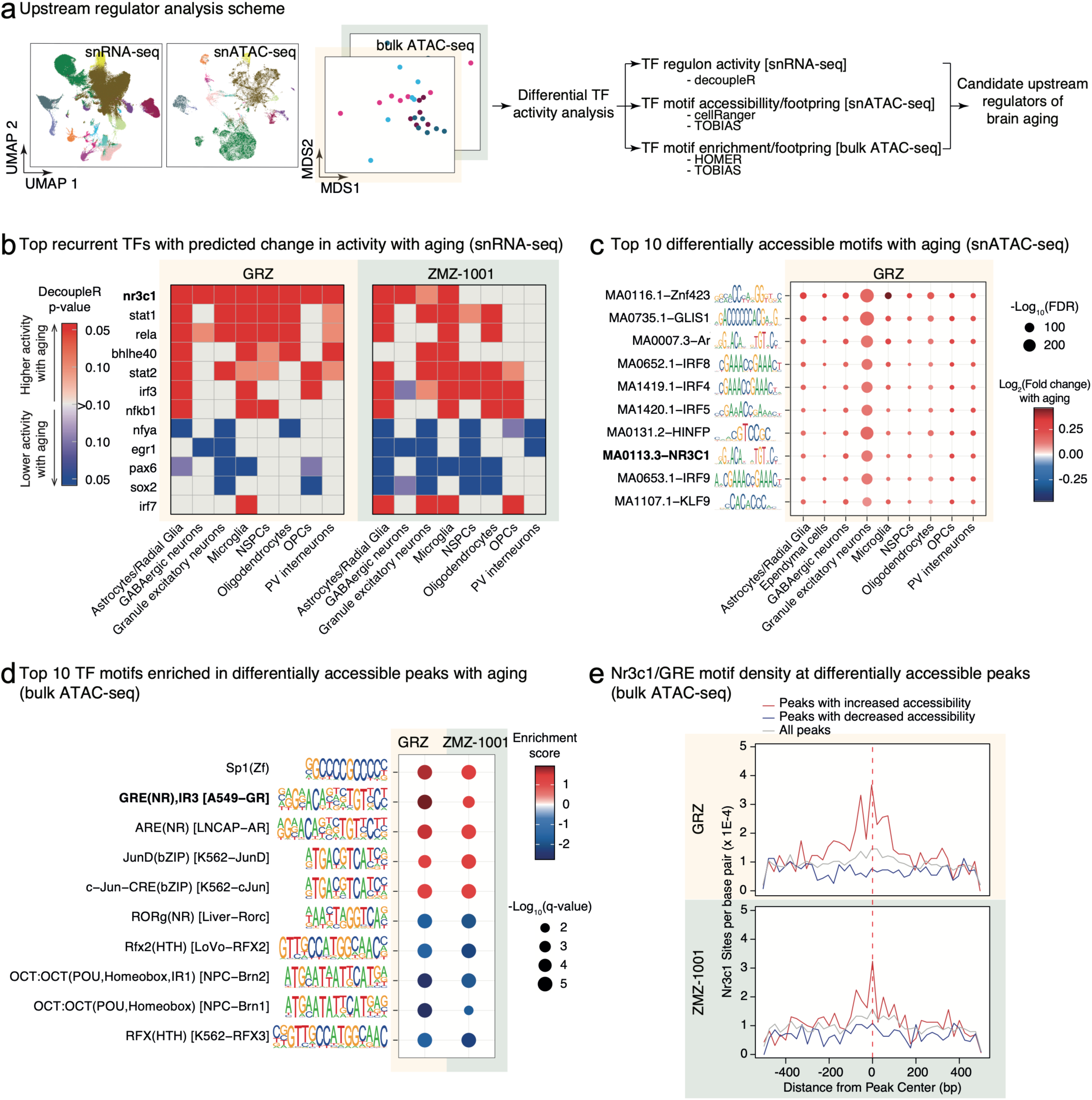
Multi-omic analysis of differential transcription factor activity in aging African turquoise killifish brains. (**a**) Scheme of the analytical pipeline to identify upstream regulators driving age-related changes in the African turquoise killifish brain. We leveraged data from our snRNA-seq, snATAC-seq and bulk ATAC-seq to quantify differential transcription factor [TF] activity, using TF regulon scoring (snRNA-seq), TF footprint accessibility analysis (snATAC-seq and bulk ATAC-seq), and known TF motif enrichment analysis (bulk ATAC-seq). (**b**) Heatmap of recurrent TF regulons with predicted age-related changes in activity using decoupleR in the snRNA-seq data derived from GRZ and ZMZ-1001 fish. To be included in this summary, TFs had to reach p < 0.05 in ≥4 cell types in either strain. Data was discretized for reporting: TF regulons with increased (decreased) activity are colored in red (blue), and lighter colors were used to indicate regulons reaching p < 0.1 as an additional line of evidence. (**c**) Differential TF binding site accessibility analysis from cellRanger-atac using findMarker statistics from GRZ snATAC-seq data. The top 10 most significantly differentially accessible motifs that showed accessibility change in all 9 QC tested cell types at FDR < 5% are reported. (**d**) Top TF binding motifs enriched in significantly differentially accessible bulk ATAC-seq peaks in both strains according to HOMER. The top 5 motifs enriched in peaks with increased or decreased accessibility with aging are reported. (**e**) Density of predicted Nr3c1/Glucocorticoid Receptor [GR] binding sites with respect to peak center in GRZ and ZMZ-1001 bulk ATAC-seq samples with (i) increased chromatin accessibility with aging (red line), (ii) decreased chromatin accessibility with aging (blue line), and (iii) all detected peaks regardless of chromatin accessibility changes with aging (grey line). Motif density was computed using 25bp bins and the HOMER histogram output function. Note the specific enrichment in peaks with increased chromatin accessibility with aging.

First, we leveraged decoupleR^87^ to predict TF regulon activity changes with aging across cell types in the aging brains of GRZ and ZMZ-1001 fish (**Fig. 6a,b; Extended Data Fig. 13a; Extended Data Table 8a,b**; see **Methods**). To zoom in on potential master regulators of brain aging regardless of cell types, we then filtered decoupleR results to identify TFs with significant consistent changes in predicted activity in ≥4 cell types in either the GRZ or the ZMZ-1001 strain (decoupleR p-value < 0.05), which yielded 12 candidate TF regulons with predicted age-related changes in activity across cell types (**Fig. 6b**). Among top candidates from this analysis, we identified several inflammatory signaling-related TF encoding genes, including glucocorticoid receptor *nr3c1*, inflammatory regulating *stat1*, *stat2*, *rela* and *nfkb1*, as well as interferon regulated *irf3* and *irf7* (**Fig. 6b**). To note, there was no clear trend for age-related changes in the expression for genes encoding these TFs (**Extended Data Fig. 13b**), suggesting that predicted TF activity changes are not a mere consequence of overall gene expression changes. However, we note that the genes encoding two top TFs, *irf3* and *stat1,* were among the top differentially genes shared across cell types from our snRNA-seq (**Fig. 3d**) that underwent RNAscope validation, providing a potential mechanism for predicted differential activity for these specific TFs.

We next asked whether predicted age-related changes in TF regulon activity across brain cell types in the African turquoise killifish were conserved in mammals (*i.e*. mouse, human) or were specific to this species (**Extended Data Fig. 14a-d**; see **Methods**). Specifically, we collected publicly available bulk RNA-seq and snRNA-seq datasets across brain regions with aging from mice^41,76,82,88–90^ and humans^91–93^, and evaluated predicted TF regulon activity using decoupleR as before (see **Methods**). We then focused on the top recurrently age-misregulated TF regulons identified in our African turquoise killifish dataset (**Fig. 6b**). Although predicted TF regulons with decreased activity in the aging brain tended to show relatively modest cross-species consistency, we noted strong concordance of our top TF regulons upregulated with turquoise killifish brain aging in our reanalysis of publicly available mouse and human brain aging bulk and snRNA-seq datasets (**Extended Data Fig. 14a-d**). Specifically, we noted either significant upregulation of the *Nr3c1/NR3C1* regulons (decoupleR p-value < 0.05), or suggestive trends (decoupleR p-value < 0.1), with mouse and human brain aging across many cell types and brain regions (*i.e*. 56 conditions evaluated for mouse aging, 25 of which with p < 0.05 and an additional 5 with p < 0.1; 27 conditions tested for human brain aging, 21 of which with p < 0.05; **Extended Data Fig. 14a-d**; see **Methods**). Conservation of differential TF regulon activity was also strong for *Stat1/STAT1*, *Rela/RELA* and *Nfkb1/NFKB1*, but weaker for *Irf3/IRF3*, *Irf7/IRF7* and *Stat2/STAT2* (**Extended Data Fig. 14a-d**). Together, these analyses suggest that several TFs show predicted patterns of activation across cell types in the aging turquoise killifish brain, patterns which are at least partially conserved with mouse and human brain aging.

Second, we analyzed our snATAC-seq data to identify potential changes in predicted TF footprint accessibility with aging using cellranger-atac and TOBIAS^94^ (**Fig. 6a,c**; **Extended Data Fig. 15a-c**; **Extended Data Table 8c,d**). We found that the chromatin accessibility footprint of many TFs was impacted with aging across cell types (**Fig. 6c**; **Extended Data Fig. 15a-c**). Consistent with our findings from snRNA-seq, we saw significantly increased chromatin accessibility with aging around canonical NR3C1 motifs (MA0113.3) and canonical IRF motifs (MA0652.1, MA1419.1, MA1420.1, MA0653.1) across cell types (**Fig. 6c; Extended Data Fig. 15d**). We also observed significantly increased chromatin accessibility with aging around canonical KLF9 motifs (MA1107.1), which was recently identified as a key downstream effector/mediator of glucocorticoid receptor activity in zebrafish^95^. Finally, we also observed significantly increased chromatin accessibility with aging around canonical Androgen Receptor (AR) motif (MA0007.3), regardless of animal sex (**Fig. 6c**), suggesting potential changes in sex steroid signaling shaping turquoise killifish brain aging.

Third, we evaluated evidence of differential TF activity in our bulk ATAC-seq data using a 2-pronged approach: (i) motif enrichment analysis in ATAC-seq peaks showing differentially accessibility with aging and (ii) differential TF footprint analysis using TOBIAS^94^ (**Fig. 6d; Extended Data Fig. 16a-c**; **Extended Data Table 8e-i**). Interestingly, we identified significant enrichment of the Glucocorticoid Response Element [GRE] and Androgen Response Element [ARE] motifs in bulk ATAC-seq peaks showing increased chromatin accessibility with aging (**Fig. 6d**). Consistently, TOBIAS footprint analysis of bulk ATAC-seq data revealed significantly increased chromatin accessibility with aging around canonical NR3C1 (MA0113.3) and Androgen Receptor (AR) motifs (MA0007.3), regardless of animal sex and strain (**Extended Data Fig. 16c**). To note, we observed high average canonical GRE motif density in bulk ATAC-seq peaks that show increased accessibility with aging compared to peaks with decreased accessibility with aging, or the entire peak set (*i.e*. “background”) (**Fig. 6e**).

Together, when integrating information from multiple ‘omic’ layers across sex, strains and cell types and across species, we identify one particular TF, nr3c1 (the turquoise killifish glucocorticoid receptor), that constitutes a particularly compelling candidate master regulator of brain aging – with multiple lines of ‘omic’ evidence supporting increased activity (snRNA-seq, snATAC-seq, bulk ATAC-seq) and conservation of increased predicted activity in publicly available mouse and human brain aging datasets.

### African turquoise killifish show increased cortisol production with aging across sex and strains

Cortisol is the main glucocorticoid in teleosts^96^, and its circulating levels are known to dramatically increase in response to stress^96^, including in the African turquoise killifish^97^. Thus, we reasoned that our observations of increased activity of glucocorticoid receptor signaling across cell types in the aging turquoise killifish brain may be the result of an age-related increase in cortisol production. Since teleost synthesize cortisol in their kidney^96^ (**Fig. 7a**), we next measured the amount of cortisol in banked kidney tissue (**Fig. 7b**). Consistent with our predictions, we observed a significant increase of cortisol levels in aging kidney tissue from both GRZ and ZMZ-1001 strains, regardless of sex (**Fig. 7b**). Interestingly, increased circulating glucocorticoid levels have been reported with mouse and human aging and have been hypothesized to drive impairments in spatial memory^98,99^. Thus, age-related increases in cortisol production may lead to aberrantly increased glucocorticoid receptor signaling, notably in the brain, ultimately driving aspects of brain aging in the African turquoise killifish.

**Figure 7:**
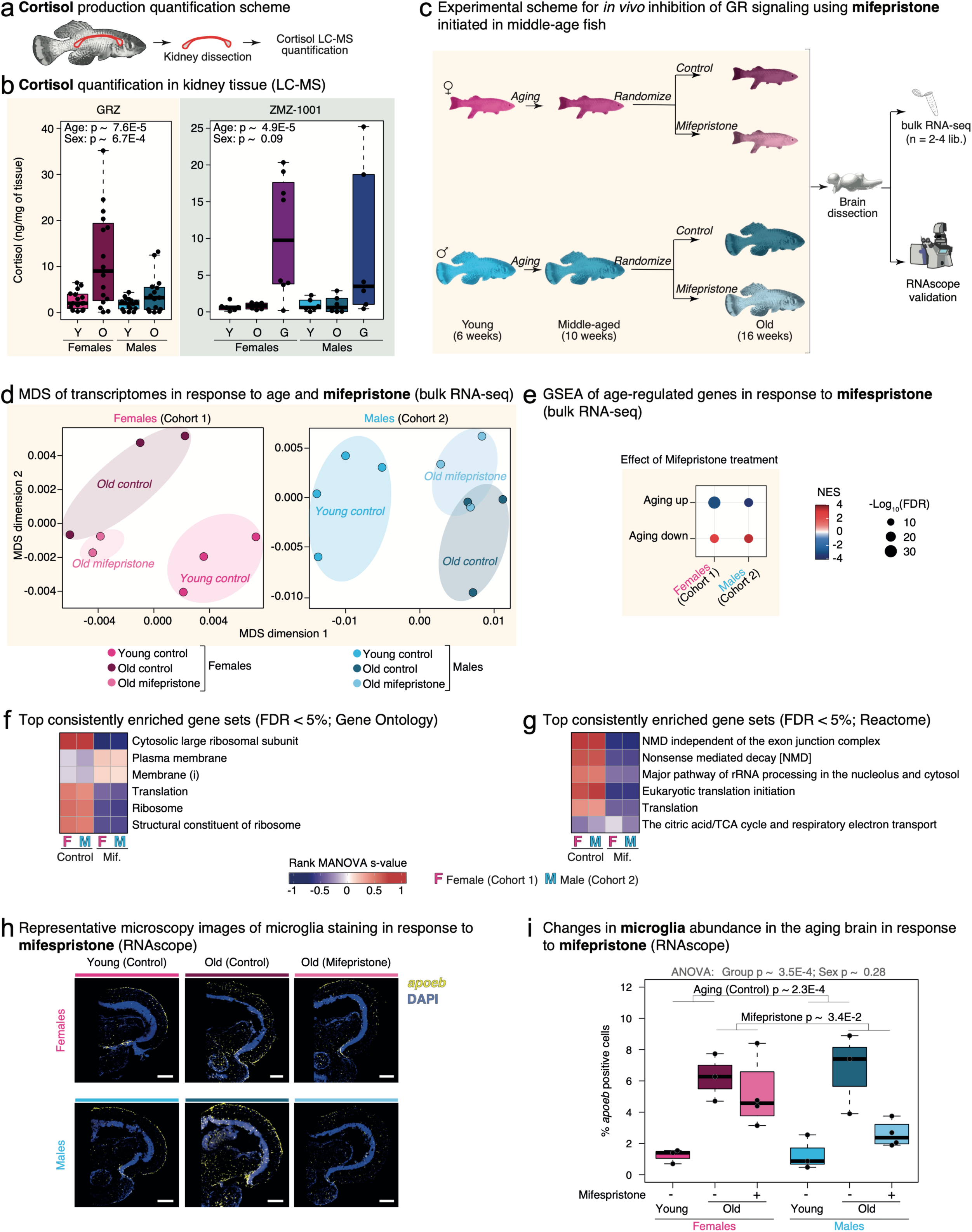
Importance of glucocorticoid signaling in shaping the molecular landscape of African turquoise killifish brain aging. (**a**) Schematic of kidney anatomical position in the African turquoise killifish. (**b**) Quantification of Cortisol, the main form of glucocorticoids in teleosts, per unit of mass of kidney tissue by LC-MS. One significant outlier among GRZ young female samples and 2 significant outliers among GRZ young male samples were identified using the Rosner test and are omitted from the plot and analysis. The effects of aging and sex were evaluated using non-parametric ART-ANOVA. (**c**) Schematic of experiment to determine whether GR antagonism with food-based mifepristone supplementation initiated at 10 weeks of age (middle-age) can revert aspects of brain aging. (**d**) Multidimensional scaling [MDS] analysis of bulk RNA-seq brain datasets. Note that transcriptomes of brains from old fish that underwent mifepristone supplementation are grouping between the young and old control brain transcriptomes, consistent with a “younger” transcriptomic phenotype. (**e**) GSEA of age-regulated genes in response to mifespristone treatment in our female and male bulk RNA-seq cohorts. (**f-g**) Top Gene Ontology (**f**) and Reactome (**g**) gene sets regulated consistently across the female and male cohort with brain aging and in response to mifepristone supplementation. The top 6 most enriched gene sets are represented, all with FDR < 5%. Heatmaps are colored based on the s statistic of a rank MANOVA test performed with mitch. To disambiguate GO naming redundancy: membrane (i) refers to GO:0016020, and membrane (ii) refers to GO:0016021. (**h**) Representative maximum z-projected images of microglia (*apoeb* signal; yellow) overlaid with DAPI (blue). Age and treatment are organized by column: young controls (left), old controls (middle), and old mifepristone-treated (right) animals. All scale bars are 250mm in length. (**i**) Quantification of microglial abundance in African turquoise killifish brain samples using *apoeb* for *in-situ* hybridization by RNAscope. Values for each animal are reported as the average QuPath inferred values obtained across 3 Z-planes. The effect of treatment group (young control, old control, old mifepristone) and sex were evaluated using ANOVA. Normality of ANOVA residuals was confirmed using a Shapiro-Wilkes test (p > 0.05). Significance of the effect of treatment group, regardless of sex, was evaluated using the Tukey test for post-hoc analysis and is reported on the graph.

### Glucocorticoid receptor antagonism through mid-life-initiated mifepristone treatment rescues aspects of brain aging

Our results support the notion that, with aging, increased production of cortisol leads to chronic activation of glucocorticoid signaling through its receptor nr3c1, which promotes molecular aspects of aging across brain cell types (**Fig. 6b-e, 7b**). These observations prompted us to ask whether antagonizing glucocorticoid signaling before cortisol production reaches its peak at old age could help delay or mitigate aspects of brain aging. In addition to its original drug development purpose as a progesterone receptor blocker, mifepristone is a potent competitive antagonist of the glucocorticoid receptor, and the only FDA-approved for the treatment of chronically elevated cortisol levels (*i.e*. Cushing’s syndrome)^100^. Thus, we leveraged mifepristone to antagonize glucocorticoid signaling in African turquoise killifish of the shorter-lived GRZ strain, initiating treatment at middle-age (10 weeks of age), in females and males, then analyzing the effects of treatment at old age (16 weeks of age; **Fig. 7c**; see **Methods**). To note, this experimental paradigm did not significantly impact animal morphometric parameters (*i.e*. length and weight), suggesting it is well tolerated (**Extended Data Fig. 17a-b**).

Next, we performed bulk RNA-seq analysis of brain tissue harvest from 2 independent cohorts of African turquoise killifish with our mifepristone exposure paradigm, one female cohort and one male cohort (**Fig. 7c**; **Extended Data Fig. 18a**; see **Methods**; **Extended Data Table 9a-d**). Interestingly, MDS analysis showed that, in both cohorts, brain transcriptomes of fish treated with mifepristone clustered in an intermediate position compared to young and old control fish, consistent with a delayed or slowed down aging phenotype (**Fig. 7d**). Using gene set enrichment analysis [GSEA], overlap analysis and correlation analyses, we observed that age-regulated genes were significantly regulated in the opposite direction to that of aging in the brain transcriptomes of fish treated with mifepristone (**Fig. 7e**; **Extended Data Fig. 18b,c**). To note, the impact of aging and mifepristone on bulk RNA-seq profiles was consistent between the 2 cohorts (**Extended Data Fig. 18d**), and similar reversal of age-related transcriptional changes were also observed when using a meta-analysis approach (**Extended Data Fig. 18e**; **Extended Data Table 9e,f**; see **Methods**).

We then asked which functional gene classes were significantly impacted by mifepristone treatment in the aged turquoise killifish brains (**Fig. 7f,g**; **Extended Data Fig. 18g,h**; **Extended Data Table 9 g,h**). Multi-contrast pathway enrichment analysis showed a reversal of age-related pathway regulation upon mifepristone treatment, especially relating to proteostasis (*e.g*. GO “Ribosome”, GO and Reactome “Translation”, Reactome “Eukaryotic translation initiation), membrane biology (*e.g*. GO “Membrane”, “Plasma Membrane”), and RNA metabolism (*e.g.* Reactome “Nonsense mediated decay”) (**Fig. 7f,g**; **Extended Data Table 9g,h**). Consistently, overrepresentation analysis of genes showing reversal of age-regulation upon mifepristone treatment identified by meta-analysis showed significant enrichment for similar pathways (**Extended Data Fig. 18f,g**; **Extended Data Table 9i,j**). In addition to shared functional categories, this analysis also revealed enrichment for synapse-related genes (*i.e*. Reactome “Transmission across chemical synapses”) among genes normally downregulated with aging that were upregulated in the brains of mifepristone-treated fish (**Extended Data Fig. 18g**; **Extended Data Table 9i,j**).

Next, we asked whether mifepristone treatment had any impact on the microglia compartment of the aged brain, since microglia expansion was a major feature of African turquoise killifish brain aging (**Fig. 2**). We reasoned that previous work in mice showed that elevated glucocorticoid signaling leads to *in vivo* proliferation of microglia, promoting neuroinflammation^101^, suggesting a potential mechanistic link. Interestingly, we observed non-significant trends for decreased transcript levels of *apoeb*, our top microglia marker in the turquoise killifish, in the brains of old fish treated with mifepristone (**Extended Data Fig. 19a**). Consistently, deconvolution analysis of our bulk RNA-seq datasets showed that the predicted proportion of microglia significantly increased with aging in control animals, although this was no longer significant when comparing young animals with old animals that received mifepristone treatment (**Extended Data Fig. 19b**). Thus, we leveraged our RNAscope ISH cell identity panel to directly quantify microglia abundance in the brain of young control, old control and old mifepristone-treated female and male GRZ fish (**Fig. 7h,i**). As before (**Fig. 2g**), we saw significant microglia expansion in the brains of young vs. old turquoise killifish receiving the control diet (**Fig. 7h,i**), validating our previous findings. Excitingly and consistent with our computational predictions from bulk RNA-seq, we observed a significant reversal of the age-related microglia expansion in the older animals that received mifepristone compared to controls, as measured by the percentage of *apoeb*-positive cells (**Fig. 7h,i**). Together, our observations suggest that mid-life-initiated treatment with mifepristone is sufficient to blunt aspects of brain aging (i.e. transcriptomic response, microglia expansion).

## Discussion

In this study, we generated and characterized a comprehensive atlas of brain aging in the naturally short-lived African turquoise killifish, combining single nuclei transcriptomics to single nuclei and bulk chromatin accessibility mapping. Importantly, our atlas includes data from both sexes, as well as two independent genetically distinct strains: the standard inbred short-lived GRZ strain, as well as the longer-lived, wild-derived ZMZ-1001 strain. We were able to define 16 distinct brain cell types and their associated marker genes, which will have great utility for further work into understanding brain aging in the African turquoise killifish model in a cell-aware manner.

Our multi-omic atlas allowed us to identify key changes, robust to sex and genetic background, in cell type proportions and in gene regulatory programs in the brain of this species with aging. Age-related changes in the brain were similar across genetic strains and sex, albeit occurring at a slower pace in longer-lived ZMZ-1001, consistent with the influence of strain on lifespan^102^ and with a model of overall “rate of aging” rather than of abrupt cliff-like transitions. Our data showed age-regulation of gene expression of previously defined human/mouse aging signatures (*i.e*. GTEx, CAG, Senmayo), supporting broad conservation of transcriptional hallmarks of brain aging across vertebrate models^103^. More broadly, our functional enrichment analyses revealed enrichment of many pathways related to the hallmarks of aging^1^: dysregulation of pathways related to inflammation, NAD metabolism, DNA-damage, ECM, protein homeostasis, and membrane metabolism. In addition, age-related changes in pathways relating to RNA processing and translational regulation were identified across cell types, consistent with a recent multi-omic study of brain aging in turquoise killifish of the long-lived MZMZ-0410 strain^30^. Together, our dataset supports the usefulness of the turquoise killifish as a short-lived model organism with naturally compressed lifespan to accelerate research into brain aging mechanisms.

The African turquoise killifish brain does not share the regenerative capacity of other teleosts, including the much longer-lived zebrafish^20^. Turquoise killifish have been previously reported to exhibit age-related loss of dopaminergic and noradrenergic cells^104^, as well as changes in serotonin circuitry^105^. In regenerative organisms, microglia contribute to much of the age-related hemostasis of the brain to prolong health and lifespan^106,107^. Our analyses revealed a clear, significant expansion of the microglia population with aging in the turquoise killifish, which was robust to sex and strain. To note, microglial signatures relating to inflammatory tone (*e.g*. DAM, IRM, LDAM, TIM, and M1/M2 polarization) showed similar age-related dynamics in turquoise killifish microglia as that reported in human or mice^108–110^. Independent validation using RNA-ISH also identified an increase in the density and volumetric properties of microglia. To note, the mechanistic driver behind microglia expansion in the aging turquoise killifish brain remains unclear at this time. Indeed, in zebrafish, adult microglia can derive from two compartments: (i) yolk sac-derived, locally-proliferating microglia, and (ii) colonizing microglia from the dorsal-aorta axis and the head kidney^111–113^. Nonetheless, our data suggests that the turquoise killifish can be a powerful model to understand conserved age-related changes in microglia function and regulation in a naturally short-lived vertebrate.

We next demonstrated the usefulness of our aging turquoise killifish brain resource to identify new, actionable pathways that are dysregulated with brain aging. Specifically, our multi-level analysis of TF activity predicted a robust increase in the activity of the glucocorticoid receptor, *nr3c1*, at the transcriptomic and epigenomic levels across cell types in the aging brain. Further, by comparing our data to existing human and mouse bulk/single cell RNA-seq datasets, we found that the age-related increase in predicted glucocorticoid signaling was not a model-specific anomaly or idiosyncrasy, but a conserved feature of mammalian brain aging. Glucocorticoid receptors are key regulators of the immune and stress response^114^, can modulate inflammatory gene expression^115^, and control neuronal survival^116^. Furthermore, microglial glucocorticoid signaling restrains inflammatory injury and support neuronal survival during stress and aging^117–119^. In both murine and human models, chronic glucocorticoid receptor activation leads to changes in adrenal cortex function and structure^120^, as well as neurodegeneration, synaptic loss, cognitive decline, and Alzheimer’s disease pathology^99,121–123^. Genetic variation in the human *NR3C1* gene influences several longevity-associated phenotypes^124–128^. However, to our knowledge, whether glucocorticoid signaling is an actionable target to rescue aspects of physiological brain aging was not previously investigated.

To assess the potential mechanistic involvement of increased glucocorticoid receptor signaling in driving brain aging phenotypes, we leveraged systemic glucocorticoid receptor antagonism through the drug mifepristone. In addition to its progesterone receptor blocker activity, mifepristone is a potent competitive antagonist of the glucocorticoid receptor and remains the only FDA-approved for the treatment of chronically elevated cortisol levels (*i.e*. Cushing’s disease)^100^. Glucocorticoid antagonism has been shown to dampen microglial activation^129^, sensitization^130^, and proliferation^101^, and augment recovery from traumatic head injury^131^. In the context of disease, mifepristone can reduce amyloid and tau pathology and can improve cognitive outcomes both preclinically and clinically^132–134^. Thus, it presents an attractive target in the context of brain aging. To limit potential off-target toxicity, we decided to initiate treatment at mid-life, after the peak reproductive window, but before cortisol levels increase with age. Importantly, mid-life-initiated mifepristone treatment significantly reversed key aspects of turquoise killifish brain aging in both female and male fish, including age-related (i) transcriptional changes in proteostasis, membrane biology, and RNA metabolism-related pathways and (ii) expansion of the microglia population in the brain. Together, our data supports the notion that glucocorticoid antagonism is a powerful strategy to mitigate aspects of vertebrate brain aging.

In summary, we generated a comprehensive multi-omic atlas of brain aging at single-cell resolution across sex and strains in the naturally short-lived African turquoise killifish model. This rich resource allowed us to identify a conserved driver of vertebrate brain aging, glucocorticoid signaling, and demonstrate that it can be targeted pharmacologically to mitigate the onset of robust aging phenotypes. Although we identify robust ‘omic’ remodeling in the aging brain, future studies (beyond the scope of our resource) should examine to which extent these molecular changes may potentially impact cognitive or behavioral aging. Together, our work provides a robust, rich resource for the study of vertebrate brain aging.

## Material and Methods

### Fish Husbandry

African turquoise killifish were reared in accordance with approved IACUC protocols for *Nothobranchius furzeri* at the University of Southern California. Fish used in this study were generated on approved IACUC protocols 21215 and 21282. African turquoise killifish from the GRZ and ZMZ-1001 strains of turquoise killifish were raised and euthanized as previously described^34^. Briefly, hatchlings were fed a diet of with live brine shrimp until reaching adulthood, and adult fish were fed freeze-dried bloodworms. Fish were fed twice daily during normal weekdays, and generally once daily over weekends and holidays, consistent with approved IACUC protocols. Females and males were used for experiments as indicated in the results.

### African turquoise killifish lifespan data collection

Lifespan data was derived from historical survival data from our African turquoise killifish colony at USC (collected 2019-2025). Time from hatching to humane endpoint or natural death was recorded. All reported data is derived from single-housed fish, to limit the impact of death or injuries from frequent fighting in this species. A portion of the historical survival data for the GRZ strain (2019-2022) was also used in a previous study from our lab^33^.

### Genomic DNA isolation and whole genome shotgun sequencing [WGS]

Genomic DNA was isolated from skeletal muscle tissue of 2 male fish from the GRZ and ZMZ-1001 line each, using a commercial genomic DNA purification kit (Thermo Fisher K0512) according to the manufacturer’s instructions. Male fish were used in the case for maximum capture of genetic variation in the strains, as they are the heterogametic sex in this species^38^. DNA extraction was started by lysing frozen muscle tissue in the kit’s lysis buffer using a dounce homogenizer. Purified genomic DNA was then sent to the Novogene corporation (USA) for library construction and sequencing, who shared sequencing results for analysis. Raw FASTQ reads have been deposited to the sequence read archive under accession PRJNA1121423.

### Analysis of African turquoise killifish genomic variation across laboratory isolates

In addition to WGS data on GRZ and ZMZ-1001 fish from our USC colony, we also obtained publicly available WGS data for other African turquoise killifish laboratory isolates for analysis of genetic profiles (**Extended Data Table 10a**). For both our and the publicly available data, paired-end FASTQ reads were mapped to the NCBI GCF_001465895.1 turquoise killifish reference genome version using bwa 0.7.17-r1188^135^. Potential PCR duplicates were marked using gatk v4.1.9.0^136^ ‘MarkDuplicates’ function. Variants were called using samtools v0.1.19 and bcftools v1.20 mpileup^137^. Called variants were obtained in the standard vcf format for further processing.

Principal Component Analysis (PCA) of genotypes of turquoise killifish isolates was performed using plink v1.90b6.21^138^. To note, genotype data was first pruned using parameters “ –-indep-pairwise 100 50 0.9” before calculation of principal components. As expected, there was a broad separation between yellow-tailed and red-tailed turquoise killifish isolates. Importantly, despite a large lifespan difference, GRZ and ZMZ-1001 fish are relatively close in the genetic PCA space, consistent with the original geographic distribution of their founding isolates in the wild.

### Brain dissection and nuclei isolation from snap-frozen brain tissue for snRNA-seq

Whole brains were dissected from GRZ and ZMZ-1001 male and female turquoise killifish and immediately snap frozen by placing into Eppendorf tubes on dry ice. Frozen brains were then stored at –80°C until dissociation. Brains from four age groups were collected in this study corresponding to approximately 6, 10, and 16 weeks post hatching for GRZ and ZMZ-1001 fish, as well as additional 26 weeks post-hatching time point for the longer-lived ZMZ-1001 fish. Numeric ages were used where relevant in downstream analyses, although in many cases age groups (*e.g*. Young, Middle-Age, Old and Geriatric) were used for ease of representation. Single nuclei suspensions were prepared from frozen brain samples in five batches processed on different days. To produce enough nuclei for downstream sequencing, two fresh frozen brains from the same group were pooled to generate high quality brain nuclei samples for snRNAseq. Nuclei suspensions were prepared in accordance to the protocol we previously optimized to isolate nuclei from frozen African turquoise killifish brains^139^.

Briefly, frozen brains were homogenized by dounce homogenization, filtered through a 70 µm strainer (Miltenyi Biotec), washed and pelleted by centrifugation, subjected to debris removal via gradient centrifugation using a debris removal solution (Miltenyi Biotec), washed and filtered through a 40µm strainer (Flowmi), and assessed for quality by light microscopy and flow cytometry as previously described^139^.

### Single-nuclei RNA-seq [snRNA-seq] library preparation

Single-cell libraries were prepared using Chromium Next GEM Single Cell 3L GEM, Library & Gel Bead Kit v3.1 (10xGenomics, PN-1000121) according to manufacturer’s instructions (10xGenomics User Guide Chromium Next GEM Single-cell 3′ Reagent Kits v3.1 (CG000204, Rev D)). Using nuclei concentration estimates from the MACSQuant Analyzer 10 (Miltenyi Biotec, #130-096-343), nuclei suspensions were loaded for a targeted cell recovery per sample of 12,000 nuclei (pilot GRZ cohort) or 7,000 nuclei (all other cohorts). Samples were run on Chromium Next GEM Chip G (10xGenomics, PN-2000177) per manufacturer’s instructions, albeit with the addition of 2 pre-amplification cycles at the cDNA amplification cycles to compensate for the inherently RNA-poor status of nuclei. Partially amplified and final single-nuclei RNA-seq libraries were quantified by using a Qubit® 3.0 Fluorometer (Thermo Fisher Scientific, Q33216) and the dsDNA high sensitivity assay (Invitrogen, Q32854). Completed single-nuclei RNA-seq libraries were assessed for quality on the 4200 TapeStation system (Agilent Technologies; G2991A) with a High Sensitivity D1000 DNA ScreenTape (Agilent Technologies 50675584). Libraries were sequenced on an Illumina Novaseq 6000 generating 150 bp paired-end reads at the Novogene Corporation (USA). Raw FASTQ reads have been deposited to the sequence read archive under accession PRJNA1105049.

### Brain dissection and nuclei isolation from fresh brain tissue for single nuclei and bulk ATAC-seq

To avoid potential artefacts from freezing on chromatin structure, African turquoise killifish brains for use in single nuclei and bulk ATAC-seq experiments were freshly dissected and immediately subjected to nuclei isolation procedures. To reduce confounding circadian effects, brains were dissected from sacrificed turquoise killifish between 10am and 12pm, as done for the snRNA-seq libraries. Single nuclei suspensions were prepared in accordance with the “Nuclei Isolation from Mouse Brain Tissue for Single Cell ATAC Sequencing” protocol from 10X Genomics^140^, following instructions for “Fresh or Cryopreserved Tissue” with the following modifications: (i) a modified lysis buffer with 75% of the stated Nonidet P40 Substitute concentration was used, and (ii) a debris removal step using Miltyeni’s Debris Removal solution (Miltenyi Biotec, #130-109-398) was included to minimize debris contribution to samples. Only GRZ strain male and female turquoise killifish corresponding to the 6-week and 16-week timepoints were used for snATAC-seq, corresponding to 4 biological groups, in 2 separate cohorts. Brains used for snATAC-seq were processed in two independent batches with each group present to reduce potential batch effects. In each batch, nuclei were isolated from 3 independent fish per group, and pooled in an equinuclear fashion for equal representation in the final snATAC-seq libraries (for a total of 8 libraries).

For the bulk ATAC-seq dataset, individual fish brains were used to generate independent ATAC-seq libraries using the Omni-ATAC protocol (for total of 48 libraries). For GRZ turquoise killifish, 6-week and 16-week-old killifish were used to make bulk ATAC-seq libraries across 6 biological replicates per group.

### Single-nuclei ATAC-seq [snATAC-seq] library preparation

Single-cell libraries were prepared using the Chromium Next GEM Single Cell ATAC Reagent Kits v1.1 (10X Genomics, PN-1000175) according to manufacturer’s instructions (10xGenomics User Guide Chromium Chromium NextGEM Single Cell ATAC Reagent Kits v1.1 (CG000209 Rev F)). Using nuclei concentration estimates from the MACSQuant Analyzer 10 (Miltenyi Biotec), nuclei suspensions were loaded for a targeted recovery of 7,000 nuclei per sample. Samples were run on Chromium Next GEM Chip H (10xGenomics, PN-1000162) per manufacturer’s instructions. Single-nuclei ATAC-seq libraries were quantified by using a Qubit® 3.0 Fluorometer (Thermo Fisher Scientific, Q33216) and the dsDNA high sensitivity assay (Invitrogen, Q32854). Completed single-nuclei RNA-seq libraries were assessed for quality on the 4200 TapeStation system with a High Sensitivity D1000 DNA ScreenTape. Libraries were sequenced on an Illumina Novaseq 6000 generating 150 bp paired-end reads at the Novogene Corporation (USA). Raw FASTQ reads have been deposited to the sequence read archive under accession PRJNA1106410.

### Bulk ATAC-seq library preparation

Using brain nuclei concentration estimates from the MACSQuant Analyzer 10, we aliquoted 50,000 brain nuclei from individual nuclei preparations to process through the Omni-ATAC protocol^35^. Libraries were quality controlled on the 4200 TapeStation system with High Sensitivity D1000 or D5000 DNA ScreenTapes. Libraries were multiplexed and sequenced on the Illumina Novaseq 6000 platform as paired-end 150bp reads at the Novogene Corporation (USA). Raw FASTQ reads have been deposited to the sequence read archive under accession PRJNA1100604.

### BLAST identification of mouse, human, and zebrafish homologs for annotation

Homology mapping of African turquoise killifish genes was performed as previously^34^. Briefly, predicted protein sequences were obtained from the African turquoise killifish genome version GCF_001465895.1 from NCBI. Mouse protein sequences were obtained from Ensembl Biomart version 107, which contains annotations derived from genome assembly version GRCm39 (downloaded 2022-10-10). Human protein sequences were obtained from Ensembl Biomart version 111, which contains annotations derived from genome assembly version GRCh38.p14 (downloaded 2024-03-11). Zebrafish protein sequences were obtained from Ensembl Biomart version 105, which contains annotations derived from genome assembly version GRCz11 (downloaded on 2022-02-11). In all cases, NCBI BLAST 2.10.0+ was used to align the other species proteomes to the turquoise killifish predicted reference proteome to obtain the best BLAST hit using parameters “-best_hit_score_edge 0.05 –best_hit_overhang 0.25”. To reflect the relative phylogenetic distance, an e-value cutoff of 10^-3^ was used for mouse and human homology mapping, and 10^-5^ for zebrafish homolog mapping.

### snRNA-seq data analysis

The following pipeline was run for each set/cohort of single-cell datasets. A custom genome reference was assembled using the GCF_001465895.1 genome assembly^38^, which has been annotated by NCBI. The genome was hard masked using a turquoise killifish transposable element [TE]/repeat library from FishTEDB^141^ (downloaded on 2021-09-06). TE sequences were then added to the masked genome reference fasta as independent contigs, and as their own entries to the gene reference file gtf. CellRanger 6.1.2 (10x Genomics) was used to create the reference.

FASTQ read files were aligned to this turquoise killifish reference genome and quantified using CellRanger 6.1.2 “cellranger count” using option –-include-introns to account for low splicing of nuclear RNAs. The ambient RNA ‘soup’ was estimated and removed using the DecontX algorithm from package ‘celda’ v1.16.1 in R v4.3.1^142^ using the raw feature barcode matrix folder from cellranger to estimate ambient RNA. Quality control was performed in R v4.3.1 using Seurat v4.3.0.1^51,143–145^ by removing dead and low-quality cells by selecting for cells that contained between 250-5000 UMIs, <10% mitochondrial-mapping reads, < 25000 UMI counts and < 25% estimated background contamination according to DecontX.

A list of mouse cell cycle genes was obtained from the Seurat Vignettes (https://www.dropbox.com/s/3dby3bjsaf5arrw/cell_cycle_vignette_files.zip?dl=1), derived from a mouse study of cell cycle^146^. Cell cycle phase was predicted using homologous African turquoise killifish cell cycle genes based on BLAST unidirectional mapping (see above) and using the function CellCycleSorting() to assign cell cycle scores to each cell. Multiplets were annotated using a combination of Doubletfinder v2.0.3^147^ and scds v 1.16.0^148^ for each individual 10x Genomics library. Only cells called as singlets by both methods were considered to be high confidence singlets and used to create the final snRNA-seq cell atlas object.

After pre-processing, the 5 cohort datasets were merged into a single Seurat object in R v4.3.1 using Seurat v4.3.0.1. Due to the dataset size, gene expression was normalized on a global scale using “LogNormalize”, with a scale.factor of 10000. Dimensionality analysis revealed an optimal number of PCs of 19 (the minimum of the PC rank (i) where change of % of variation is more than 0.1% and (ii) which exhibits cumulative percent greater than 90% and % variation associated with the PC as less than 5), which was used for all downstream analyses (including UMAP construction). Next, 5,000 variable features were identified, and the dataset was scaled using the ScaleData function, regressing to the variables “nCount_RNA”, “nFeature_RNA”, “percent.mito”, “Phase”, and “Batch”. Harmony v0.1.1^149^ was used to integrate data across snRNA-seq cohorts and mitigate batch effects. We selected to focus Harmony integration using variables “Batch” and “Strain”, each with theta = 3, and the algorithm converged within 8 iterations.

### snRNA-seq cell type annotation

We first used Seurat v4.3.0.1 to perform unsupervised cell clustering with a single nearest neighbor resolutions and a resolution of 1.2, which was chosen to provide granularity pre-annotation with 36 clusters. This clustering served as the basis for following annotation efforts.

Two different R packages, scSorter v0.0.2^150^ (marker-based) and scMAGIC v0.1.0^151^ (reference-based) were used, each in conjunction with 2 reference sets. Specifically, scSorter was used using BLAST turquoise killifish homologs of (i) mouse brain cell type markers from PanglaoDB (accessed 2020-03-27), and (ii) zebrafish brain cell type markers from a single cell RNA-seq atlas obtained from the online supplement of the publication^39^ (see above for homology mapping). In parallel, for use with scMAGIC, our turquoise killifish snRNA-seq Seurat object was first converted to a “mousified” object using the BLAST mouse homologs of turquoise killifish genes. Then, scMAGIC was run using two reference annotated mouse snRNA-seq brain atlases^40,41^ using parameters: “method_HVGene = ‘SciBet_R’, method_findmarker = ‘Seurat’, cluster_num_pc = 19, min_cell = 10, method1 = ‘spearman’, percent_high_exp = 0.8 “. First pass predictions of cell type identity from both scSorter and scMAGIC runs were gated back to the original Seurat object for further processing.

We then proceeded to annotate each of the 1.2 SNN Seurat clusters using combined information from (i) scSorter and scMAGIC automated annotations, (ii) expression of cell proliferation markers *pcna* and *mki67*, (iii) top enriched markers for the cluster using Seurat FindAllMarkers with parameters only.pos = TRUE, min.pct = 0.25, logfc.threshold = 0.5. After annotation, 4 distinct population of neurons (labeled Neuron_misc_1..4) remained ambiguously annotated (*i.e*. with non-canonical main markers), although they displayed expression of glutamatergic markers vGluT1/ vGluT2, suggesting they constitute subtypes of glutamatergic neurons.

### R shiny application generation

An interactive R Shiny application for the killifish brain atlas was generated using ShinyCell2 (v1.0.0) and is publicly available at https://minhooki.shinyapps.io/killifish-brain-atlas/.

### snATAC-seq data analysis

A custom genome reference was assembled using the GCF_001465895.1 genome assembly^38^, which has been annotated by NCBI, modified using hardmasking of FishTEDB sequences (see above), as well as the non-redundant core vertebrate set of motif matrices from JASPAR (JASPAR2018_CORE_vertebrates_non-redundant_pfms_jaspar.txt), specifying “NC_011814.1” as the mitochondrial chromosome. cellranger-atac 2.1.0 (10x Genomics) was used to create the reference.

FASTQ read files were aligned to this custom reference genome and quantified using cellranger-atac 2.1.0 “cellranger-atac count”. CellRanger peak sets derived from individual libraries were loaded into R v4.3.1 using “makeGRangesFromDataFrame” from the GenomicRanges v1.52.1 package and reduced to a single merged peak set. Only peaks with a length < 10000 and > 20 bp were retained for downstream analysis (yielding 126,692 genomic ranges). CellRanger fragment files were also imported into R v4.3.1 for processing and filtered for low quality cells with ≤ 250 fragments. Each library was then processed using Signac v1.10.0^152^. Cells were quality filtered by selecting for cells that contained > 50% of reads in peaks, peak_region_fragments > 3000, peak_region_fragments < 40000, nucleosome_signal < 2, TSS.enrichment > 2 and mitochondrial < 1000. Potential doublets were annotated and removed using scDoublet from scDblFinder v1.14.0^153^. Normalization and linear dimensional reduction were conducted using RunTFIDF, FindTopFeatures (with min.cutoff = ‘q70’), RunSVD. For UMAP, we used the number of PCs derived from our snRNA-seq dataset (19) as an estimate of the dimensionality of brain nuclei in this species on the LSI reduction. A gene-activity matrix was computed using the “GeneActivity” function, which was normalized using LogNormalize.

The gene-activity assay was then used for integration with our snRNA-seq Seurat object to enable label transfer and cell annotation for the snATAC-seq dataset. Due to memory constraints, a random subset of the snRNA-seq dataset was used, restricted to 2,500 nuclei per annotated cell type from the GRZ strain (matching the strain of the snATAC-seq dataset). CCA was used for integration and label transfer to the snATAC-seq dataset using Seurat predictions. The annotated snATAC-seq object was then used for downstream analyses.

### Bulk ATAC-seq data analysis

Raw FASTQ reads were first trimmed using NGmerge^154^ with options “-v –z –n 2 –q 34” to remove dovetailing ends before further processing. Next, we used bowtie2 v2.5.1^155^ to map the reads to the GCF_001465895.1 turquoise killifish genome reference, using options “--sensitive –-no-discordant –-no-mixed –k 2 –p 2 –X 2500”. Next, we used MACS2 v2.2.7.1^156^ to call peaks on each individual sample using an effective genome size of 1.25e9. We next obtained a set of consensus ATAC-seq peaks in the turquoise killifish brain using MSPC v5.5.0^157^ and options “-c 20 –r bio –w 1e-5 –s 1e-9 –m Lowest” so as to yield a robust peak set with evidence from at least 20 independent libraries.

### Cell type prioritization analysis with Augur

For single nuclei RNA-seq and ATAC-seq datasets, we split objects by sex and strain to investigate the impact of aging in each homogenous biological group. We applied the Augur v1.0.3^43^ algorithm to determine which cell types were most transcriptionally and epigenetically impacted by aging across each of these groups/datasets. Rank ordering of computed AUC values for each comparison were collated to determine recurrence/conservation of high AUC values across comparisons. Values are displayed from top to bottom based on increasing rank product order across the 6 Augur analyses.

### Analysis of cell type proportion changes with aging across single nuclei RNA and ATAC libraries

Cell type proportions were calculated from the final Seurat object using the R package scProportionTest v0.0.0.9000^44^ in R v4.3.1 on both the snRNA-seq and snATAC-seq objects, to compare ages in a pairwise fashion, after splitting the objects for each sex and strain. This algorithm runs a permutation test on pairs of conditions for each cell type and returns the relative change in cell type abundance between groups with a confidence interval for each comparison. To avoid biases due to rare cell types, the proportion change analysis was only performed for cell types with ≥10 cells in each snRNA-seq or snATAC-seq libraries.

### Analysis of variance explained for gene expression and chromatin accessibility

For single nuclei RNA-seq and ATAC-seq datasets, the variance explained by experimental covariates was computed using the scater v1.28.0 package^158^. For bulk ATAC-seq analysis, this was calculated using the variancePartition v1.32.5 package^159^.

### Gene set activity scoring in snRNA-seq using UCell

Aging-relevant gene sets were collected for analysis, for both all cell types in the brain and for the microglia specifically. For brain-wide analysis, we used gene sets of: (i) genes regulated in at least 4 regions of the brain with aging in humans from GTEx^66^, (ii) common aging genes [CAG] from a mouse brain aging study^50^, and (iii) genes associated with cell senescence in mouse (SenMayo signature)^67^. For the microglia-specific analysis, we compiled gene sets of: (i) mouse microglia aging regulated genes (combined from 3 independent studies^75–77^), (ii) mouse gene signatures of microglia-activation states (*i.e.* terminally inflammatory microglia [TIM]^78^, damage-associated microglia [DAM]^75^, interferon-response microglia [IRM]^75^, and lipid-droplet-accumulating microglia [LDAM]^75^), and (iii) mouse macrophage M1/M2 polarization genes from *in vitro* cytokine-treated BMDMs^79,80^. In all cases, we use homologous genes as determined by BLAST alignments to the cognate species (see above) to obtain corresponding turquoise killifish gene sets for evaluation. Gene set activity levels were then computed using UCell v2.4.0^65^.

### Pseudobulking and differential gene expression analysis

We used muscat v1.14.0^52^ to pseudobulk the single-nuclei gene expression data by biological sample and cell type for downstream analysis. Only cell types with (i) well-resolved annotation (excluding the 4 Neuron_misc clusters) and (ii) ≥10 cells across all samples for both GRZ and ZMZ-1001 strains were used for downstream analyses to minimize technical noise and maximize interpretability. Analyses were conducted separately for samples from the GRZ and ZMZ-1001 strains. The package sva v3.48.0^160^ was used on the pseudobulked objects to estimate spurious noise linked to batch effects on gene expression, and limma v3.56.2 was used to regress these effects from the counts. DESeq2 v 1.40.2^161^ was used for differential gene expression analysis using the cleaned gene counts matrices from sva. Differential gene expression analysis was performed for genes and transposable elements (TEs). This approach was used to determine the number of differentially expressed genes in each cell type as a linear function of age (in weeks) using sex as a covariate (since the impact of sex on gene expression variance was minimal based on our analyses), to maximize the use of all ages. Genes with FDR <5% were considered significantly impacted by aging. This DEseq2-based analysis was used for all downstream analyses (*i.e.* functional enrichment, regulon activity, etc.).

### Pseudobulking and differential chromatin accessibility analysis

The pseudobulk-level analysis of our snATAC-seq was performed similarly to that our snRNA-seq (see above). Briefly, we used muscat v1.14.0^52^ to pseudobulk the single-nuclei chromatin accessibility profiles data by biological sample and cell type for downstream analysis, restricting analysis to cell types with (i) well-resolved annotation and (ii) ≥10 cells across all samples. We eliminated spurious technical noise effects as before using sva v3.48.0 and limma v3.56.2. Differential accessibility analysis was conducted using DESeq2 v1.40.2 on the cleaned peak accessibility matrices from sva. This approach was used to identify differentially accessible peaks in each cell type as a linear function of age (in weeks) using sex as a covariate. Due to the lower number of independent replicates, peaks with FDR <10% were considered significantly impacted by brain aging. This DEseq2-based analysis was used for relevant downstream analyses (*i.e.* functional enrichment).

### Reanalysis of brain H3K4me3 ChIP-seq data for comparison to bulk ATAC-seq

We obtained a previously generated H3K4me3 ChIP-seq dataset generated on whole-brain tissue isolated from adult male GRZ fish^22^ (accession number PRJNA259247). We used bowtie2 v2.5.1^155^ to map the ChIP and corresponding input reads to the GCF_001465895.1 turquoise killifish genome reference, using options “--sensitive –k 2 –p 1”. Next, we used MACS2 v2.2.7.1^156^ to call peaks using an effective genome size of 1.25e9, together with the “--broad” option. Peaks were annotated to closest genes using HOMER v4.11.1^162^.

### Bulk ATAC-seq differential accessibility analysis

We leveraged DiffBind v3.10.0 to generate peak/chromatin accessibility matrices from consensus MSPC peaks and mapped bam files. Analyses were conducted separately for samples from the GRZ and ZMZ-1001 strains. Final intervals calculated by DiffBind were annotated to the closest genes using HOMER v4.11.1^162^ together with the gene annotation gtf for the NCBI GCF_001465895.1 genome reference. Cleaned peak count matrices were then loaded into R. As above, spurious technical noise effects from experimental batches were removed using sva v3.48.0 and limma v3.56.2. We then performed differential accessibility analysis with DESeq2 v1.40.2. This approach was used to identify differentially accessible peaks in each strain as a linear function of age (in weeks) using sex as a covariate (since the impact of sex on peak accessibility variance was minimal). Peaks with FDR < 5% were considered significantly impacted by aging. This DEseq2-based analysis was used for relevant downstream analyses (*i.e.* functional enrichment).

Differentially accessible peaks were annotated using HOMER v4.11.1^162^, which enabled analysis of genomic distribution (*i.e*. promoter, intergenic, *etc.*). To determine overlap of differentially accessible ATAC-seq peaks with active or poised promoters in the brain, we overlapped DiffBind peaks with H3K4me3 peaks using bedtools v2.17.0^163^ intersectBed command with “-wao” option.

### HOMER motif density analysis along bulk ATAC-seq peaks

We provided HOMER v4.11.1^162^ with bulk ATAC-seq peaks with (i) increased chromatin accessibility with aging, (ii) decreased chromatin accessibility with aging, and (iii) all MSPC ATAC-seq peaks (*i.e*. background; see above), for both the GRZ and ZMZ-1001 datasets. To determine local density of nr3c1 canonical binding motifs around these peaks, we leveraged the annotatePeaks.pl script and the GRE (Glucocorticoid Response Element) motif file from the HOMER library (“gre.motif”; “GRE(NR),IR3/A549-GR-ChIP-Seq(GSE32465)/Homer”) in histogram mode with along 25bp bins centered around peak centers for a total of 1000bp. Motif density curves were then plotted in R using the output matrices from HOMER.

### Functional enrichment analysis for differential gene expression and chromatin accessibility

For gene set enrichment analysis, we obtained Gene Ontology [GO] terms for Zebrafish from ENSEMBL biomart v111 (accessed 2024-03-20) and converted them to turquoise killifish gene sets using our turquoise killifish/zebrafish homology table (see above). Importantly, terms with NAS (non-traceable author statement), TAS (traceable author statement), or ND (undetermined) evidence were excluded from analyses. In addition, we obtained mouse Reactome pathway definitions from MSigDB (v2023.2) and converted them to turquoise killifish gene sets using our turquoise killifish/mouse homology table (see above). In both the single nuclei and bulk ATAC-seq cases, peaks were assigned to the closest gene using the HOMER annotatePeaks.pl script, and data for the peak closest to the transcriptional start site [TSS] of the gene was extracted to represent that gene. Any gene for which the closest peak was >10kb away from the annotated TSS was not considered for analysis as association might be spurious.

To determine which pathways/gene sets were reliably and consistently enriched across ‘omic’ layers, we used multi-contrast pathway enrichment using the rank MANOVA (Multivariate Analysis of Variance) statistics, as implemented in the mitch v1.12.0^69^ R package, using our turquoise killifish GO and Reactome gene set definitions (see above). Gene sets with FDR < 5% were considered significantly regulated with aging across ‘omic’ datasets. For the single cell analysis, for each cell type, we considered snRNA-seq for each strain and snATAC-seq for GRZ, and the top 6 most significant terms with consistent directionality across ‘omic’ layers were reported in the figure. For the bulk ATAC-seq analysis, we considered regulation in each strain with aging, and the top 10 most significant terms with consistent directionality across strains were reported.

### RNAscope staining and imaging

To spatially verify our findings on both cell-type proportion and DE gene increases across age, killifish brains from both strains, GRZ (n=20) and ZMZ-1001 (n=30) were used as input for RNA-ISH using the RNAscope Multiplex Fluorescent Reagent Kit V2 kit for both 3-plex and 4-plex assays (Advanced Cell Diagnostics, 323110). The 4-plex assays also required the ancillary kit for the 4^th^ channel (C4; 323120). Brains were carefully dissected following euthanasia, fixed in 4% PFA for 24 hours, cryoprotected by a sucrose gradient (10, 20, and 30%) for 3 days, and then embedded in a Tissue-Tek® Cryomold® (REF#4566) with Tissue-Tek® OCT Compound (REF#4583). Brains are then stored in –80°C until being sectioned in a Leica, CM1860 cryostat at a 15mm thickness. Each slide consisted of 8-10 slices per brain and were pretreated following the fixed frozen sample preparation protocol per manufacturer instructions. Slides were air dried for 1hr at RT prior to running the assay to ensure optimal adhesion of the tissue. All remaining RNAscope procedures were done in accordance with the user manual provided by ACD (UM 323100). For the cell-type proportion assay, brain slices were hybridized to a combination of custom-designed probes complementary to *apoeb* (microglia marker; Nf-apoeb-C1; 1293241-C1), *mpz* (oligodendrocyte marker; Nf-mpz-C2: 1235131-C2), *s100b* (astrocyte marker; Nf-s100b-C3; 1235141-C3), and *map2* (neuron marker, Nf-map2-C4, 563391-C2). To develop the fluorescent signal, TSA Vivid Fluorophore 520 (Advanced Cell Diagnostics, 323272) was used to mark *apoeb*, TSA Vivid Fluorophore 570 (Advanced Cell Diagnostics, 323271) was used to mark *mpz*, Opal 620 (Akoya Biosciences, FP1495001KT) was used to mark *s100b*, and Opal 690 (Akoya Biosciences, FP1497001KT) was used to mark *map2*. For our first DE gene set, custom-designed probes were hybridized to zinc finger NFX1-type containing 1 (*znfx1;* Nf-znfx1-C1; 1556621-C1), helicase with zinc finger domain 2 (*helz2*; Nf-helz2-C2; 1556571-C2), and interferon induced with helicase C domain 1 (*ifih1*; Nf-ifih1-C3; 1556611-C3). The second DE gene set’s probes were hybridized to optineurin (*optn;* Nf-optn-C1; 1556611-C1), interferon regulatory factor 3 (*irf3*; Nf-irf3-C2; 1556631-C2), and signal transducer and activator of transcription 1a (*stat1a*; Nf-stat1a-C3; 1556581-C3). Both DE gene sets used TSA Vivid Fluorophore 570 for C1, Opal 620 for C2, and Opal 690 for C3. All fluorophores were diluted in TSA buffer (ACD, 322809) at a 1:1500 dilution. Negative controls for both Completed slides were preserved in ProLong Gold Antifade Mountant with DAPI (Thermo Fisher Scientific, P36931) and stored in 4°C until they were imaged. Each sample was imaged using a Leica Stellaris 5 confocal microscope (Leica microsystems) equipped with a DMI8 inverted microscope stand. The objectives used to both locate and image tissue were 10X dry objective lens (10x/0.4 HC PL APO CS2 506424) and a 63x oil-immersion lens (63x/1.4 HC PL APO OIL CS2 506350) lubricated with Leica Microsystems Type F immersion oil (11513859). Confocal z-stacks using a 0.3mm step size were taken for each by running Leica Application Suite X v4.4.0 (LAS X) software. Cell-type 4-plex images were taken with a 1024 x 1024 resolution at 600Hz (8000Hz for the mifepristone experiment), and DE gene sets were imaged with a 512 x 512 resolution using the resonant scanner (8000Hz) built into the Stellaris system. Laser intensities were 10 for all channels with an automatic digital gain of 2.5%. The laser mode was chosen based on the fluorophores’ excitation and emission spectra: DAPI for Channel 1, FITC for TSA Vivid fluorophore 520 (Channel 2), Cy3 for TSA Vivid fluorophore 570 (Channel 3), Texas Red for Opal 620 dye (Channel 4), and Cy5.5 for Opal 690 dye (Channel 5). Each image consisted of one hemisphere of the brain slice containing the optic tectum, which was anatomically guided by the African turquoise killifish coronal brain atlas established by D’Angelo (2013)^25^.

### Cell type abundance analysis by RNAscope

To measure cell proportion across age for each of the 4 cell types (microglia, oligodendrocytes, astrocytes/radial glia, and neurons), QuPath (Quantitave Pathology; QuPath-0.5.1-64x; Bankhead et al., 2017; https://qupath.github.io/) software was employed. For each image, three z-planes were chosen to perform cell segmentation on, and full image annotations were created. Cell detection parameters (pixel size of 0 to set it to the image default, background radius of 10mm^2^, sigma value of 1.5, minimum area of 10mm^2^, maximum area of 100mm^2^, threshold of 1) used a consistent value for each image in the experiment and had a cell expansion size of 3mm^2^. To measure cell proportion consistently across images of hemispheres of variable sizes, an artificial neural network (ANN) was built on training images for each experiment using manual single measurement classifiers for each of the four channels. Single measurement classifiers were then loaded onto the training image. Then, ‘classes’ for each cell type, including an ‘ignore/other’ class to account for cells that do not fall into one of the four categories (DAPI signals without any other channel colocalization) were created with manual annotation. Using the classes, the ANN was made and applied to all three z-planes for each image. Final measurements were the average of cell-specific counts divided by the total number of cells detected.

### Microglia soma volume determination using apoeb RNAscope staining

To make a 3-D reconstruction of the microglial soma and primary processes in cell-type images, the software Imaris (Imaris 11.0; Oxford Instruments; https://imaris.oxinst.com/) was used. The software’s ‘surface’ creation enabled the use of the *apoeb* signal from each 3-D image. Segmentation and creation parameters were as follows: Enable smoothing, surface grain size of 0.722mm (automatic), enable background elimination using the largest soma diameter of 17mm (chosen using the ruler tool on the ‘slice’ view. Detection thresholds used the automatic function. Detection thresholds for some images were manually adjusted due to variability in background detection to remove false positive detections. The created surfaces were then filtered by volume to remove surfaces that were smaller than 5mm^3^ and larger than 500mm^3^.

To determine whether microglia soma size showed overall changes with aging, each individual fish was represented by the median value of the reconstructed volumes of microglia. Due to strong batch-to-batch variation in absolute reconstructed volumes, that value was normalized to the median of all values from the cognate processing batch (2 batches for each strain). After the visual examination of boxplots revealed potential unique outlying values for specific biological groups, we performed a Grubbs test (using the ‘outliers’ v0.15 R package) to identify and exclude potential significant outliers.

### Differential gene expression analysis by RNAscope

Three representative z-planes were chosen for each image to capture the staining throughout the sample. To do this, full image annotations were created, and then using the DAPI channel intensity, cells were segmented using a watershed approach. Due to the lower resolution of the image compared to the cell-type images, new cell segmentation settings were made. The parameters were as follows: pixel size of 0 (defaults to the image pixel size), a background radius of 25, sigma value of 1.5, a minimum area of 5mm^2^, a maximum area of 100mm^2^, and cell expansion of 5mm^2^. This cell expansion size was chosen to fully capture subcellular detection for the mRNA puncta of interest. Thresholds for cell detection were 20 for 8-bit images (all GRZ images and the first cohort of ZMZ1001 images), and 2000 for 16-bit images (second cohort of ZMZ1001 images). These were based on the DAPI pixel value and blindly verified by another researcher. Following cell segmentation, subcellular spot detection was performed for channels 2, 3, and 4. The subcellular detection settings were as follows: Process all ‘Cells’, split by intensity and shape, spot minimum size was 0.5mm^2^, expected spot size was 1mm^2^, and the maximum spot size before being classified as a ‘cluster’ was 3mm^2^. Each channel threshold value was half of the automatic max channel intensity, and this was blindly verified by another researcher. After creating the subcellular detections in each of the three z-planes, the average value of the number of cells and number of intracellular spots for each channel combined with the number of clusters was recorded. Reported values were a ratio of the numbers of spots and clusters by the number of cells for each image. Analysis of the ZMZ1001 cohorts required the values to be normalized by the median value of each cohort to account for potential batch-to-batch variability.

### Classification analysis of microglia transcriptomes

Machine-learning classification was used to dissect age-related transcriptome shifts, their robustness and potential drivers in microglia. Specifically, we focused the training phase on microglia from the shorter lived GRZ strain as a baseline (profiled in batches pilot, cohorts 2, 3 and 4). This yielded 95 microglia from the pilot cohort, 505 from cohort 2, 735 from cohort 3 and 326 from cohort 4. Importantly, we decided to focus the training on classification models comparing young (6 weeks) to old (16 weeks) microglia, since classification models tend to have higher performance than regression models without deep coverage of the response variable (here, only 3 ages). These models can yield predictions in terms of “likelihood a microglia nucleus is derived from an old animal”, with middle-aged animals expected to have intermediate likelihoods of old age. This strategy was used successfully on machine-learning models trained on mouse brain aging snRNA-seq^83^.

To limit training issues due to imbalanced representation of sex and age groups, we randomly sampled 125 cells from young females, young males, old females and old males for a total of 500 training cells. This left a remainder of 1161 microglia nuclei from GRZ fish withheld from training (44 young, 392 middle-aged and 725 old). From the training data only, we identified 826 <u>genes</u> that (i) were among the top 3,000 most variable genes the data subset, and (ii) had an average expression across cells of > 0.25 UMIs in all 4 biological groups in our training set, to be used as features in our models. We then binarized scaled gene expression data as proposed with the CellBiAge framework^83^ to improve model performance. We also included sex as a potentially important covariate. Together, binarized gene expression and sex were used as training features.

To train classification models, we used the R package ‘caret’ v6.0-94. Random forest (RF) and elasticNet (eNET) were based on caret’s usage of packages ‘randomForest’ v4.7-1.1 and ‘glmnet’ v4.1-8, respectively. For both model types, training was performed using 10-fold cross-validation to search a parameter grid, and the models were trained to maximize the AUROC metric. Out-of-bag predictions for the final tuned models across the 10 training folds are reported in **Extended Data Figure 12b**.

We used withheld GRZ young and old microglia nuclei to determine testing accuracy. The caret ‘predict’ function was used to predict likelihood of old age using the trained models on all withheld data and microglia nuclei from ZMZ-1001 individuals.

Variable importance scores were extracted by caret for the RF model (‘MeanDecreaseGini’). For the eNET model, we used the absolute value of the final regularized coefficients (coef(my.enet.fit$finalModel, s=my.enet.fit$finalModel$lambdaOpt)) as feature importance metrics. A ‘consensus’ ranking of top contributing variables was computed using the rank products of variable importance across models to determine the top 15 predictive features.

### Transcription Factor [TF] regulon activity estimation in snRNA-seq

To infer TF regulon activity, we used our pseudobulked DESeq2 cell-type specific aging differential gene expression results in both GRZ and ZMZ-1001 strains, together with decoupleR v2.6.0^87^. We also used the mouse CollecTRI curated collection of TF regulons^164^, accessed through OmnipathR v3.8.2, together with our BLAST-based turquoise killifish-mouse homology database (see above) to generate a putative turquoise killifish TF regulon collection. To score potential differential TF activity from this collection, we used the ‘fgsea’ scoring algorithm for the decoupleR algorithm together with the DEseq2 Wald statistic. TF regulons with p < 0.05 were considered significantly differentially impacted with aging, and those with p < 0.1 were considered to have a suggestive trend for differential impact. All significant and suggestive results are reported in **Extended Data Table 8a,b**.

To identify recurrently impacted candidate TF regulons, we identified significantly impacted TF regulon with aging in each cell type and strain. We first applied a filter to identify TFs with significant regulation in 4 or more cell types in either the GRZ or the ZMZ-1001 strain, identifying 7 and 14 age-regulated candidate TF regulons, respectively. Only one of these, stat1, was directly overlapping between both strains. Data from both strains was then merged, and a secondary filter was applied to that list to retain only TFs with significance in at ≥1 cell types in the strain they were not initially identified in. A tertiary filter was applied to focus on TFs with consistent predicted direction of activity change within each strain, which yielded a list of 12 candidate TF regulons. For plotting purposes, we included decoupleR results for these 12 TFs, adding in cells with lighter color-shading to reflect additional cell types with suggestive trends (p < 0.1), to visualize additional subthreshold evidentiary support for broad cross-cell type TF activity misregulation with aging.

### Differential DNA-binding motif accessibility in snATAC-seq

To determine whether our snATAC-seq data showed evidence of differential TF activity, we leveraged the “filtered_tf_bc_matrix” output folder from cellranger-atac, which contains accessibility scores for JASPAR2018 DNA binding motifs (579 scored motifs). This data was imported into R and added as an additional assay in our annotated snATAC-seq Signac object. Then, we performed differential accessibility analysis using Seurat FindMarkers() between cells from the young and old age groups (regardless of sex, since we saw little effect of sex in our previous analyses, thus affording us greater statistical power in this analysis), using options “only.pos = F, min.pct = 0.05, logfc.threshold = 0” to output scores for all motifs. This analysis was restricted to well annotated cell types with ≥10 cells per library. Motifs with FDR < 5% in all 9 tested cell types (23 motifs) were designated as recurrently misregulated with aging, and the 10 motifs with the highest average log_2_(Fold change in accessibility with age) were selected for plotting. All significant results are reported in **Extended Data Table 8c**.

### TF motif enrichment in differentially accessible bulk ATAC-seq peaks

Bulk ATAC-seq peaks identified as differentially accessible by our DESeq2 analysis in either of the turquoise killifish strains with aging (see above) were used as input for motif finding using the findMotifsGenome.pl script from HOMER, together with the cognate background for enrichment (i.e. all ATAC-seq peaks identified in this strain), using options “-size given –mset vertebrates” (enabling the use of the HOMER motif library). Enrichment of known vertebrate motifs from the knownResults.txt output was then loaded into R for parsing and plotting. Motifs were considered to be significantly enriched with HOMER FDR < 5%. The enrichment score was determined by dividing the percentage of differential peaks with the motif by the percentage observed in the cognate background, with a positive sign for motifs identified in peaks with increased accessibility with aging and a negative sign for motifs identified in peaks with decreased accessibility with aging. We then filtered the data to identify motifs significantly enriched in the same direction in both turquoise killifish strains, yielding 79 motifs enriched in regions with increased accessibility and 96 motifs enriched in regions with decreased accessibility. We then eliminated TFs whose motifs where enriched both in regions with increased and decreased accessibility, as these results are difficult to interpret. The top 5 most enriched motifs (as determined by rank product) in each direction were then selected for plotting. All significant results are reported in **Extended Data Table 8e,f.**

The HOMER annotatePeaks.pl script was used to generate histogram plots of candidate TF motif density at differentially regulated and background peaks with a 25bp resolution and a 1kb plotting window.

### Differential footprint accessibility analysis using TOBIAS for bulk and snATAC-seq

We leveraged TOBIAS v0.12.12^94^ to identify potentially differentially accessible TF footprints in our bulk ATAC-seq and snATAC-seq (stratified by cell type) datasets.

For the snATAC-seq dataset, we first extracted barcode/cell type information from the annotated Signac snATAC-seq object. We then used the subset-bam v1.1.0 program from 10xGenomics (https://github.com/10XGenomics/subset-bam) and the cellranger-atac output bam files to generate cell-type specific bam files for our snATAC-seq dataset. To achieve the necessary sequencing depth for TOBIAS, we then merged reads per age group (pooled sex) for each cell type. We restricted footprint analysis to the same genomic intervals as the bulk ATAC-seq analysis (due to more robust peak calling conditions in the bulk dataset). The remainder of the TOBIAS pipeline was run as described above in each cell type separately, comparing young vs. old cells. We reported the cell type-wise results for the NR3C1_MA0113.3 motif. All significant results are reported in **Extended Data Table 8d**.

For the bulk ATAC-seq analysis, we performed the analysis comparing data from young vs. old GRZ brains, young vs. old ZMZ-1001 brains, and young vs. geriatric ZMZ-1001 brains, one comparison for each sex (used as independent probing of differential TF activity). Importantly, since TOBIAS (i) does not handle biological replicates and (ii) benefits from deeper sequencing depth, aligned reads were pooled across libraries to generate a single bam file for each condition to be compared. To reduce technical noise, footprint analysis was restricted to a merged ned file generated from the significantly differentially accessible ATAC-seq peaks with aging from the GRZ and ZMZ-1001 strain obtained using mergeBed from bedtools v2.17.0. Aligned reads were then processed as recommended by TOBIAS developers. Briefly, files were subjected Tn5 bias correction with TOBIAS ATACorrect, scoring with TOBIAS FootprintScores, TOBIAS BINDetect and the JASPAR2022 CORE non-redundant motifs. Motifs were considered to be significantly differentially accessible with age if TOBIAS FDR <5%. Enrichment was ranked in each comparison based on the “change” statistic, and the top 5 motifs from the rank product across the 6 comparisons from the top or bottom of the list were selected for plotting. All significant results are reported in **Extended Data Table 8g-i**.

### Analysis of candidate TF activity change with aging in mouse and human datasets

To determine whether predicted age-related changes in TF regulon activities identified using decoupleR across cell types in the aging turquoise killifish brain were also observed with mouse and/or human brain aging, we took advantage of publicly available snRNA-seq and bulk RNA-seq datasets (see **Extended Data Table 10b** for dataset-level information).

For the snRNAseq datasets, some datasets were provided as annotated by the authors (Mouse: whole brain^41^, hypothalamus^76^, and subventricular zone [SVZ]^82^; human: hippocampus^92^, and dorsolateral prefrontal cortex^93^), while annotation was unavailable for others (Mouse: hippocampus^88^, caudate putamen & hippocampus^89^, prefrontal cortex^90^). For datasets provided as feature/barcode outputs without cell type annotation, we used a standardized Seurat processing pipeline similar to the one we used for our turquoise killifish dataset. Briefly, cells were quality with parameters subset = nFeature_RNA > 250, nFeature_RNA < 5000, percent.mito < 10, nCount_RNA < 25000, decontX_contamination < 0.25, and multiplets were annotated and removed using the union of intersection of singlet calls from Doubletfinder and scds for each individual library. Cell types present in the unannotated mouse datasets were then annotated using singleCellNet v0.1.0^165^ and the transcriptional profiles from a published hypothalamus snRNA-seq object as a reference^76^. For all snRNA-seq datasets, differential gene expression analysis with aging was then performed for cell types with ≥10 cells in all samples of a dataset using muscat pseudobulking, removal of unknown technical noise using sva/limma, and differential gene expression testing with DESeq2 using age as a linear variable of interest (and sex as a covariate when metadata was available and non-uniform across samples). DEseq2 results were then used for TF regulon activity analysis with decoupleR.

For bulk RNA-seq profiling of aging human brain regions from the GTEx consortium^91^, we downloaded gene count matrices from 13 brain regions from GTEx v10 open access data [https://www.gtexportal.org/home/downloads/adult-gtex/bulk_tissue_expression]. To be able to take advantage of the full age range of available samples with a linear analysis, the age in years for each sample was imputed to the middle of the provided age-ranges (*e.g*. 25 years for the 20-30 years bracket). We then performed linear differential gene expression analysis with aging using DESeq2 on sva-corrected counts using sex as a covariate. DEseq2 results were saved for TF regulon activity analysis with decoupleR.

For bulk RNA-seq profiling of aging mouse brain regions, we obtained published paired end fastq raw sequencing data for a dataset with cerebellum and olfactory bulk tissue with aging^166^, a dataset with 13 brain regions with aging^50^, and a dataset with hippocampus bulk tissue with aging^167^. Paired reads were mapped to the mm39 mouse genome reference using STAR 2.7.7a^168^ with options “--outFilterScoreMinOverLread 0.4 –-outFilterMatchNminOverLread 0.4 –-readFilesCommand gzcat –-runThreadN 1 –-outFilterMultimapNmax 50 –-outFilterIntronMotifs RemoveNoncanonicalUnannotated –-outSAMtype BAM SortedByCoordinate –alignEndsProtrude 15 ConcordantPair”. Read counts were assigned to genes from the mm39 gene models using featureCounts from subread 1.5.3^169^, using options “-t exon –D 1500 –p –-primary –T 1 –s 0”. Count tables were imported into R 4.3.1 for downstream processing. Differential gene expression analysis with aging was then performed for each brain region with our standard pipeline, including removal of unknown technical noise using sva/limma, and differential gene expression testing with DESeq2 using age as a linear variable of interest. DEseq2 results were then used for TF regulon activity analysis with decoupleR.

DecoupleR was run (as above) using the CollecTRI regulon data for the cognate species (*i.e*. mouse for mouse datasets, human for human datasets). Predicted TF regulon activity for homologs of recurrent age-regulated regulons in the brain of turquoise killifish were extracted for plotting as above.

### Measurement of cortisol in turquoise killifish kidney tissue

Cortisol was measured at the Endocrine Technologies Core at Oregon National Primate Research Center using liquid chromatography-tandem triple quadrupole mass spectrometry (LC-MS/MS) on a Shimadzu Nexera-LCMS 8050 instrument (Shimadzu, Kyoto, Japan), using a previously established method, with modifications^170^. Briefly, kidney samples were weighed and loaded into 12 x 75mm glass tubes containing cortisol-d4 internal standard in 350µL ultrapure water (Milli-Q, EMD Millipore, Billerica, MA). Samples were briefly homogenized after addition of 500µL methanol, then 125µL of each homogenate was then mixed with 125µL ultrapure water and transferred to a 96-well SLE plate (Biotage, Uppsala, Sweden). Steroids were eluted with 3 x 600µL dichloromethane (Sigma, St. Louis, MO) into plates containing 100µL isopropanol (Sigma), dried under forced air and reconstituted in 50µL of 25:75 methanol:ultrapure water. Then, 15µL of each sample were injected into the LC-MS/MS system for analysis as previously described^170^. Excellent linearity was observed for cortisol within the range of the calibration curve (R>0.999). The assay range was 0.091-200ng/mL, which translated to a quantifiable range of 0.015 – 38.09ng/sample. The lower limit of quantification was 4.1pg on the column. Intra– and inter-assay coefficients of variation were <5%. Conditions with potential outliers were visually identified, and significant outliers were excluded if the Rosner test for outliers (as implemented in R package ‘EnvStats’ v3.0.0^171^) was significant with p < 0.05.

### Mifepristone intervention experiment

#### Preparation of a mifepristone stock solution

A 50 mg/mL stock solution of mifepristone was prepared by dissolving 100 mg of mifepristone powder (Sigma-Aldrich, Cat# M8046; stored at 4L°C) in 2 mL of 100% molecular biology-grade ethanol. Ethanol was first aliquoted into a sterile 15LmL conical centrifuge tube. The mifepristone powder was carefully weighed and transferred into the ethanol, ensuring complete transfer of the compound._The conical tube was securely capped and vortexed vigorously until the solution became a uniform, clear champagne color with no visible particulates or clumps. No heat was applied during the dissolution process._The resulting solution was sterile-filtered using a 0.22μm filter. The final sterile-filtered mifepristone stock solution (50Lmg/mL) was stored at –20 °C for long-term use.

#### 1% Agar gel and experimental drug food prep

Food grade agar-agar powder (Hznxolrc) was mixed with filtered sterilized MilliQ water in a microwave-safe container and heated in a microwave oven until fully dissolved to form the 1% mixture, then allowed to cool to ∼45–50 °C. For each fish, 1/8 tablespoon (i.e., the amount an adult fish would have for one meal) of freeze-dried bloodworms (Hikari) was placed on a clean petri dish, and an equal volume (∼1/8 tablespoon) of the warm agar solution was poured over the food to coat it. The mixture was gently mixed in this semi-fluid state. Immediately afterward, 0.6μL of the experimental drug was pipetted directly into the warm agar-food mixture (corresponding to 30μg of mifepristone). For the control food, we added the same amount of pure ethanol (the vehicle for mifepristone). The portion was gently mixed to ensure even distribution of the drug/pure ethanol while the agar was still semi-fluid. The mixture was then left at room temperature to cool and solidify, forming a gel-sealed food portion. Once solidified and at room temperature, the food was broken into smaller pieces to facilitate ingestion by the fish and was fed immediately. All preparations were made fresh daily and used immediately without storage. The chosen daily dosage is consistent with well-tolerated dosage in rodents (12.5-50mg.kg^-1^.day^-1^)^172,173^.

#### Mifepristone supplementation experiment

Turquoise killifish were allowed to hatch and age without intervention on our recirculating system until they reached 10 weeks of age. Then, they were moved to a non-recirculating flow-through shelf of the system to avoid any inadvertent recirculation of drug byproduct. Fish were randomly assigned to the control or mifepristone treatment groups. From that point, one of the daily meals was replaced using the agar-coated food (see above), and fed to single-housed animals using tweezers^23^, guaranteeing full consumption of the meal and limited contamination of the tank water. For young control fish, 5-6-week-old fish were moved to the flow-through shelf and fed the control diet in the same way as above to match dietary and water-quality parameters of the aged group as closely as possible. Fish were euthanized at 2-5pm in a snaking order to minimize batch and circadian effects.

### Bulk RNA-seq of turquoise killifish brain tissue

For RNA extraction of tissues from our Mifepristone intervention cohorts, each sample was transferred to a Lysing Matrix D tube (MP Biomedicals 6913500) with 1 mL of TRIzol (Invitrogen #15596018) and homogenized using a Beadbug 6 microtube homogenizer (Benchmark Scientific D1036) at 3,500 rpm for 30 seconds for 3 cycles. RNA was isolated according to the manufacturer instructions, with the addition of 1µL Glycoblue reagent (Thermo Fisher Scientific AM9516) to improve RNA pelleting and recovery. Purified total RNA samples were sent to the Novogene corporation for bulk unstranded mRNA-sequencing. Briefly, the company performed mRNA selection using polyA capture, RNA fragmentation, cDNA reverse transcription and then unstranded RNA sequencing library construction. RNA libraries were sequenced as paired-end 150bp reads on the Illumina Novaseq X platform. Raw FASTQ sequencing reads were deposited to SRA under Bioproject accession PRJNA1298850.

### Analysis of turquoise killifish brain tissue bulk RNA-seq

Paired end 150bp FASTQ reads were hard-trimmed for their first 9bp at the 5’ end to eliminate RNA reverse transcription priming biases^174^. This was done using Trimgalore 0.6.6 (github.com/FelixKrueger/TrimGalore). Stringency filters were added to eliminate any bases with phred score < 15 (corresponding to < 1% of bases). The first 9 bases of each read were hard-trimmed to remove known 5’ biases in RNA-seq libraries due to random RT priming^174^. Trimmed reads were mapped to the GCF_001465895.1 turquoise killifish genome reference (with the FishTEDB hard-masked repeats and addition of repeat contigs, as above) using STAR 2.7.7a^168^ with options “--outFilterScoreMinOverLread 0.4 –-outFilterMatchNminOverLread 0.4 –-readFilesCommand gzcat –-runThreadN 1 –-outFilterMultimapNmax 50 –-outFilterIntronMotifs RemoveNoncanonicalUnannotated –-outSAMtype BAM SortedByCoordinate –alignEndsProtrude 15 ConcordantPair”. Read counts were assigned to genes from the NCBI GCF_001465895.1 reference gene models using featureCounts from subread 1.5.3^169^, using options “-t exon –D 1500 – p –M –-fraction –T 3 –s 0”. Read count tables were imported into R 4.3.1 for downstream processing. Since the male and female cohorts were obtained at different times, they were analyzed separately to avoid confounding biological and batch effects. Since our previous analyses showed minimal impact of sex on brain aging, the female and male cohorts were just treated as independent cohorts for the purposes of our analyses.

The processing of our bulk RNA-seq in R was performed using the same basic pipeline as for our pseudobulk analyses described above for consistency. To note, since partial counts could be assigned from multimapping reads (due to the use of the –M –-fraction option with featureCounts), count tables were rounded before further processing. Next, counts were subjected to technical denoising using sva v3.48.0^160^ and limma v3.56.2 as before, and DESeq2 v1.40.2^161^ was used to perform differential gene expression analysis using cleaned counts matrices. Differential gene expression analysis was performed for genes and transposable elements (TEs). Differentially expressed genes and TEs were analyzed after joint normalization in DESeq2 in 2 pairwise comparisons: (i) young vs. old controls (*i.e*. effect of baseline brain aging), and (ii) old control vs. old mifepristone supplemented (*i.e*. effect of supplementation on the aged brain transcriptome). Genes and TEs with FDR <5% were considered significantly impacted by aging or mifepristone. These results were used for downstream analyses (*i.e.* functional enrichment, regulon activity, deconvolution, etc.). DESeq2 results are reported in **Extended Data Table 9a-d**. Multidimensional scaling was performed using 1 – Rho as the distance metric using the base R ‘cmdscale’ function.

### Gene Set Enrichment Analysis of brain aging genes in response to mifespristone treatment

We used the Gene Set Enrichment Analysis (GSEA)^175^ paradigm through the ‘phenotest’ v1.48.0 R package implementation. *Ad hoc* gene sets related to brain aging in the female/cohort 1 and male/cohort 2 RNA-seq datasets were obtained by using genes called as significantly regulated by DEseq2 at FDR < 5%, split in upregulated and downregulated with aging gene sets based on the size of the log_2_(Fold change in expression with aging) calculated by DESeq2 (see above). GSEA significance was calculated using 10,000 permutations.

### Functional enrichment analysis for bulk brain RNA-seq with aging and in response to mifepristone

Similar to before, we used multi-contrast pathway enrichment using the rank MANOVA s statistics from mitch v1.12.0^69^ together with the same GO and Reactome gene set definitions. Gene sets with FDR < 5% were considered significantly regulated. We report the top 6 most significant terms with consistent directionality across our cohorts (*i.e.* consistent directionality in aging male vs. female fish AND consistent directionality in mifepristone-treated male vs. female fish, with no constraint on agreement or disagreement between the aging and mifepristone comparisons). All significant enrichment results are reported in **Extended Data Table 9g,h**.

### Meta-analysis of bulk brain RNA-seq cohorts

We decided to leverage the full power of our 2 independent bulk brain RNA-seq experiments with aging and mid-life-initiated mifepristone treatment (1 female cohort, 1 male cohort) using a meta-analysis statistical framework. For this purpose, we used results from the DESeq2 differential gene expression analyses for the female and male cohorts (see above), together with R package ‘metaRNAseq’ v1.0.8^176^. Importantly, we considered genes to be differentially regulated as part of the meta-analysis if: (i) the FDR from inverse normal p-value combination was < 5% (since the inverse normal combination yielded more conservative gene lists than the Fisher method in our hands), and (ii) the sign of regulation of gene expression was the same in both datasets. Meta-analysis was performed in parallel for the effect of aging and the effect of mifepristone. Gene-level results for meta-analyses are reported in **Extended Data Table 9e,f**.

For genes exhibiting a reversal of the aging trend upon mifepristone treatment in the meta-analysis, we performed functional enrichment with the same GO and Reactome gene set definitions as before, using an overrepresentation analysis paradigm through R package ‘clusterProfiler’ v4.8.2^177^. All significantly enriched gene sets are reported in **Extended Data Table 9i,j**.

### Deconvolution-based estimation of microglia proportion in bulk brain RNA-seq datasets

Deconvolution of our bulk brain RNA-seq was done using R package CSCDRNA v1.0.3^171^, with parameters “min.p = 0.3; markers = NULL”, and using the our annotated snRNA-seq atlas restricted to nuclei from GRZ fish (matching the strain of our bulk RNA-seq dataset). Predicted proportions of microglia in our samples were combined across sexes/cohorts to improve our statistical power, and the potential impact of aging and mifepristone treatment was tested for significance using a Wilcoxon rank-sum test.

### Statistics and reproducibility

All statistical and bioinformatics analyses were conducted using R v4.3.1^178^. All statistical analyses were performed as specified in figure legends and in the corresponding section of the methods. We used nonparametric statistical tests whenever possible to avoid assuming normality/homoscedasticity of data distributions. For analysis of variance, we applied parametric ANOVA when a Shapiro-Wilkes test yielded non-significant results (p > 0.05) on ANOVA residuals, consistent with data normality. When ANOVA residual distribution was incompatible with the normality assumption, non-parametric ART-ANOVA [Aligned Rank Transform ANOVA]^179^, which doesn’t require data normality, was performed instead with R package ‘ARTool’ v0.11.2. No statistical methods were used to predetermine *a priori* sample size, since effect sizes and variance could not be known prior to our experiments in this exploratory study. When relevant, the Grubbs test (using the ‘outliers’ v0.15 R package) or Rosner test (using the ‘EnvStats’ v3.0.0^171^ R package) were used to detect and exclude significant outliers (with p < 0.05), as indicated in relevant figure legends. Replicate samples were always obtained from independent animals to sample true biological variation. For each experiment, animals were processed in an alternating/snaking order rather than homogeneous stretches to minimize group, batch and circadian effects. For observational aging studies, no randomization was possible as groups were biologically determined. For the Mifepristone intervention paradigm, 10-week-old fish were randomized to the treatment *vs*. control group when initiating treatment. This process was performed independently for each Mifepristone treatment cohort.

Blinding was not relevant to studies conducted with automated systems-mediated data collected (*i.e*. sequencing-based data, LC-MS), which cannot be influenced by subjectivity from the experimenter. For RNAscope analysis, analyses were performed using standardized batch QuPath and Imaris workflows, methods which are condition-agnostic. For analyses requiring manual thresholding in the initial cell segmentation and/or gene expression quantification by RNAscope, analyses were validated by blinded observers.

## Code Availability

Scripts for this study were run using R v4.3.1 or 4.3.3.

Scripts used to analyze ‘omic’ datasets and perform statistical analyses are available on the Benayoun lab GitHub at https://github.com/BenayounLaboratory/Killifish_Brain_Atlas.

## Data Availability

The raw FASTQ files have been deposited to the Sequence Read Archive under accession PRJNA1121423 (whole genome sequencing), PRJNA1105049 (snRNA-seq), PRJNA1100604 (Bulk ATAC-seq), PRJNA1106410 (snATAC-seq) and PRJNA1298850 (bulk RNAseq). The final annotated Seurat object for the snRNA-seq analysis has been deposited to Figshare (Doi: 10.6084/m9.figshare.31847155) and used to build an interactive R shiny application for ease of search (https://minhooki.shinyapps.io/killifish-brain-atlas/). Compressed TIFF of the RNAscope microscopy pictures have been deposited to Figshare (https://doi.org/10.6084/m9.figshare.31931247).

## Acknowledgments

We thank Delana Obenauer, Brandon Cho, Jomille Jerez and Isabel Ollerton for assistance with killifish husbandry. We are grateful to feedback from Mr. Clayton Baker and Ms. Shiqi (Skye) Chen for their insights during the preparation of this manuscript. We thank Dr. Dario Valenzano for kindly sharing the ZMZ-1001 turquoise killifish strain with our laboratory.

This work was supported by NIA T32 AG052374 Postdoctoral Training Grant fellowship to B.B.T; Shenoy Undergraduate Research Fellowship in Neuroscience (SURFiN) award SFI-AN-SURFiN-00014329 to G.M.C.; Simons Collaboration on Plasticity in the Aging Brain grant SF811217 and SFI-AN-NC-AB-Research-00018042 to B.A.B. The authors acknowledge the Center for Advanced Research Computing (CARC) at the University of Southern California for providing computing resources that contributed to the results reported within this publication (https://carc.usc.edu). We thank Dr. David Erikson and the Endocrine Technologies Core (ETC) from the Oregon National Primate Research Center (ONPRC) for performing the quantification of cortisol levels from African turquoise killifish. The Endocrine Technologies Core (ETC) at Oregon National Primate Research Center (ONPRC) is supported (in part) by NIH grant P51 OD011092 for operation of the Oregon National Primate Research Center. The funders had no role in the design and reporting of the research.

Some panels were created using graphical elements from the NIAID bioart resource [https://bioart.niaid.nih.gov], bioicons [https://bioicons.com/] and scidraw [https://scidraw.io/]. We thank Diogo Losch De Oliveira John Chilton, and Servier for specific icons used in this manuscript [https://zenodo.org/records/3925953; doi.org/10.5281/zenodo.3926033; https://smart.servier.com/].

## Author contributions

B.A.B. designed the study. H.Z., and A.A. performed fish husbandry, and dissected killifish tissues for experiments. H.Z. optimized and executed mid-life-initiated delivery of mifepristone in killifish. B.B.T. and A.A. isolated brain nuclei. B.B.T. and B.A.B. constructed 10xGenomics libraries. R.G.W., B.B.T., R.B., and G.M.C. performed brain *in-situ* hybridization with RNAscope. R.G.W. performed all analyses of RNAscope images, with help from A.K. A.J.J.L. and E.H.L. performed tissue RNA extractions for bulk RNA-seq. B.B.T., A.J.J.L., M.K., S.A.M., and B.A.B. performed computational analyses. J.T. helped design and interpret mifepristone supplementation experiments. R.G.W., A.J.J.L. and B.A.B. wrote the manuscript with input from all authors. All authors edited and commented on the final manuscript.

## Legends to Extended Data Figures

**Extended Data Figure 1:** Genetic architecture of African turquoise killifish lab strains. (**a**) Experimental scheme for genetic variation mapping of the GRZ vs. ZMZ-1001 strains used in this study and hosted at USC by whole-genome shotgun [WGS] sequencing. For each strain, genomic DNA from 2 male individuals were used to map genetic, since males are the heterogametic sex in this species^19,38^. (**b**) Plink PCA of laboratory African turquoise killifish lab isolates, annotated by observed captive lifespan, tail color and hosting institution. Note genetic proximity of yellow-tailed vs. red-tailed isolates, regardless of captive lifespan. (**c**) Analysis of shared genetic variation in USC-hosted GRZ and ZMZ-1001 male killifish. Only SNPs consistently detected in both individuals of the same strain are considered in this analysis. Consistent with its use for assembly of the species reference genome, less SNPs are observed in GRZ animals. ZMZ-1001 fish show ∼610K unique genetic variants compared to the GRZ fish.

**Extended Data Figure 2:** Quality control analysis of African turquoise killifish snRNA-seq brain aging dataset. (**a-d**) UMAP plots of African turquoise killifish snRNA-seq brain aging atlas colored by (**a**) strain of origin, (**b**) sex, (**c**) age group and (**d**) data generation batches. (**e**) Dotplot of marker gene expression, showing the average expression level and percentage of cells expressing each marker gene in the final annotation. Overall markers of mature neurons (*rbfox3*, *map2*, *ncam1*) are provided on the rightmost side of the plot, to illustrate shared neuronal identity of annotated neuronal nuclei cohorts. Since many NCBI gene names were uninformative, names corresponding to their predicted homologs are used. Original NCBI gene names are provided in **Extended Data Table 1d**. OPCs: Oligodendrocyte Progenitor Cells; NSPCs: Neural Stem and Progenitor Cells. See **Extended Data Table 1d** for NCBI gene name correspondence.

**Extended Data Figure 3:** Quality control analysis of African turquoise killifish snATAC-seq brain aging dataset. (**a-c**) UMAP plots of African turquoise killifish snATAC-seq brain aging atlas colored by (**a**) sex, (**b**) age group and (**c**) data generation batches. Note imperfect batch correction was achieved at the UMAP level for the granule excitatory neurons and oligodendrocytes nuclei clusters. (**d**) Dotplot of marker gene activity (measured as promoter snATAC-seq accessibility), showing the average activity level and percentage of cells with active marker gene in the final annotation. Overall markers of mature neurons (*map2*, *ncam1*) are provided on the rightmost side of the plot, to illustrate shared neuronal identity of annotated neuronal nuclei cohorts. Since many NCBI gene names were uninformative, names corresponding to their predicted homologs are used. Original NCBI gene names are provided in **Extended Data Table 1d**. OPCs: Oligodendrocyte Progenitor Cells; NSPCs: Neural Stem and Progenitor Cells. See **Extended Data Table 1d** for NCBI gene name correspondence.

**Extended Data Figure 4:** Augur cell type prioritization analysis across snRNA-seq and snATAC-seq datasets. (**a**) Heatmap of Augur AUC rank in each data subset, (i) GRZ female aging snRNA-seq, (ii) GRZ male aging snRNA-seq, (iii) ZMZ-1001 female aging snRNA-seq, (iv) ZMZ-1001 male aging snRNA-seq, (v) GRZ female aging snATAC-seq, (vi) GRZ male aging snATAC-seq. Note consistent high AUC scores across groups for OPCs, oligodendrocytes and microglia, suggesting their ‘omic’ profiles are most responsive to aging.

**Extended Data Figure 5:** Validation of RNAscope staining for overall cell types. **(a)** Representative maximum z-projected images using 4-plex RNAscope for the ZMZ-1001 conditions’ cell-type markers: *apoeb* in yellow*, mpz* in green*, s100b* in magenta*, map2* in red and DAPI is in blue. Young and geriatric (not middle/old) brain images from both sexes are shown. Scale bars are 250mm in length.

**Extended Data Figure 6:** Differential gene expression analysis with aging in the African turquoise killifish brain. (**a**) Strip plots of differentially expressed FishTEDB transposable elements [TE] per cell type (FDR < 5%) in aging GRZ (left) and ZMZ-1001 (right) brains with aging (sex was used as a covariate). Differentially expressed transcripts are highlighted while non-significant genes are shown in gray. Transcripts above the midline are upregulated with aging while transcripts below the midline downregulated with aging. (**b**) Overlap analysis of genes upregulated and downregulated with aging across cell types in the GRZ vs. ZMZ-1001 strains. Genes upregulated at FDR < 5% with aging in the GRZ vs. ZMZ-1001 strains were compared. Significance of overlaps according to Fisher’s exact test are reported below Venn diagrams. Note strong sharing of upregulated genes, and generally weaker sharing of downregulated genes. (**c**) Spearman rank correlation Rho of gene expression changes per cell type between GRZ and ZMZ-1001 datasets, showing significant rank agreement for gene and TE regulation with aging in all cel types (p < 0.05). (**d**) Heatmaps of Z-scored median pseudobulked gene expression with aging of top shared genes upregulated across cell types with aging from Fig. 3d, showing robust upregulation patterns even in cell types where changes did not reach statistical significance.

**Extended Data Figure 7:** Validation of RNAscope staining for top shared aging differentially expressed genes. Representative maximum z-projected images of differentially expressed gene set validation across age using two different 3-plex RNAscope assays for spatial validation in a geriatric female ZMZ-1001 brain. All channels for mRNA targets of interest were split for this representation. **(a)** Set 1 shows *znfx1* in green*, helz2* in yellow, and *ifih1* in red. **(b)** Set 2 shows *optn* in green*, irf3* in yellow, and *stat1a* in red. Each image possesses an overlay with the DAPI channel. Scale bars are 30mm in length.

**Extended Data Figure 8:** Differential chromatin accessibility analysis with aging across cell types in the African turquoise killifish brain. **(a)** Scheme of differential chromatin accessibility analysis for snATAC-seq. Explained chromatin accessibility variance was estimated using scater. Cell types were partitioned in each strain according to their annotation in our snATAC-seq atlas and analyzed for differential regulation using pseudobulking with muscat. Differential chromatin accessibility analysis was performed with DESeq2. To maximize robustness and interpretability of pseudobulk analyses, only cell types with (i) ≥ 10 nuclei per snRNA-seq library and (ii) well-defined cell identity were analyzed. (**b**) Density plot of explained chromatin accessibility variance in snATAC-seq dataset as a function of cell type, age, and sex. Notice that aging is the largest contributor to chromatin accessibility variance after cell type. (**c**) Strip plots of differentially accessible peaks per cell type (FDR < 5%) in aging GRZ brains (sex was used as a covariate). Differentially accessible peaks are highlighted while non-significant peaks are shown in gray. Peaks above the midline become more accessible with aging while peaks below the midline become less accessible with aging.

**Extended Data Figure 9:** Differential chromatin accessibility analysis with aging in the African turquoise killifish bulk brain. **(a)** Scheme of differential chromatin accessibility analysis from bulk ATAC-seq. Explained chromatin accessibility variance was estimated using variancePartition. Differential gene expression analysis was performed with DESeq2. (**b**) Density plots of explained chromatin accessibility variance in bulk GRZ (left) and ZMZ-1001 (right) bulk brain ATAC-seq datasets as a function of aging and sex. Note that aging is the largest contributor to chromatin accessibility variance. (**c**) Strip plot of differentially accessible peaks per strain (FDR < 5%) in aging brains (sex was used as a covariate). Differentially accessible peaks are highlighted while non-significant peaks are shown in gray. Peaks above the midline become more accessible with aging while peaks below the midline become less accessible with aging. (**e**) Pie charts of genomic distributions for differentially accessible ATAC-seq peaks with aging, annotated using HOMER. (**f**) Pie charts of genomic overlap for differentially accessible ATAC-seq peaks with aging, compared to a published H3K4me3 ChIP-seq dataset from young male GRZ brain^22^.

**Extended Data Figure 10:** Functional enrichment analysis for differential chromatin accessibility with aging in the African turquoise killifish bulk brain. (**a**) Scheme of the functional enrichment analysis pipeline for bulk ATAC-seq. Multi-contrast pathway enrichment was performed using mitch to identify gene sets consistently regulated with aging in GRZ and ZMZ-1001 bulk ATAC-seq datasets using Gene Ontology and Reactome gene set definitions. (**b-c**) Top Gene Ontology (b) and Reactome (c) gene sets regulated consistently with brain aging. The top 10 most enriched gene sets are represented, all with FDR < 5%. Heatmaps are colored based on the s statistic of a rank MANOVA test performed with mitch. To disambiguate GO naming redundancy: membrane (i) refers to GO:0016020, and membrane (ii) refers to GO:0016021.

**Extended Data Figure 11:** Analysis of aging microglia molecular and cellular phenotypes in the aging African turquoise killifish brain. (**a**) Multidimensional scaling [MDS] analysis of pseudobulked microglia snRNA-seq and snATAC-seq datasets. Note that samples roughly separate by age along MDS1. (**b**) UCell gene set activity for genes homologous to mouse (i) macrophage polarization genes^79,80^ and (ii) gene signatures of microglia-activation states (*i.e.* terminally inflammatory microglia [TIM]^78^, interferon-response microglia [IRM]^75^, and lipid-droplet-accumulating microglia [LDAM]^75^).

**Extended Data Figure 12:** Machine-learning performance metrics for microglia transcriptional aging classification models. (**a**) Receiver-Operator Curve (ROC) performance evaluation of microglia classification models on training data shows high AUC values (∼0.85) for ElasticNet (eNET) and Random Forest (RF) algorithms. (**b**) AUROC values over the 10 training folds show right distribution consistent with model robustness. (**c**) Balanced accuracy of eNET and RF model on the training (out-of-bag) and testing predictions shows high performance of models (>70% balanced accuracy).

**Extended Data Figure 13:** Analysis of TF regulon activity in snRNA-seq profiling of the aging African turquoise killifish brain. (**a**) Strip plots of differential regulon activity per cell type (decoupleR p < 5%) in aging GRZ (left) and ZMZ-1001 (right) brains with aging (sex was used as a covariate in the input DESeq2 data). Differentially active regulons are highlighted while non-significant regulons are shown in gray. Regulons above the midline are predicted to show increased activity with aging while regulons below the midline decreased activity with aging. (**b**) Heatmaps of DESeq2 log_2_(Fold change in gene expression per week of life) from pseudobulked snRNA-seq data across cell types for the genes encoding the TFs corresponding to the top recurrently regulated decoupleR regulons in GRZ and ZMZ-1001 datasets (also see Figure 6b).

**Extended Data Figure 14:** Conservation of significantly regulated TF regulons in the aging African turquoise killifish brain in mouse and human aging brain. For all analyses in this figure, decoupleR was run across all CollecTRI regulons for mouse and human datasets, and results for the TF regulons homologous to the top African turquoise killifish regulons were plotted as heatmaps. (**a**) Heatmap of predicted age-related changes in activity using decoupleR in publicly available mouse bulk RNA-seq and snRNA-seq brain aging datasets. (**b**) Heatmap of predicted age-related changes in activity using decoupleR in publicly available mouse human bulk RNA-seq (GTEx) and snRNA-seq brain aging datasets. **(c)** Heatmap of predicted age-related changes in activity using decoupleR in publicly available human bulk RNA-seq (GTEx) brain aging datasets. **(d)** Heatmap of predicted age-related changes in activity using decoupleR in publicly available human snRNA-seq brain aging datasets. Note that data was discretized for plotting, similar to our African turquoise killifish analysis: TF regulons with increased (decreased) activity are colored in red (blue), and lighter colors were used to indicate regulons reaching p < 0.1 as an additional line of evidence.

**Extended Data Figure 15:** Analysis of TF footprint accessibility in snATAC-seq profiling of the aging African turquoise killifish brain. (**a**) Scheme of the differential TF footprint accessibility analysis pipeline from snATAC-seq. Analysis at the single-cell level was performed using TF accessibility cell matrices from cellranger-atac, and pseudobulked analyses were performed using TOBIAS. (**b**) Strip plots of differential accessible TF binding motifs per cell type (Seurat findMarkers FDR < 5%) in aging GRZ brain snATAC-seq data with aging. Differentially accessible TF binding motifs are highlighted while non-significant motifs are shown in gray. TF binding motifs above the midline are predicted to show increased accessibility with aging while motifs below the midline decreased activity with accessibility. (**c**) Strip plots of differential TF footprints accessibility per cell type (Pseudobulk TOBIAS FDR < 5%) in aging GRZ brain snATAC-seq data with aging. Differentially accessible TF footprints are highlighted while non-significant footprints are shown in gray. Footprints above the midline are predicted to show increased accessibility with aging while footprints below the midline decreased activity with accessibility. (**d**) TOBIAS differential footprint accessibility analysis for JASPAR glucocorticoid receptor motif NR3C1_MA0113.3 matrix, showing increased accessibility in all assayed cell types except ependymal cells.

**Extended Data Figure 16:** Analysis of TF motif enrichment and footprint accessibility in bulk ATAC-seq of the aging African turquoise killifish brain. (**a**) Scheme of the differential TF motif enrichment and footprint accessibility analysis pipeline from bulk ATAC-seq. Motif enrichment analysis on differentially accessible peaks was performed using HOMER, while differential footprint accessibility analysis was performed using TOBIAS. (**b**) Pairwise scatterplots across sexes in each strain to evaluate change in TF footprint accessibility with aging in GRZ (left) and ZMZ-1001 (right) datasets. Note the position of the Nr3c1 footprint, highlighted in red, as robustly showing increased accessibility with aging regardless of the comparison groups. (**c**) Bubble plot of the top 10 differentially accessible footprints in bulk ATAC-seq from aging brains according to TOBIAS (5 with increased accessibility, 5 with decreased accessibility with aging). TOBIAS footprint scores and FDR are reported. Note significant recurrent increased in Nr3c1 footprint accessibility across comparisons in bold.

**Extended Data Figure 17:** Impact of mid-life-initiated mifepristone treatment on African turquoise killifish biometrics. (**a**) Boxplot of length female (left) and male (right) African turquoise killifish from the young control, old control and old mifepristone treated fish. (**b**) Boxplot of weight female (left) and male (right) African turquoise killifish from the young control, old control and old mifepristone treated fish. All fish from the GRZ strain. Data from 3 independent cohorts of animals. P-values in non-parametric Wilcoxon rank-sum tests are reported.

**Extended Data Figure 18:** Impact of mid-life-initiated mifepristone treatment on the aging African turquoise killifish brain transcriptome. (**a**) Strip plots of differentially expressed genes and TEs in bulk RNA-seq of turquoise killifish brains in response to aging and to mid-life-initiated mifepristone treatment in our female and male transcriptomic cohorts. Differentially expressed transcripts are highlighted while non-significant genes are shown in gray. Transcripts above the midline are upregulated while transcripts below the midline downregulated. (**b**) Overlap analysis of genes and TEs upregulated and downregulated with aging vs. downregulated and upregulated in response to mid-life-initiated mifepristone treatment across our female and male transcriptomic cohorts. Genes passing FDR < 5% were compared. Significance of overlaps according to Fisher’s exact test are reported below Venn diagrams. Note significant overlaps, consistent with a partial (but significant) reversal of age-related transcriptional changes in the aged brain after mifespristone exposure. (**c**) Scatterplot of gene and TE expression log_2_(Fold changes) with aging vs. in response to mid-life-initiated mifepristone treatment across our female and male transcriptomic cohorts. Inset are the Spearman Rho values and their significance. Significant anti-correlation is consistent with partial transcriptome-wide reversal of age-related transcriptional changes in the aged brain after mifespristone exposure. (**d**) Scatterplot of gene and TE expression log_2_(Fold changes) with aging and in response to mid-life-initiated mifepristone treatment, comparing across our 2 independent cohorts. Inset are the Spearman Rho values and their significance. Significant correlation is consistent with good agreement between results obtained across cohorts/sexes. (**e**) Overlap analysis of genes and TEs upregulated and downregulated with aging vs. downregulated and upregulated in response to mid-life-initiated mifepristone treatment in the meta-analysis of our female and male transcriptomic cohorts. Genes passing FDR < 5% for the inverse normal combination method were compared. Significance of overlaps according to Fisher’s exact test are reported below Venn diagrams. (**f-g**) Dotplots of top enriched Gene Ontology (GO) and Reactome gene sets by overrepresentation analysis for genes upregulated with aging and downregulated with mifepristone (**f**) and genes downregulated with aging and upregulated with mifepristone (**g**). The top 5 most significantly enriched gene sets (or as many gene sets that reached significance) are plotted.

**Extended Data Figure 19:** Impact of mid-life-initiated mifepristone treatment on microglia abundance in the African turquoise killifish brain. (**a**) Boxplot of VST-normalized DEseq2 count expression of microglia marker gene *apoeb* in bulk brain RNA-seq datasets. Reported DESeq2 FDRs showing significance of pairwise comparisons. (b) CSCDRNA deconvolution predicted microglia cell proportions in the brains of young control fish, old control fish and old mifespristone-treated fish. P-values in non-parametric Wilcoxon rank-sum tests are reported.

## Inventory of Extended Data Tables

**Extended Data Table 1: Information and metrics for snRNA-seq, snATAC-seq and bulk ATAC-seq libraries.**

(**a**) Cell Ranger snRNA-seq library statistics. (**b**) Cell Ranger snATAC-seq library statistics. (**c**) Bulk ATAC-seq bowtie2 mapping statistics. (**d**) Conversion table for NCBI African turquoise killifish gene names to “human readable” equivalents used in this study throughout text and figures.

**Extended Data Table 2: Differential gene-expression analysis with aging for snRNA-seq across cell types**.

Results of the Muscat/DESeq2 analyses with one tab for each cell type in each strain. (*Note that sex was used as a model covariate for these analyses*).

**Extended Data Table 3: Differential chromatin accessibility analysis with aging for snATAC-seq across cell types**.

Results of the Muscat/DESeq2 analyses with one tab for each cell type in GRZ strain. (*Note that sex was used as a model covariate for these analyses*).

**Extended Data Table 4: Differential chromatin accessibility analysis with aging for bulk ATAC-seq across cell types**.

(a) Annotated DESeq2 results for bulk ATAC-seq (age-regulation of chromatin accessibility) aging **GRZ** brains. (**b**) Annotated DESeq2 results for bulk ATAC-seq (age-regulation of chromatin accessibility) aging **ZMZ-1001** brains. (*Note that sex was used as a model covariate for these analyses*).

**Extended Data Table 5: Mitch multi-contrast enrichment results in aging turquoise killifish brains for Gene Ontology gene sets (FDR < 5%).**

Results of Mitch analyses for GO (BP, MF and CC) gene sets, with one tab for each cell type.

**Extended Data Table 6: Mitch multi-contrast enrichment results in aging turquoise killifish brains for REACTOME gene sets (FDR < 5%).**

Results of Mitch analyses for Reactome gene sets, with one tab for each cell type.

**Extended Data Table 7: Parameters related to machine-learning models of microglia transcriptional aging**.

(a) Results of Dunn Test with Holm multiple testing correction for predicted class probabilities across microglia from different age groups in RF and ENET models. (**b**) Machine-learning variable importance parsing derived from random forest (RF) and elastic net (ENET) trained models.

**Extended Data Table 8: Results of differential predicted TF activity analyses across ‘omic’ layers**.

(a) Differentially regulated decoupleR TF regulons in aging GRZ brains per cell type (FDR < 10%). (**b**) Differentially regulated decoupleR TF regulons in aging ZMZ-1001 brains per cell type (FDR < 10%). (c) Differential motif accessibility analysis in snATAC-seq dataset using cellranger-atac in aging GRZ brains per cell type (FDR<5%). (**d**) Differential motif accessibility analysis in snATAC-seq dataset using TOBIAS in aging GRZ brains per cell type (FDR<5%). (**e**) HOMER motif enrichment in bulk ATAC-seq differentially accessible peaks in aging GRZ brains (FDR < 5%). (**f**) HOMER motif enrichment in bulk ATAC-seq differentially accessible peaks in aging ZMZ-1001 brains (FDR < 5%). (**g**) Differential motif accessibility analysis in bulk ATAC-seq dataset using TOBIAS in aging GRZ brains (FDR<5%). (**h**) Differential motif accessibility analysis in bulk ATAC-seq dataset using TOBIAS in aging ZMZ-1001 brains (Young vs. Old; FDR<5%). (**i**) Differential motif accessibility analysis in bulk ATAC-seq dataset using TOBIAS in aging ZMZ-1001 brains (Young vs. Geriatric; FDR<5%).

**Extended Data Table 9: Analyses of the impact of mid-life-initiated mifepristone treatment on brain gene expression**. (**a**) DESeq2 results for differential gene expression in <u>female</u> GRZ brains with aging (6 vs. 16 weeks; cohort 1). (**b**) DESeq2 results for differential gene expression in <u>female</u> GRZ brains with mifepristone treatment (16 weeks; cohort 1). (**c**) DESeq2 results for differential gene expression in <u>male</u> GRZ brains with aging (6 vs. 16 weeks; cohort 2). (**d**) DESeq2 results for differential gene expression in <u>male</u> GRZ brains with mifepristone treatment (16 weeks; cohort 2). (**e**) metaRNAseq results of DESeq2 results for differential gene expression in <u>female</u> and <u>male</u> GRZ brains with aging (6 vs. 16 weeks). (**f**) metaRNAseq results of DESeq2 results for differential gene expression in <u>female</u> and <u>male</u> GRZ brains with mifepristone treatment (16 weeks). (**g**) mitch multi-contrast enrichment results in aging and mifepristone-treated GRZ brains for Gene Ontology gene sets (FDR < 5%). (**h**) mitch multi-contrast enrichment results in aging and mifepristone-treated GRZ brains for REACTOME gene sets (FDR < 5%). (**i**) Overrepresentation analysis of GO/REACTOME gene sets for genes upregulated with aging and downregulated in response to mifepristone (FDR < 5%). (**j**) Overrepresentation analysis of GO/REACTOME gene sets for genes downregulated with aging and upregulated in response to mifepristone (FDR < 5%).

**Extended Data Table 10: Accession numbers of publicly available datasets reanalyzed in this study**.

(a) Whole Genome Shotgun sequencing datasets used for turquoise killifish strain genetic diversity analysis. (**b**) Publicly available bulk and snRNA-seq datasets on mouse and human brain aging used in conservation analyses.

