## Extended Data Figures for "A multi-omic atlas in the African turquoise killifish reveals increased glucocorticoid signaling as a hallmark of brain aging"

Extended Data Figure 1

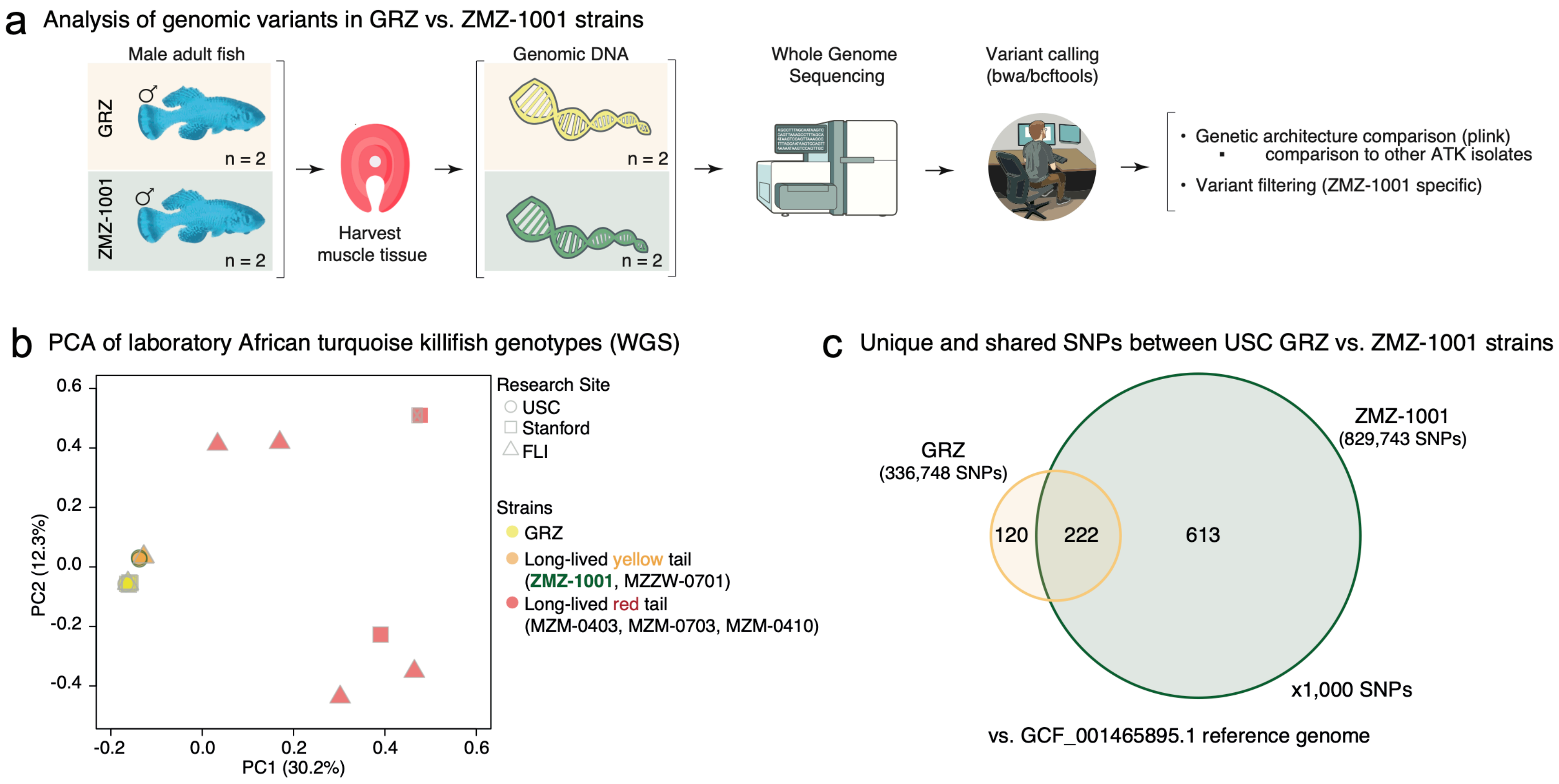

Extended Data Figure 2

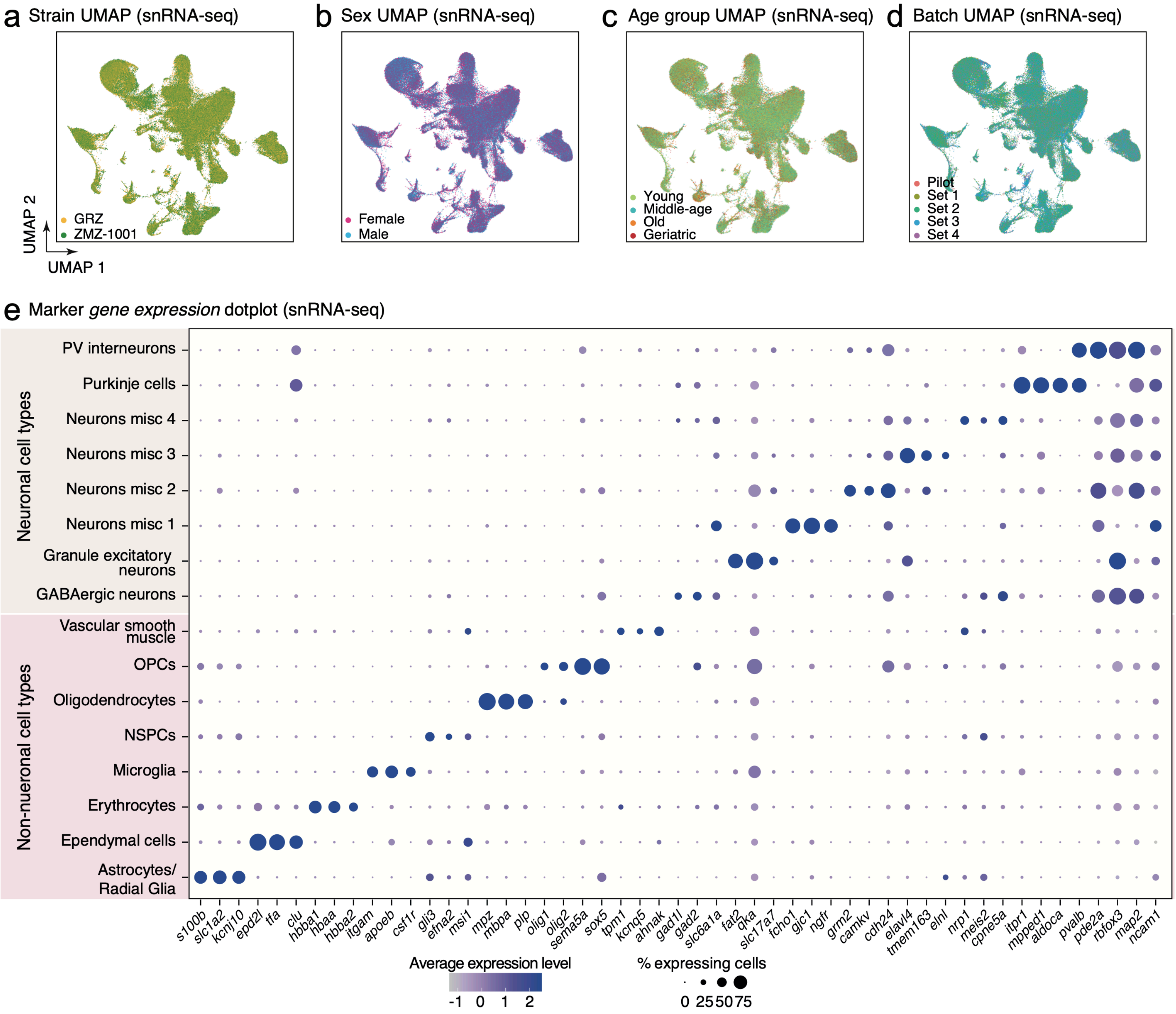

Extended Data Figure 3

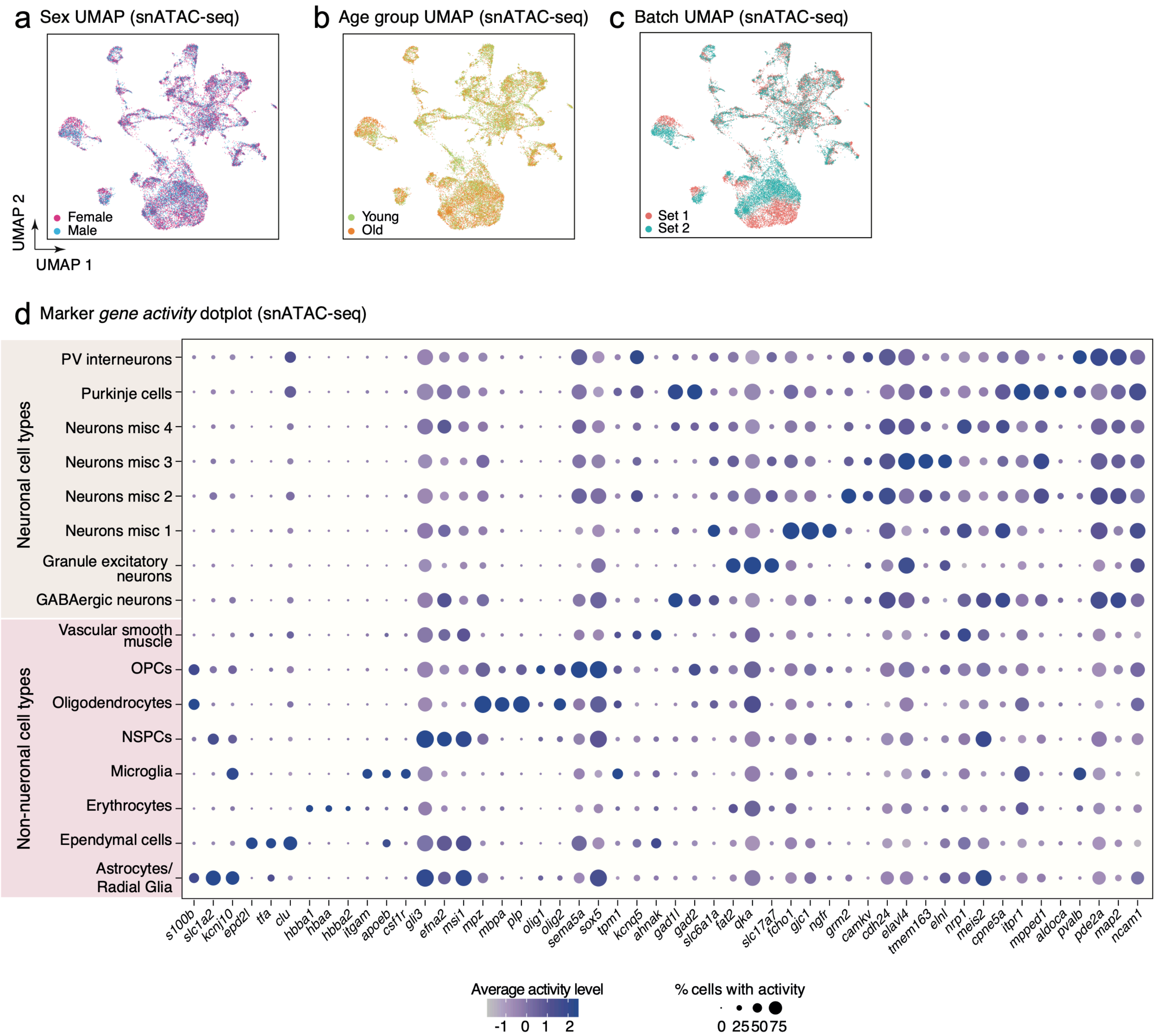

Extended Data Figure 4

**a** Augur AUC scores for separation as a function of age group by cell type

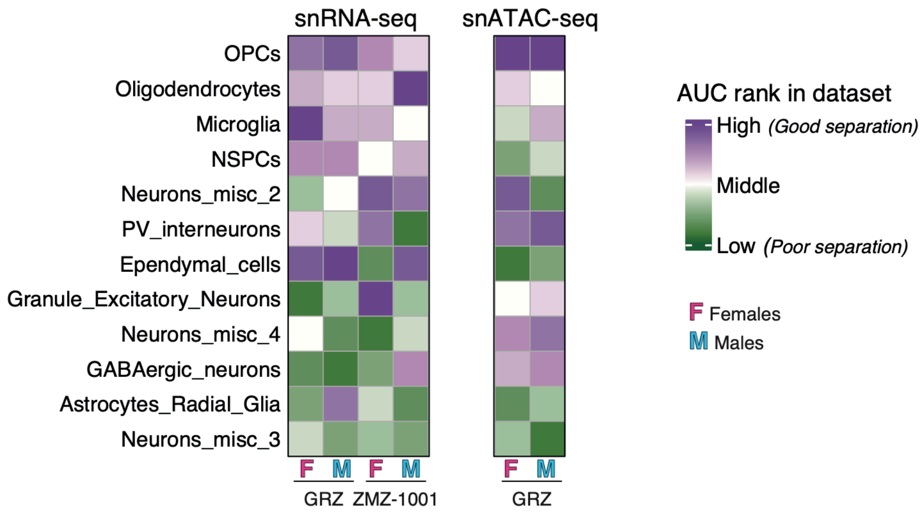

### Extended Data Figure 5

**a** Representative images of RNAscope labeling in ZMZ-1001 brain (RNAscope)

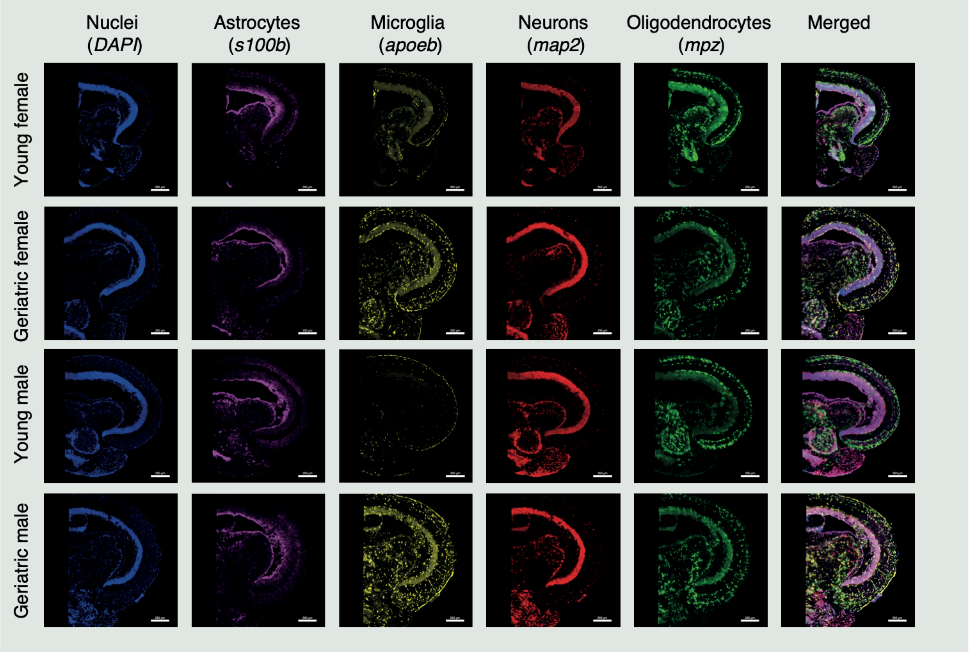

Extended Data Figure 6

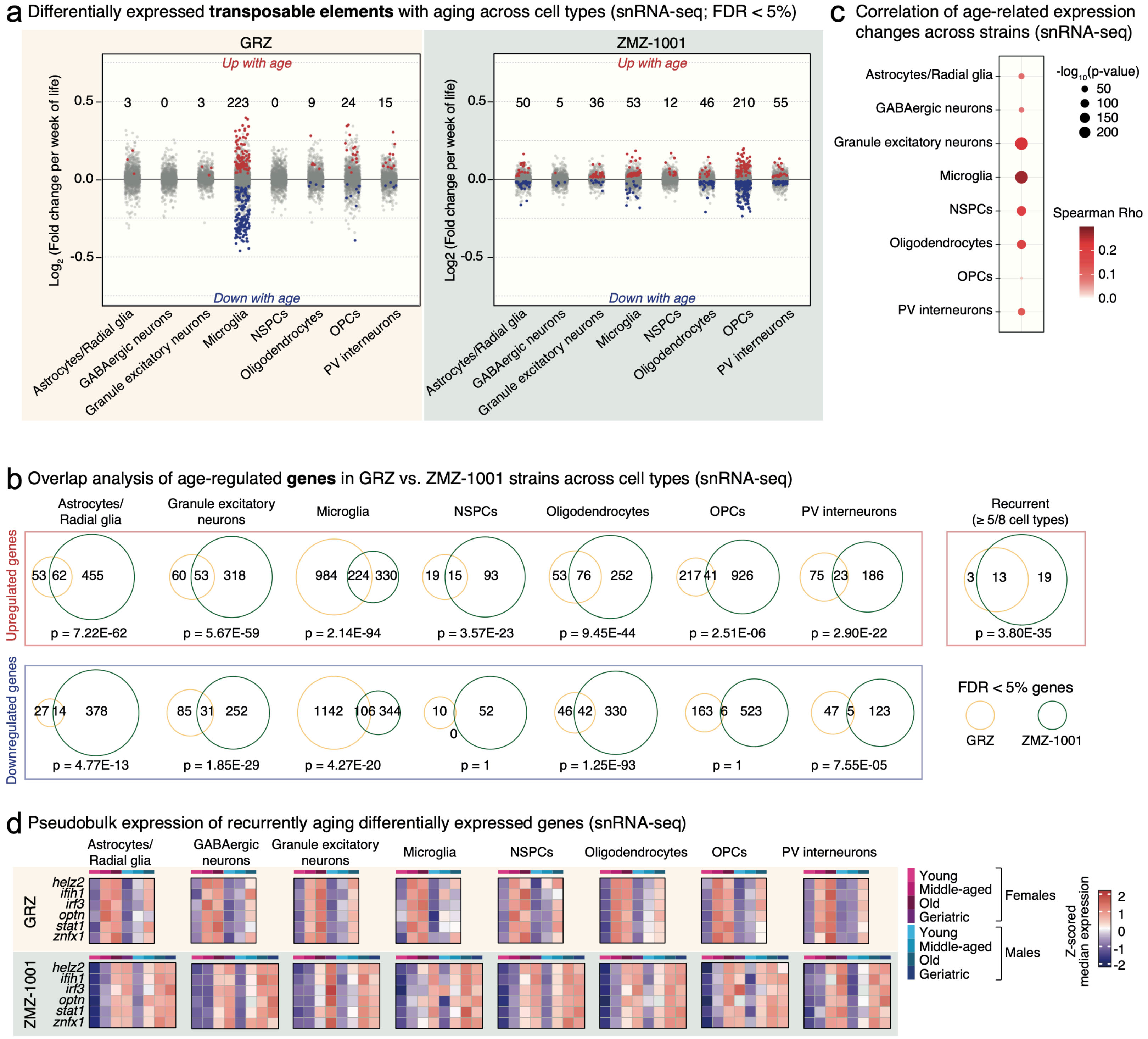

#### Extended Data Figure 7

**a** Representative images of differentially expressed gene validation set labeling (RNAscope **Set 1**; ZMZ-1001 GF animal example)

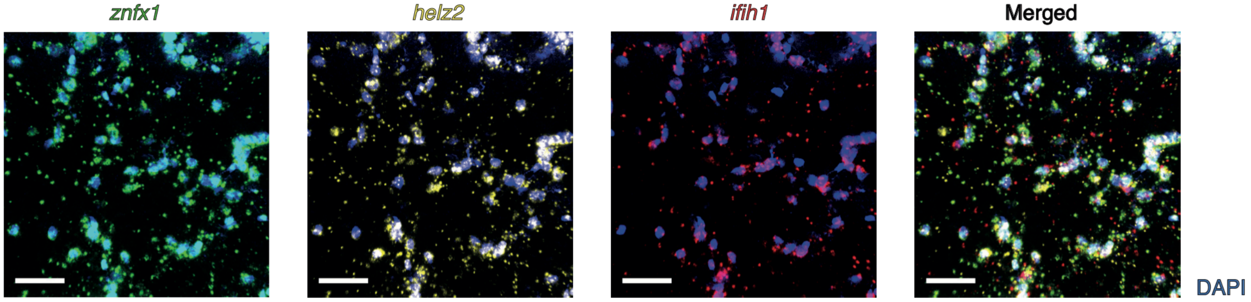


**b** Representative images of differentially expressed gene validation set labeling (RNAscope **Set 2**; ZMZ-1001 GF animal example)

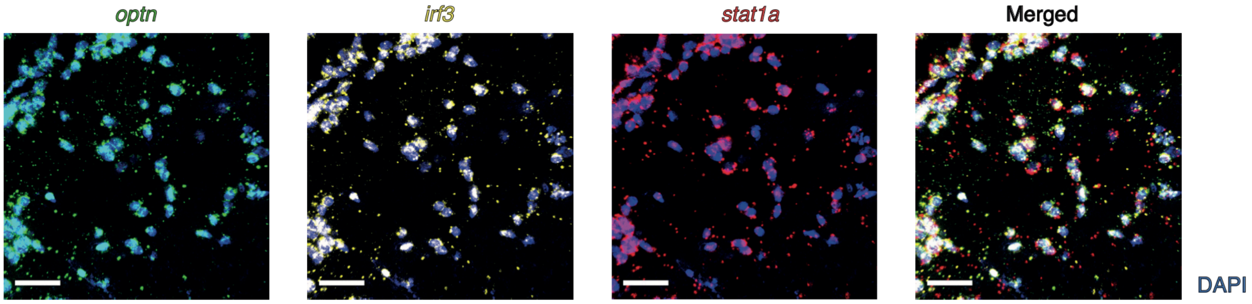

Extended Data Figure 8

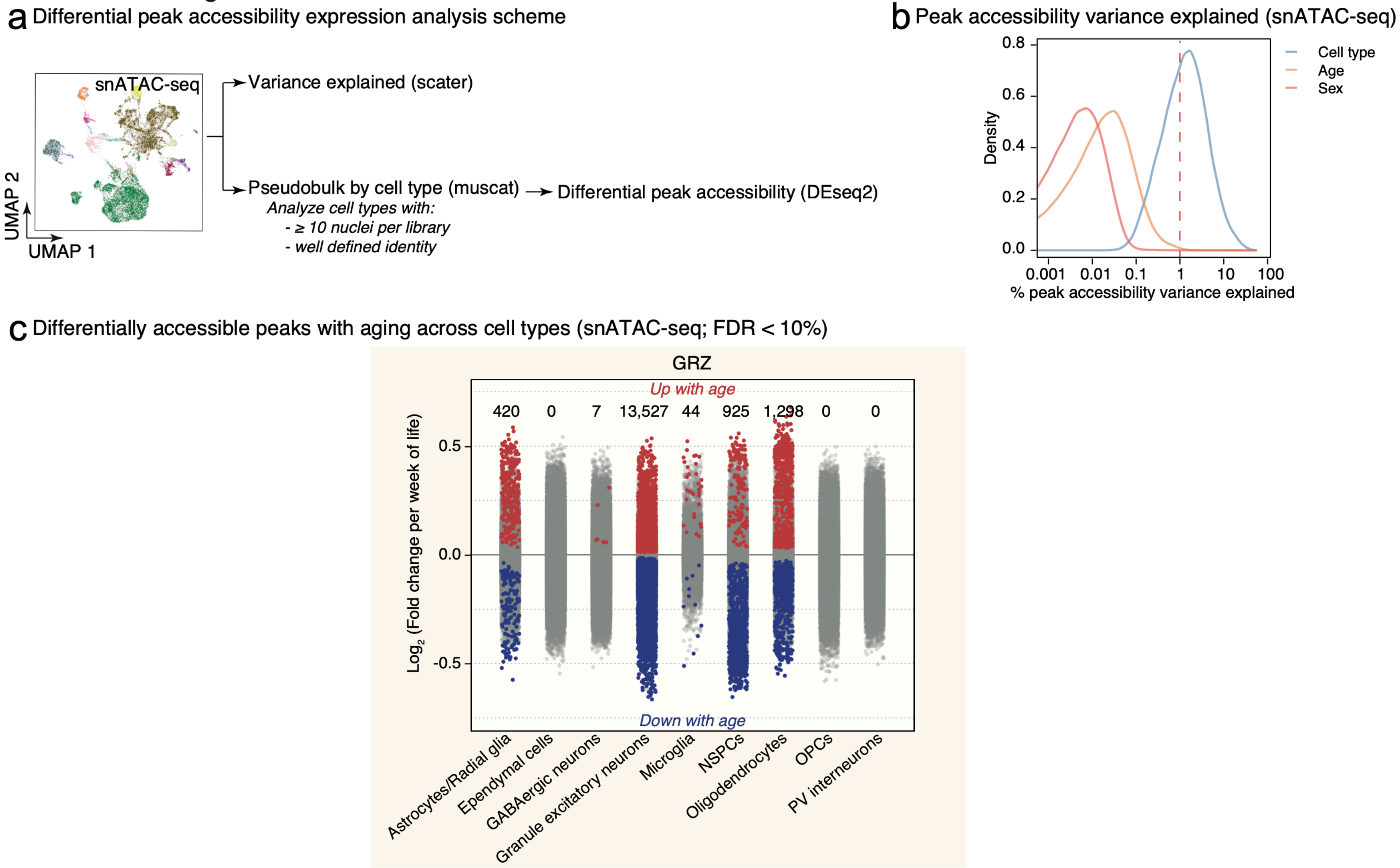

Extended Data Figure 9

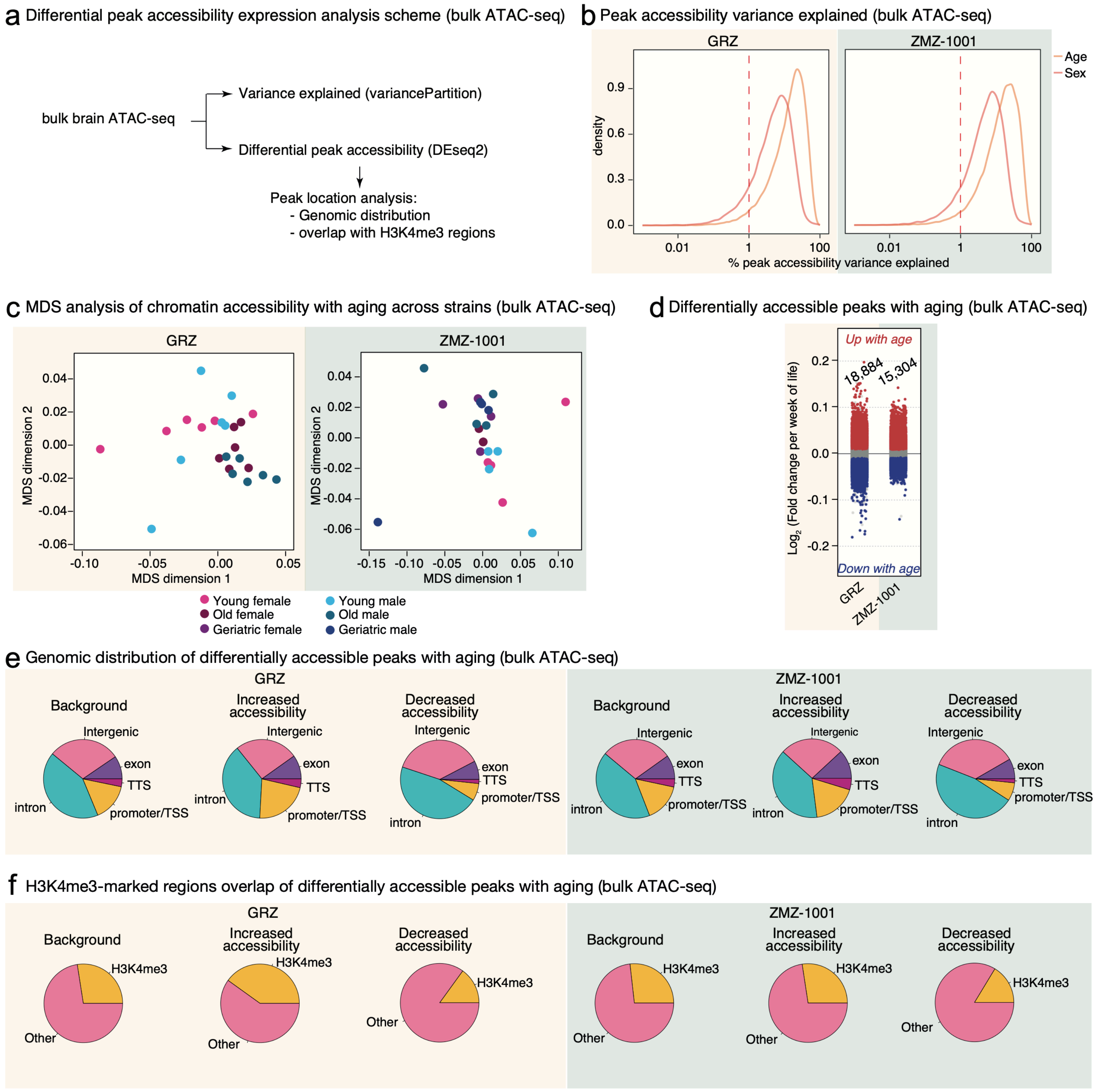

### Extended Data Figure 10

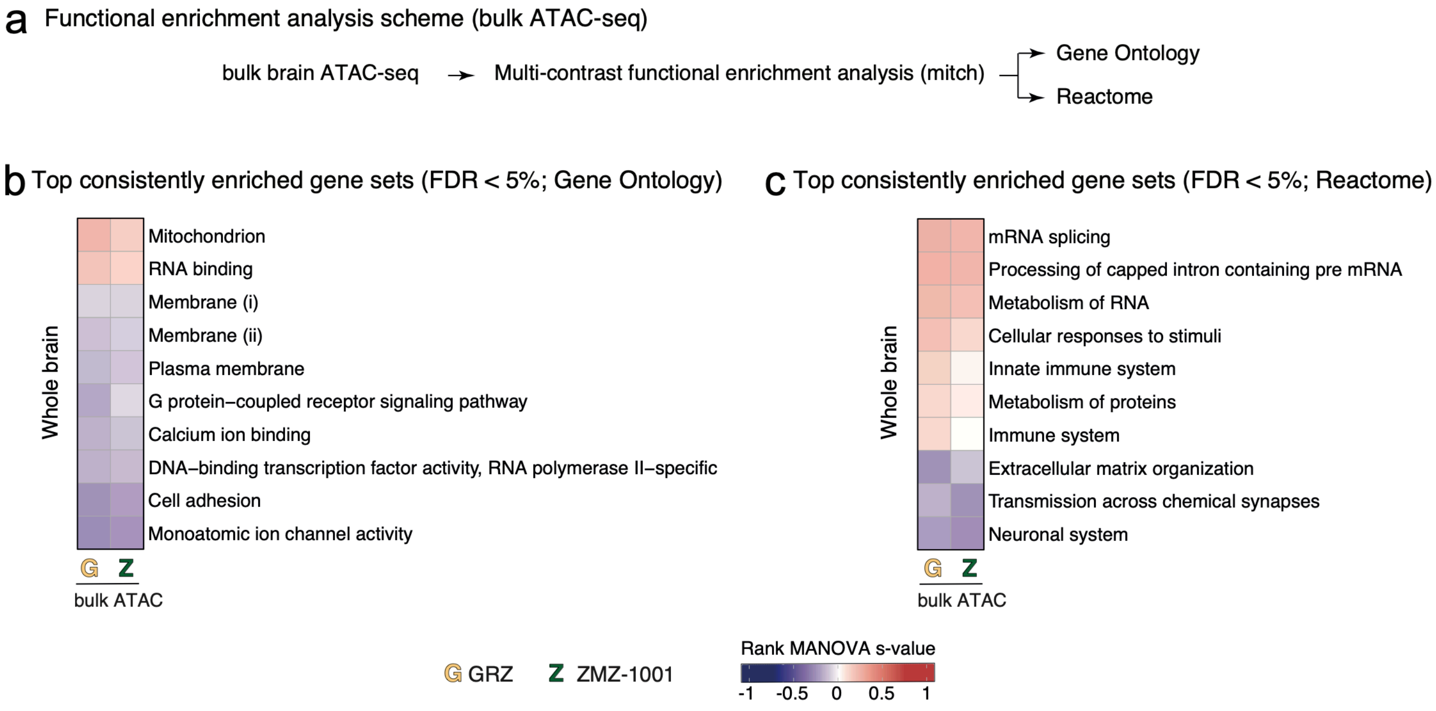

### Extended Data Figure 11

**a** Multidimensional scaling analysis of pseudobulked **microglia** data

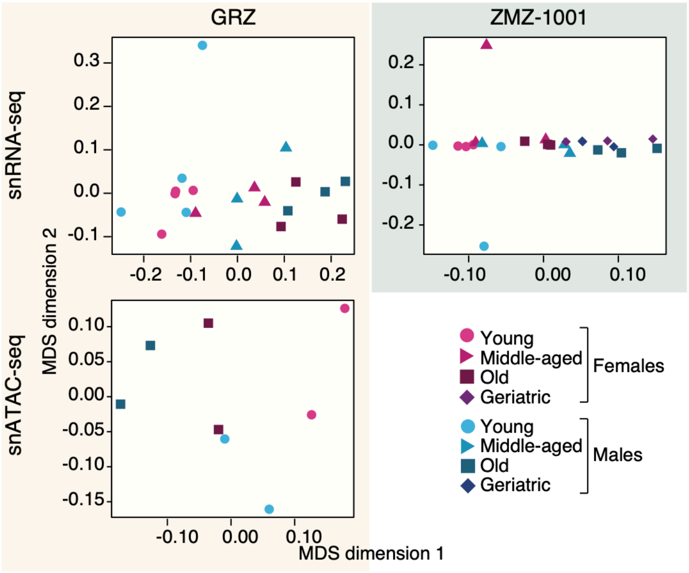


**b** Activity of **microglia**-aging relevant gene sets (snRNA-seq)

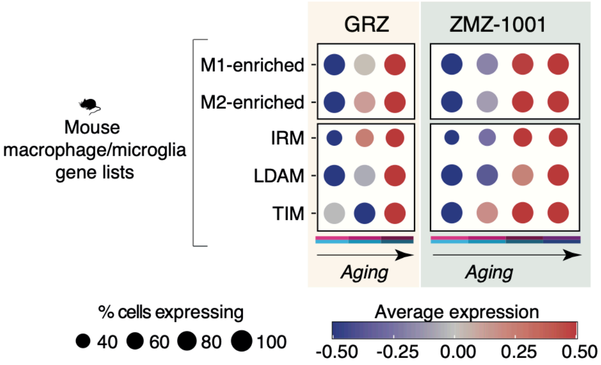

### Extended Data Figure 12

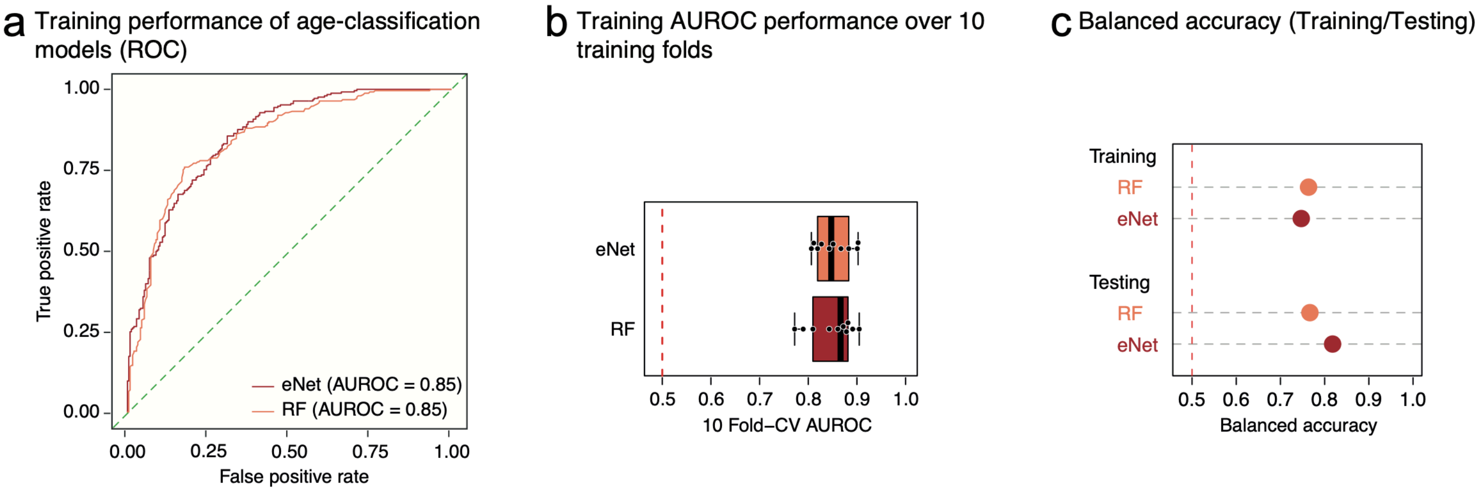

Extended Data Figure 13

**a** Differentially active **TF regulons** with aging across cell types (snRNA-seq; p-value < 5%)

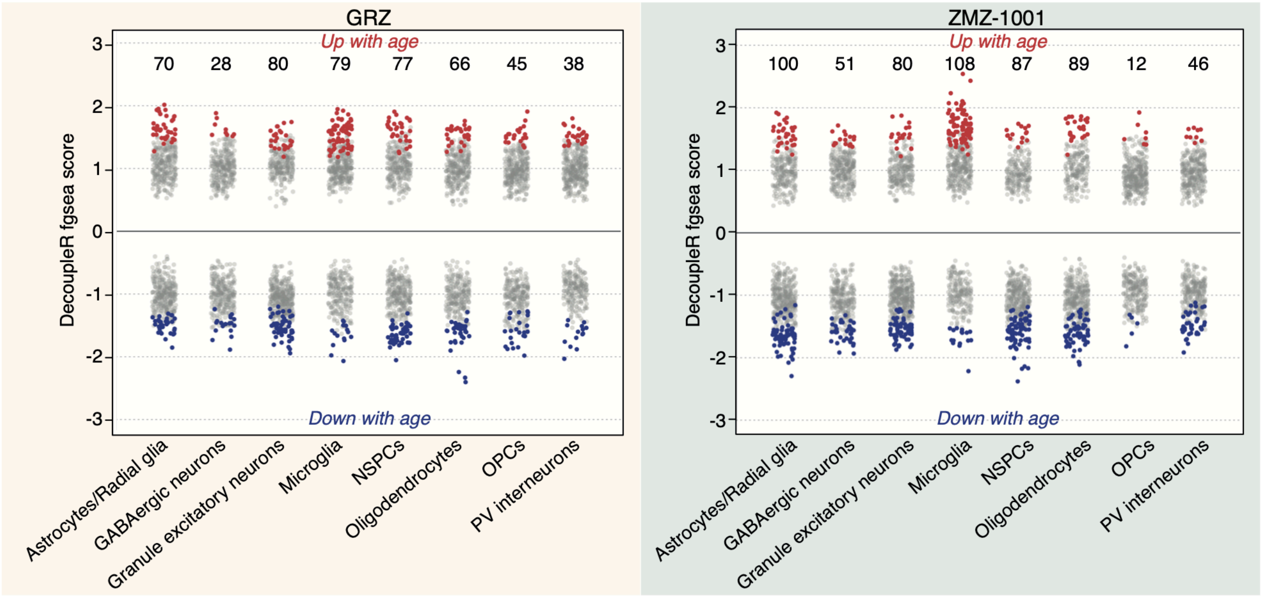


**b** Top **TF candidate expression** change with aging (snRNA-seq)

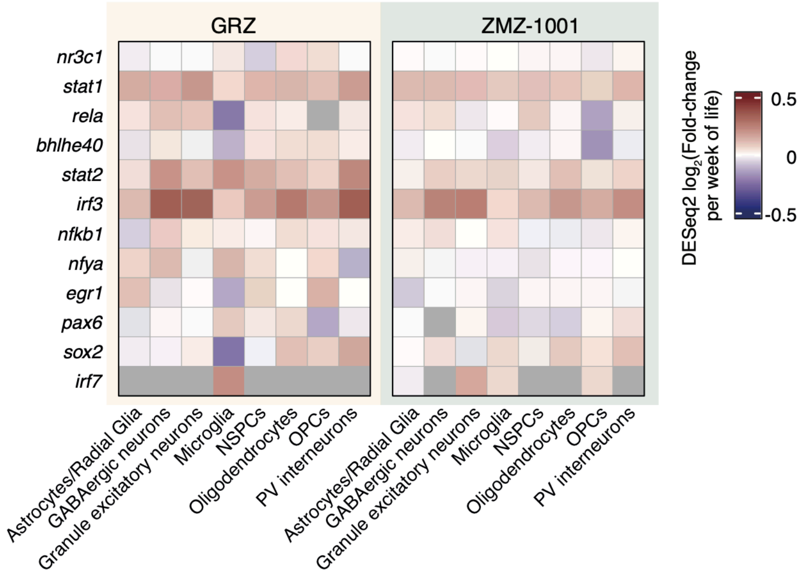

Extended Data Figure 14

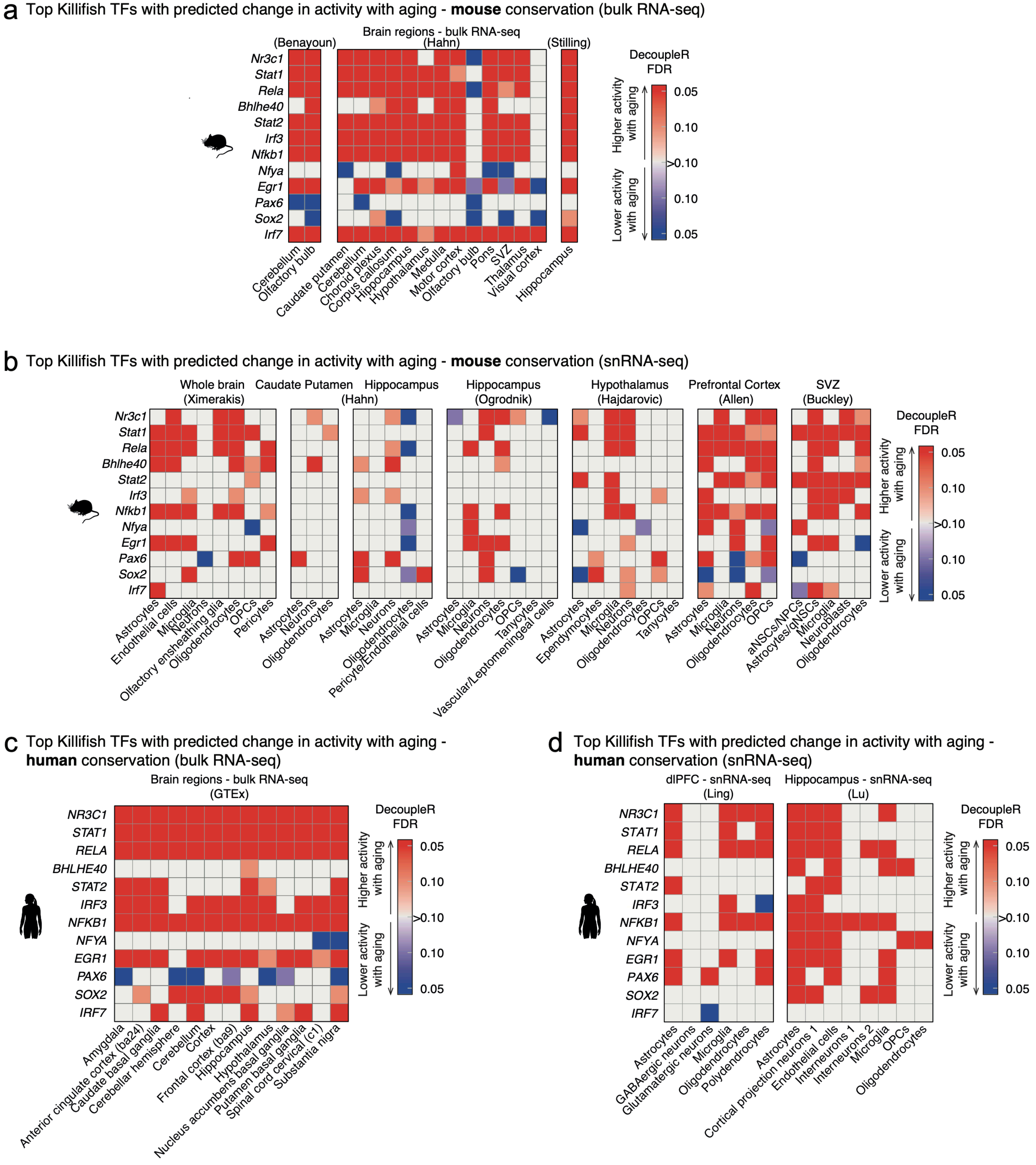

Extended Data Figure 15

a Differential footprint accessibility analysis scheme (snATAC-seq)

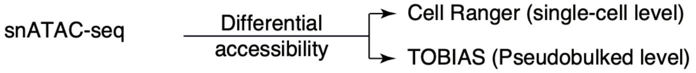


b Differentially accessible footprints with aging across cell types (**cellranger-atac** and **findMarkers**; FDR <5%)

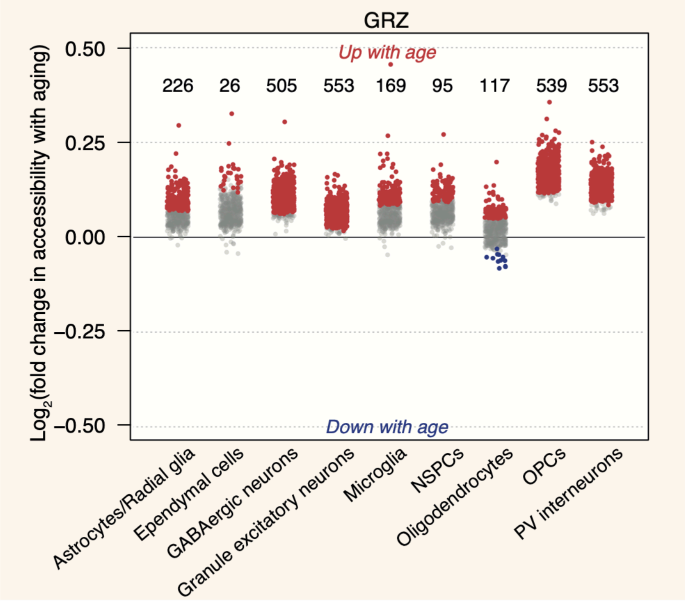


c Differentially accessible footprints with aging across cell types (**TOBIAS**; FDR <5%)

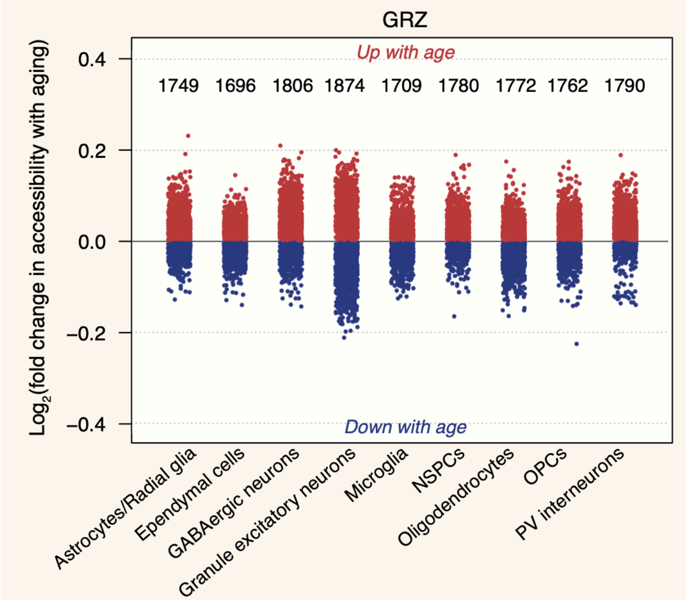


d Differential accessibility analysis for NR3C1_MA0113.3 footprints with aging across cell types (TOBIAS)

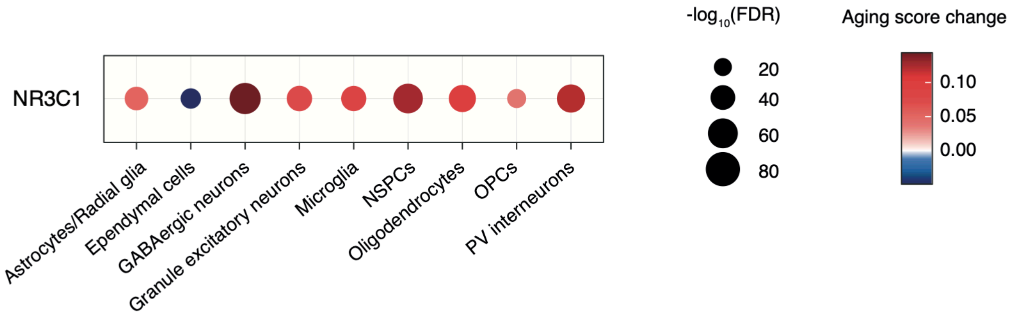

Extended Data Figure 16

a Differential footprint accessibility analysis scheme (bulk ATAC-seq)

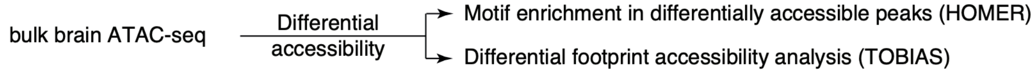


b TOBIAS differential footprint accessibility scores (bulk ATAC-seq)

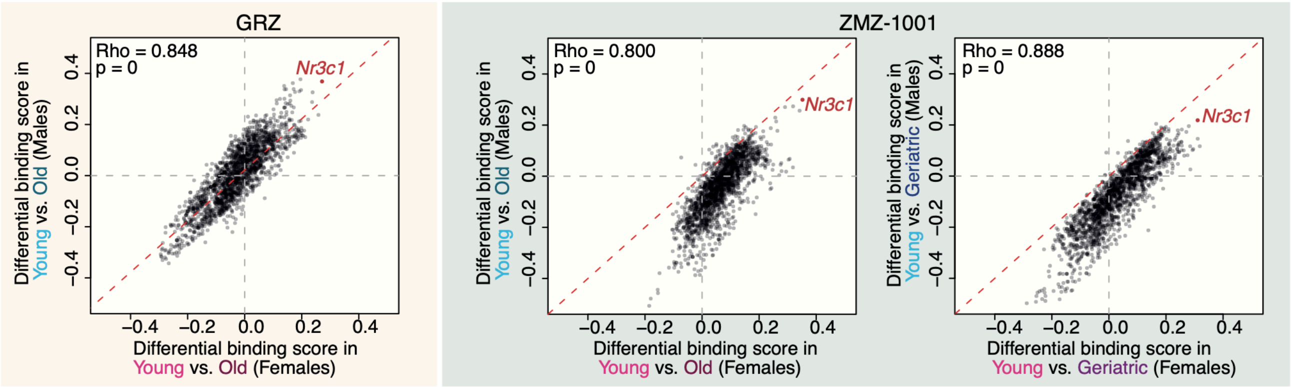


c Top 5 consistently differentially accessible footprints with increased/decreased accessibility with aging (FDR < 5%; TOBIAS) (bulk ATAC-seq)

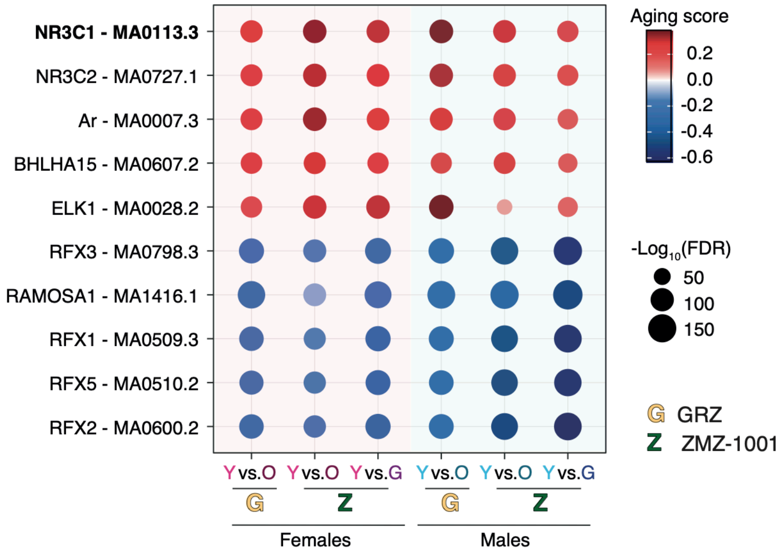

Extended Data Figure 17

a Effect of mid-life initiated mifepristone treatment on fish length

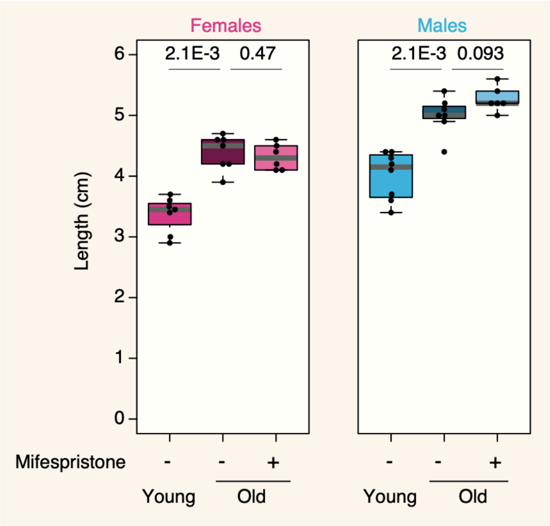


b Effect of mid-life initiated mifepristone treatment on fish weight

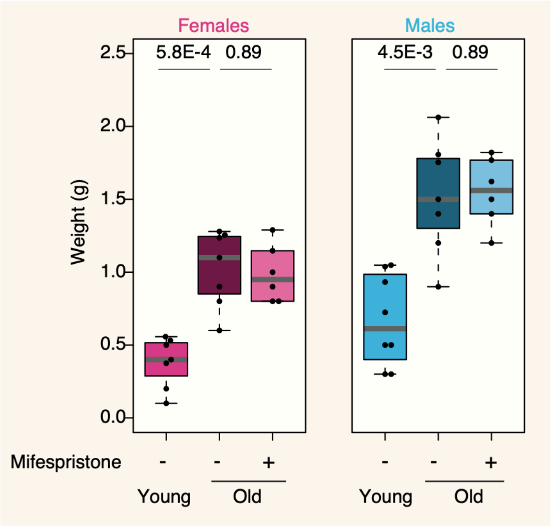

Extended Data Figure 18

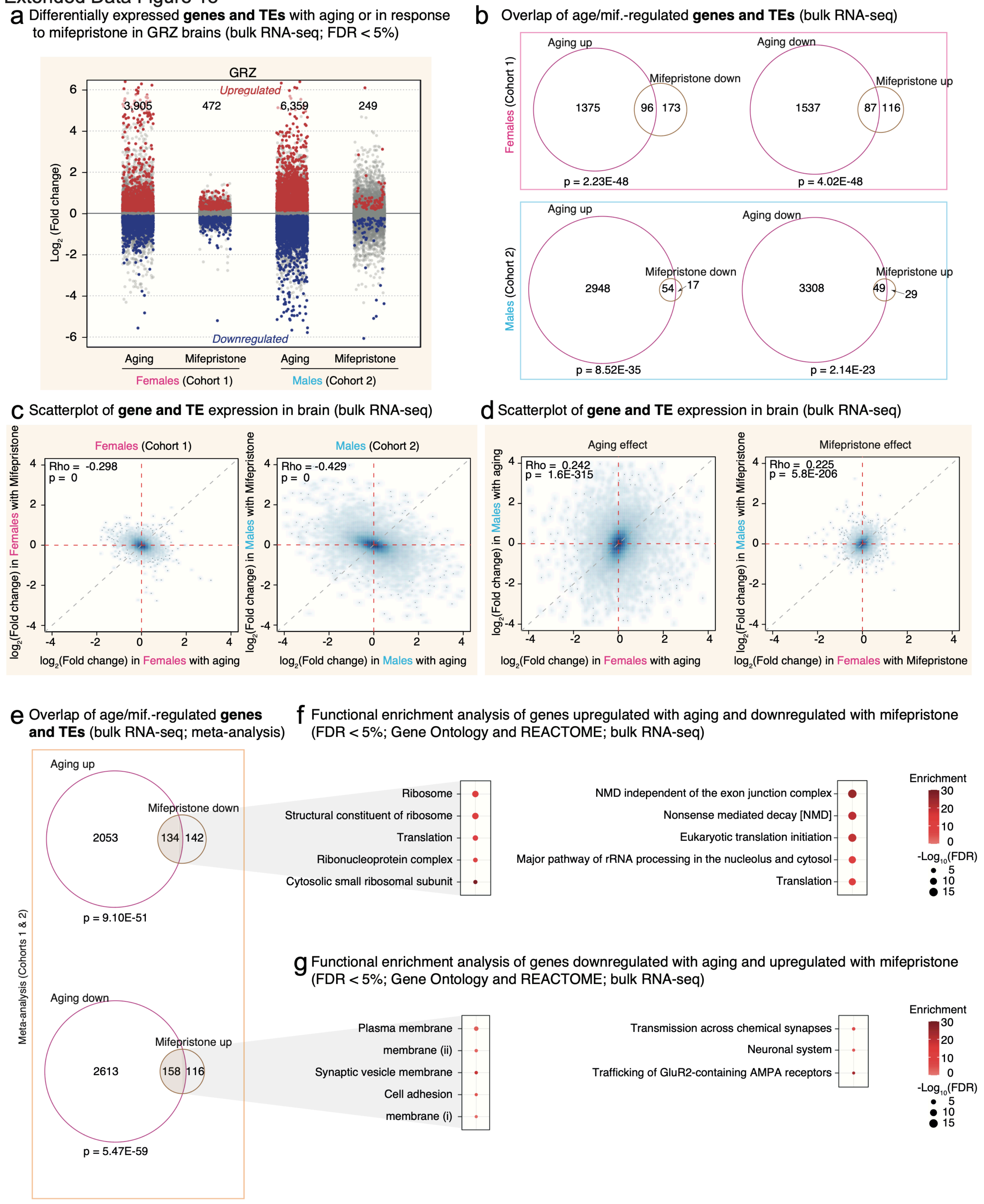

Extended Data Figure 19

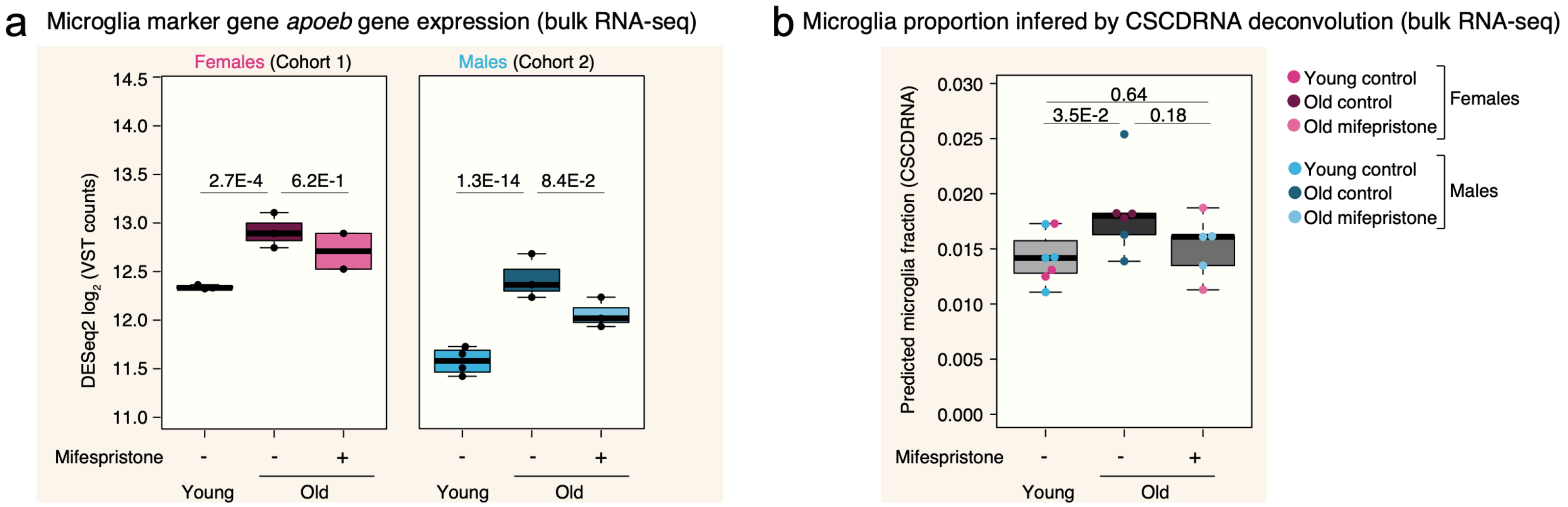
